# Sediment substrate size influences fish diversity in tributary mouth areas in impounded boreal rivers in Sweden

**DOI:** 10.1101/2022.06.30.498239

**Authors:** J. Näslund, R. Bowes, L. Sandin, E. Bergman, L. Greenberg

**Affiliations:** Department of Aquatic Resources, Institute of Freshwater Research, Swedish University of Agricultural Sciences, Drottningholm, Sweden; Department of Environmental and Life Sciences, River Ecology and Management, Karlstad University, Karlstad, Sweden; Biological Sciences Department, Emporia State University, Emporia, KS, USA; Section for Nature-based solutions and Aquatic Ecology, NIVA - Norwegian Institute for Water Research, Oslo, Norway

**Keywords:** Aggradation, Boreal rivers, Fish biodiversity, River morphology, River sediment size, Tributary confluence

## Abstract

Large boreal rivers in Sweden are generally impounded by hydropower dams and a large proportion of their main stem shallow flowing habitats have been lost. Tributaries often contain relatively undisturbed habitats and could be important for the conservation of species diversity. Tributary mouth areas could be biodiversity hot-spots, due to the vicinity to the main stem and favorable environmental conditions. In this study, we investigate whether tributary mouth areas in two impounded boreal rivers (Ume- and Lule River) could be regarded as biodiversity hot spots for fish. Based on electrofishing in 20 tributary mouths, we found that overall fish diversity is generally low. The highest species richness and diversity was found in mouth areas dominated by intermediate substrate sizes (gravel – cobble). Few, if any, species were found in association with fine sediment substrates (smaller than sand). The tributary mouth areas had similar species richness and diversity as areas in the tributaries located 1-km upstream of the mouth, but the fish community composition often differed between these sites. Management action favoring fish diversity in the tributary mouth areas could include protection or rehabilitation of areas dominated by medium sized substrate and reduction of erosion and transport of fine sediments in the tributaries. Overall, we find no support for tributary mouths being hot-spots for fish biodiversity and while some patterns in diversity gives hints on suitable management action, it is important to further understand impacts in tributaries and their mouths especially in relation to temporal dynamics of the fish community.

## INTRODUCTION

Large rivers are important ecosystems for aquatic biodiversity, typically housing a higher fish biodiversity than smaller rivers, with some species being adapted specifically for habitats characteristic of large rivers (Jackson et al. 2001; Oberdorff et al. 2011; Miranda et al. 2019). The biodiversity of many of these large ecosystems has been negatively impacted during the last century as they have been heavily exploited for many purposes, including energy production (Grill et al. 2019). While damming of main stem rivers can secure the production of affordable energy for society (Rex et al. 2014; Schäfer 2021), damming also disrupts longitudinal connectivity and alters hydrological, hydrogeomorphological, and thermal conditions in river systems (Baxter 1977; Ligon et al. 1995; Vörösmarty et al. 2010; Anderson et al. 2015). The result is spatially and seasonally homogenized, slow-flowing river sections between dams, with severely disrupted connection among them, which reduces their overall biodiversity (Poff et al. 2007; Malm Renöfält et al. 2010).

The northern European large rivers discharging into the Baltic Sea are today generally exploited for large-scale hydropower production, with only a few remaining in a completely or near free-flowing state (Dynesius & Nilsson 1994; Grill et al. 2019). In Sweden, hydropower development in these Baltic rivers was initiated in the early 1900’s, without much regard to the ecology of the river ecosystems, and today constitute the most important source of hydropower in the country (Ödmann et al. 1982; Schäfer 2021). With national and international legal requirements for sustainable hydropower (GOS 2020) and riverine ecosystem functioning (EC 2000), the pressure to restore riverine biodiversity and processes is high. Sweden is a net-exporter of energy, with hydropower making up an important proportion of the energy production, and sources of renewable energy are increasingly important in light of climate change (SEA 2021). Hence, while complete restoration to reference conditions is unfeasible in these heavily modified rivers, rehabilitation measures and mitigation of continued biodiversity loss are still required to secure good ecological potential (EC 2000, 2020; GOS 2020). Such management actions, however, requires much information about the current ecological state of different areas, so that measures can be optimally designed and directed to areas where they are most likely to have the strongest positive effects (Jansson et al. 2007; Widén et al. 2016).

Tributaries to large impounded rivers often have a lower degree of habitat alteration and more natural sediment transport, flow dynamics, and temperature regimes, as compared to the main stem, given that they themselves are not impounded (Rice et al. 2008; Ziv et al. 2012; Pracheil et al. 2013). Protection of unaltered tributaries and restoration of ecologically degraded tributaries could thereby retain or improve the remaining ecological values of impounded large rivers. Focus on this potential source of biodiversity has mainly been proposed for large tributaries in very large river systems (Ziv et al. 2012; Pracheil et al. 2013; Dunn & Paukert 2021), where network dispersal (extensive upstream dispersal in tributaries) is more likely than in small tributaries (Grenoulliet et al. 2004). Nevertheless, similar effects may also be achievable in smaller systems where the main stem-tributary movements of species mainly relates to confluence exchange (localized movements near confluences), if conditions are favorable (Rice et al. 2008; Thornbrugh & Gido 2010; Laub et al. 2018). The confluence areas of tributaries (mouth) and the main stem have previously been described as important for habitat heterogeneity and biodiversity in the main stem, due to e.g. aggradation, concentrated nutrient flow, favorable thermal and chemical conditions, and environmental features allowing prey species to shelter from large-sized predators (Power & Dietrich 2002; Wipfli & Gregovich 2002; Kiffney et al. 2006; Rice et al. 2006). Tributary confluences can indeed constitute significant biological ‘hot spots’ in riverine ecosystems (Power & Dietrich 2002; Benda et al. 2004; Kiffney et al. 2006). Previous studies have shown that benthic invertebrate diversity is high at tributary confluences in reaches below impoundments in the main stem (e.g. Greenwood et al. 1999; Vinson 2001; but see Milner et al. 2019). Studies on fish also suggest that species richness can be higher in tributary segments near main stem confluences (Thornbrugh & Gido 2009; Miranda et al. 2019). These patterns make tributary mouths possible focal areas for 1) conservative management plans when intact, 2) restorative actions when degraded, or 3) ecological compensation measures when the main stem is degraded and there is still potential to increase biodiversity (Allan & Castillo 2007; Rice et al. 2008; Erkinaro et al. 2017; Sandin et al. 2017; Miranda et al. 2019).

The tributaries themselves (i.e. sections upstream from the mouth) often have lower species richness than the main stem (Czeglédi et al. 2015; Miranda et al. 2019), but can nevertheless be important for biodiversity in the river network. For instance, tributaries can be highly heterogeneous in terms of flow and environmental features, offering a variety of habitats (Jackson et al. 2001; Wohl 2017). They also offer refuge from extreme temperature-and flow events in the main stem, contain spawning grounds or juvenile habitat, supply allochtonous food items from the tributary catchment area, and create migration corridors between the main stem and upstream lakes and smaller streams (Jackson et al. 2001; Fausch et al. 2002; Meyer et al. 2007; Rice et al. 2008; Wohl 2017).

Management actions to improve habitats in the main stems of these rivers are often costly and incongruous with hydropower production, at least with high production, but measures in their tributaries may be more feasible from a socio-economic perspective. Hence, tributaries, and the tributary mouth areas in particular, are possibly key target areas for restoration and rehabilitation activities. The aim of this study was to provide baseline information about fish biodiversity in tributary mouths in two heavily impounded boreal rivers in northern Sweden, Ume-and Lule River, both of which are classified as strongly affected by fragmentation (Dynesius & Nilsson 1994). We also investigate the effects of some key hydrogeomorphological and bottom substrate characteristics of tributary mouth areas on fish fauna composition and diversity. We focus our investigation to the mouth area in the tributaries themselves, since the confluence zone in the main stem typically is affected by the impoundments caused by damming.

## MATERIALS AND METHODS

### Survey areas

The surveyed tributaries belong to two large Swedish boreal rivers, Ume River and Lule River (Fig 1). Both rivers transverse the country in a northwest-southeast direction, starting in the alpine region bordering Norway and draining into the Baltic Sea (Bothnian Sea and Gulf of Bothnia, respectively). The rivers are of similar size (Table 1) and both have multiple hydropower impoundments along a large portion of the main stem of each river (Fig. 1). In the downstream parts of both rivers, the power production follows a run-of-the-river scheme, with the flow being regulated by storage reservoirs in the upper parts.

**Figure 1.**
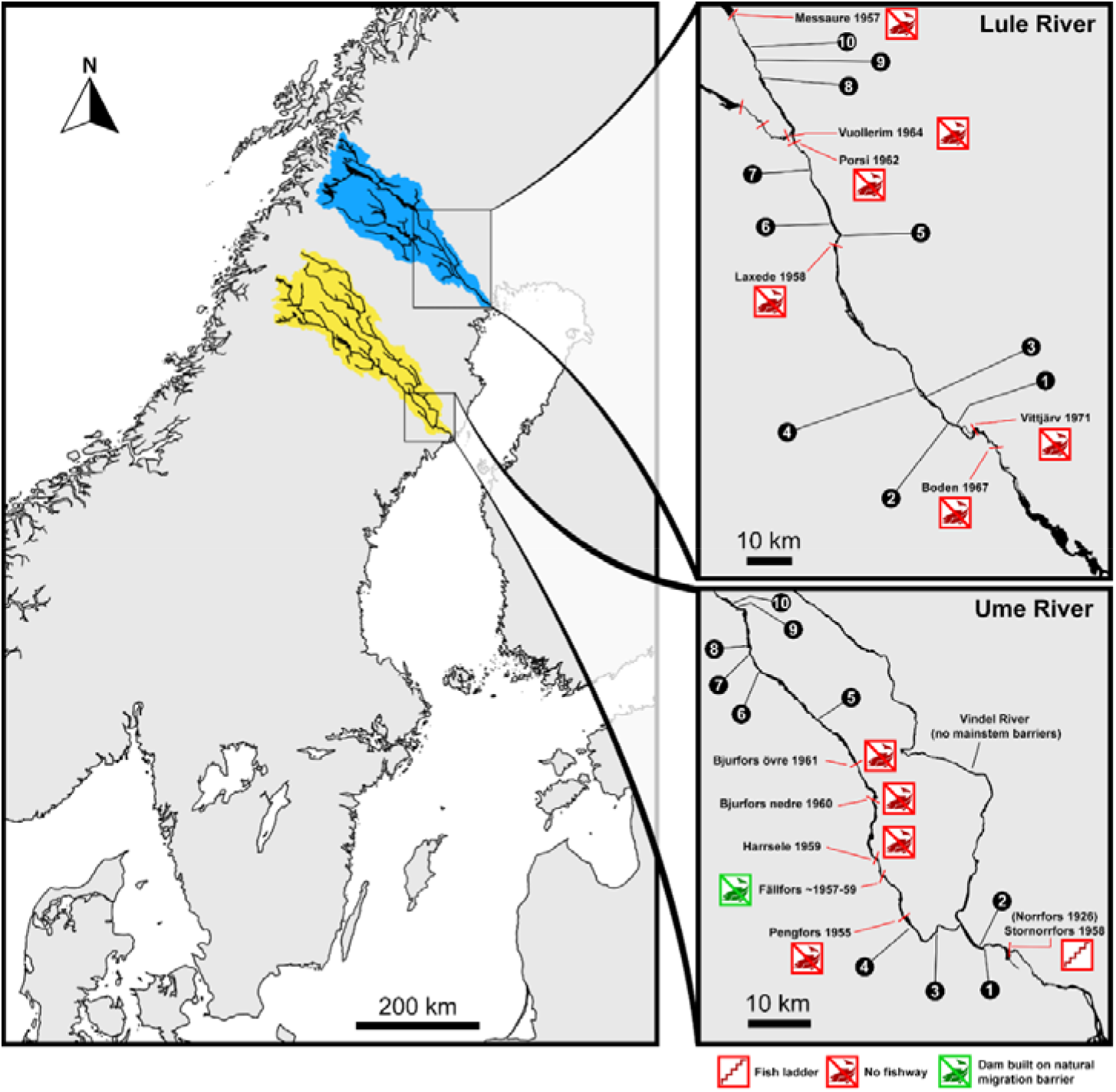
Map over Sweden with the locations of the study areas within the catchments of Ume-(yellow) and Lule (blue) Rivers. Artificial barriers are marked with red bars across the rivers in the maps. The tributaries investigated in the study are marked with numbers (for details see Table 2).

**Table 1.** Details about Ume- and Lule River (SMHI 2010; SERS 2020). MMQ: mean monthly discharge; MHQ: mean high discharge.

|  | **Ume River** | **Lule River** |
| --- | --- | --- |
| Position (mouth), WGS84 dec (N, E) | 63.75502, 20.322304 | 65.700975, 21.805115 |
| Catchment area, km<sup>2</sup> | 26782 | 25262 |
| Length (source to sea), km | 449 | 457 |
| MMQ (1900's), m<sup>3</sup> · s<sup>-1</sup> | 443 | 506 |
| MHQ (1900's), m<sup>3</sup> · s<sup>-1</sup> | 1365 | 1040 |
| Number of electrofishing records* | 2378 | 224 |

\* from the Swedish Electrofishing Register (SERS 2020-10-14)

The tributaries included in this study were limited to the region between the most downstream hydroelectric dam and the highest coastline after the Weichselian glaciation, so as to be able to work with comparable fish communities. Tributaries in Ume River were distributed over a main stem distance of 84 km (most downstream tributary located 30 km from the sea) and tributaries in Lule River were distributed over a main stem distance of 104 km (most downstream tributary located 50 km from the sea) (Fig. 1).

### Site selection

Tributaries entering Ume-and Lule Rivers have mouth areas that can be broadly classified as either aggrading or non-aggrading, which we classified based on the presence or absence of sediment plumes at the confluence, using digital aerial map surveys from public web-services (https://maps.google.se/; https://eniro.se/). We identified candidate tributary mouths based on the aerial photographs, extracted information on mean discharge (MQ) for each candidate from the S-HYPE model of national hydrological statistics (Bergstrand et al. 2014) and thereafter selected a set of tributaries along a wide range of MQ for both confluence types. Impoundments with only one type of tributary (aggrading/non-aggrading) were excluded and tributaries assessed as too deep for wading, or culverted or dammed directly at the mouth, were not considered. From the candidate set, we selected tributaries so that each confluence type was represented by equal numbers of streams (5 of each type in each of the two rivers), covering similar ranges of MQ. Assessment of tributary mouth type was also made in the field (see *Environmental survey methodology*), and final designation was based on the combined information of aerial map surveys and field measurements. The selected tributaries were located in two impoundments in Ume River and three impoundments in Lule River (Table 2).

**Table 2.**
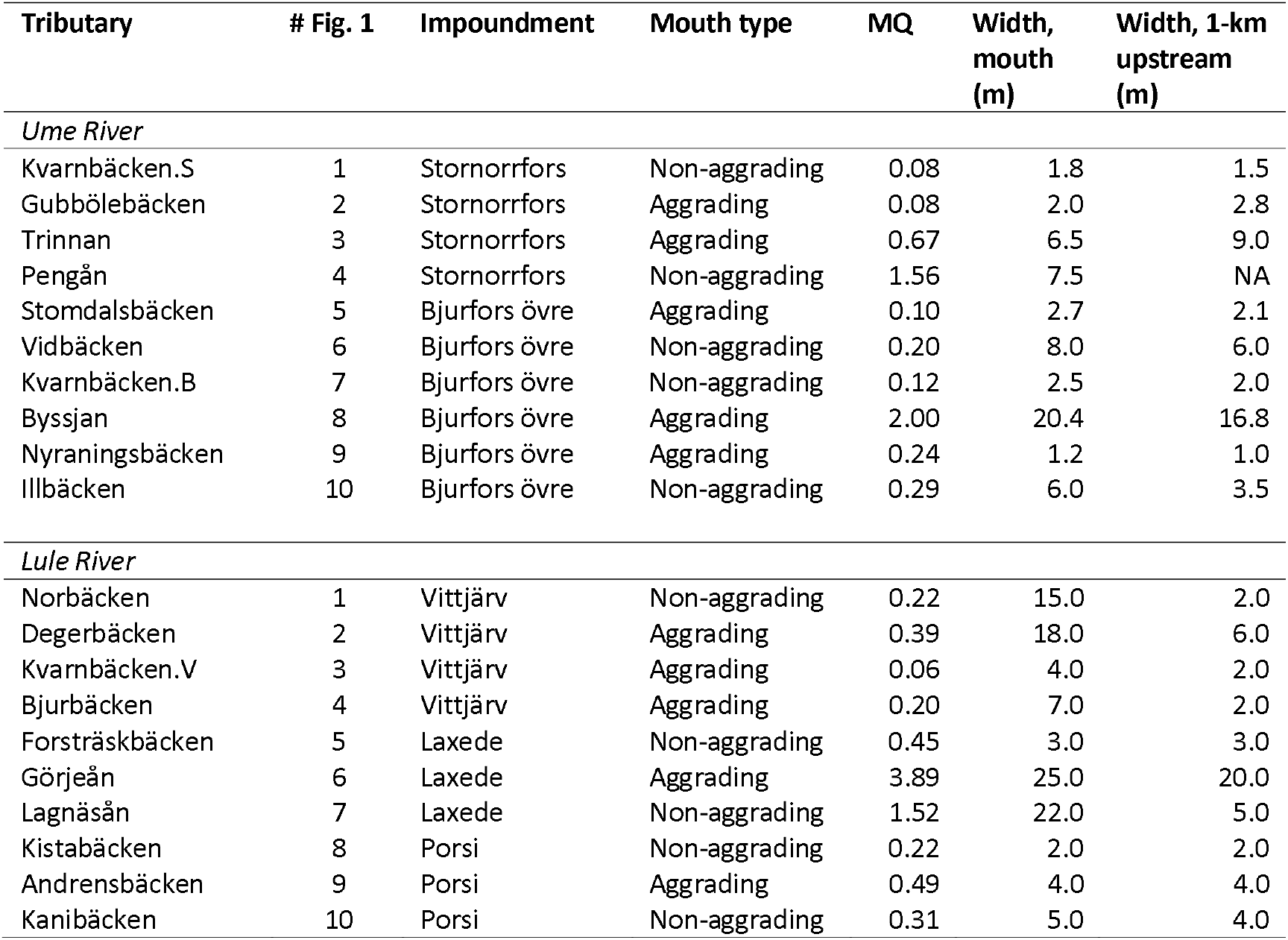
Surveyed tributaries used in the study. For location, refer to Fig. 1 following the column ‘# Fig. 1’. Note that three different tributaries are all named “Kvarnbäcken” and distinguished based on which impoundment they are located in (“.S”, “.B”, and “.V”).

### Fish communities

Fish communities in the two river systems are similar (Table 3; see references in the table legend). The main differences are the absence of Siberian sculpin *Cottus poecilopus* in Ume River records, and the absence of river lamprey *Lampetra fluviatilis* in Lule River records (GBIF 2020). River lamprey, however, does not occur upstream of the first dam in the Ume River. Anadromous salmonids (Atlantic salmon *Salmo salar* and brown trout *S. trutta*) can use a fishway to pass the same dam, reaching the most downstream surveyed impoundment of Ume River, but are either not expected in the survey area (*S. salar*, which mainly migrates up the large tributary Vindel River) or have local non-migratory populations (*S. trutta*), making species presence effectively equivalent in both rivers. Arctic charr *Salvelinus alpinus* mainly occurs in the alpine region of the catchments and populations in the lower parts are likely stocked into lakes, and hence not expected in the river tributaries. Two non-native salmonid species are present in the area, rainbow trout *Oncorhynchus mykiss* and brook charr *Salvelinus fontinalis*; the former does not reproduce and stems from unintentional escape-events from aquaculture net-pens; the latter was intentionally introduced during the 19-20^th^ centuries and established in some tributaries. With respect to historical electrofishing data, Lule River is less surveyed than Ume River (Table 1; SERS 2020). However, the sources of presence/absence data for the catchments include not only electrofishing surveys but also descriptions from the literature (Lundberg 1899, Ekman 1922; Widén et al. 2016) and data in the GBIF database (GBIF 2020). The species list also correspond with recent national distribution maps (Kullander et al. 2012).

**Table 3.**
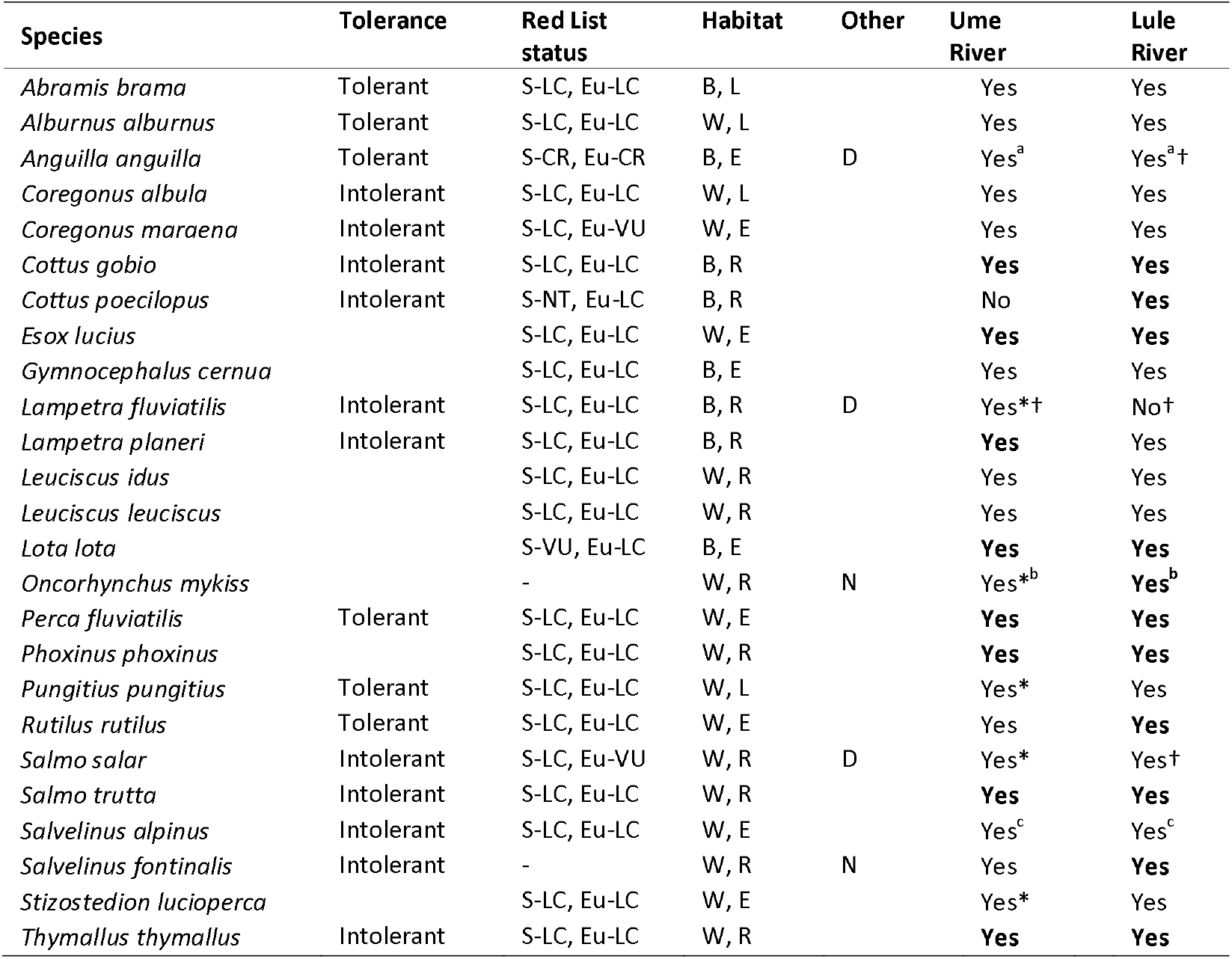
Summary of historical information on the fish species in the surveyed catchments, and the species general tolerance, Red list status and habitat preference (feeding habitat and rheophily). Red list status: S = Sweden, Eu = Europe, LC = least concern, DD = data deficient, NT = near threatened, VU = vulnerable, EN = endangered, CR = critically endangered; non-native species not considered. Feeding habitat: B = benthic, W = water column; rheophily: R = rheophilic, L = limnophilic, E = eurytopic; other: D = obligate diadromous life cycle, N = non-native. * Only recorded outside of the surveyed area. † Migration into the survey area hindered by dam. ^a^ Historically rare. ^b^ Non-reproducing. ^c^ Native to the catchment, but likely stocked within the surveyed area, native populations exists in the alpine region of the river. Record sources: GBIF (2020), Lundberg (1899), Ekman (1922), Widén et al. (2016). Species captured in this study are marked in bold font in their respective river column.

For each species, we collected additional information about tolerance, Red List status (classification of the extinction risk of a given species within a specified geographic area), and habitat preference. Tolerance, reflecting species sensitivity to impacts related to altered flow regime, nutrient regime, habitat structure and water chemistry, was based on assignments within the European Fish Index (FAME Consortium 2004; Pont et al. 2006). Red List status was obtained from the most recent lists for Sweden (ArtDatabanken 2020) and Europe (IUCN 2020). Feeding habitat and rheophily were obtained from the freshwaterecology.info database (Schmidt-Kloiber & Hering 2015). We modified the classification of brook charr to rheophilic (eurytopic in the database); this species is eurytopic in its native distribution range in North America (Scott & Crossman 1973), but naturalized Swedish populations are primarily found in headwater streams (Kullander et al. 2012).

### Fish survey methods

Multipass wading electrofishing surveys were conducted in autumn (August-September) by professional electrofishing consultants (equipment: straight-DC bank-side aggregates; Lug AB L-1000, Luleå; 700-800 V, 0.3-0.4 A), following Swedish standard practices, the so-called three-pass protocol (Bergquist et al. 2014). In each tributary, one site at the tributary mouth (i.e. near the confluence with the main stem) and one site located 1 km upstream of the mouth were electrofished. At sites with deep sections, the survey was limited to the wadable section along one of the banks, in line with Swedish standard methodology. One upstream site (Pengån, Ume River) remained unfished, as it was too deep for wading when visited. For each electrofishing pass at a given site, all fish caught were counted, identified to species, and measured for length and mass.

### Environmental survey methodology

A standardized environmental survey was conducted at each site and coordinated with the electrofishing survey, following the Swedish electrofishing protocol (Bergquist et al. 2014). In addition, a more detailed investigation of the tributary mouths was made in association to a parallel study on vascular plants in and around the same tributaries (R. Jansson, B. Malm-Renöfält, *in prep*.). Based on field observations, each mouth area was classified as 1) aggrading or non-aggrading, 2) sheltered or non-sheltered, 3) situated in an embayment or not, 4) influenced by ice-disturbance or not, 5) having boulders protruding above the water surface or not, and 6) having shallow areas or not. Altitude for each site was obtained from 2-m resolution altitude raster images over each catchment (GSD-Höjddata, grid 2+; The Swedish Mapping, Cadastral and Land Registration Authority, Gävle) in GIS (QGIS 3.16, QGIS Development Team 2021).

### Data handling and calculations

Density per 100 m^2^ was estimated for each species at each site using the sequential removal model for three passes of removal in the FAS R-package (‘Seber3’ model) (Seber 1984; Ogle et al. 2020). Fish species density estimates from each site were used to calculate the Shannon Diversity index (*H’*) (Shannon 1948), using the vegan R-package (Oksanen et al. 2019). *H’* is an estimate of the uncertainty of species identity when drawing a random individual (fish) from the data set (fish community at the site) and increases with species abundance and evenness (higher values = higher diversity). Based on *H*’, Pielou’s evenness (*J*’) (Pielou 1966) was also calculated (dividing *H*’ with *H*’_max_), as a measure of how close the number of individuals of each species are to each other at a site.

### Statistical analyses

#### General procedures

All statistical analyses were conducted in R Studio 1.2.5033 (RStudio, Inc., Boston). Types of models are abbreviated as follows: linear models – LM; linear mixed models – LMM; generalized linear models – GLM; and generalized linear mixed models – GLMM (see included factors and their abbreviations in Table 4). For statistical models with more than a single factor, the initial global models specified below were reduced based on the relative Akaike Information Criterion (modified for small sample size; AIC_c_) of all subordinate models (including the global model and the intercept-only model), using the MuMIn R-package (Barton 2020), to avoid uninfluential factors and thereby increase the residual degrees of freedom. To reduce the risk of excluding influential factors, the most complex model within two AIC_c_-units from the most parsimonious model was used for interpretation. When models were run as Poisson-regressions (i.e. for count data), overdispersion was tested using a one-sided DHARMa nonparametric dispersion test (Hartig 2021). If significant overdispersion was indicated, GLMs were re-constructed as quasi-Poisson regressions (Ver Hoef & Boveng 2007) and GLMMs were fitted with an additional observation-level random effect (Harrison 2014). Mixed models were constructed using the lme4 R-package (Bates et al. 2020), marginal means and contrasts were obtained using the emmeans R-package (Lenth 2021), and data processing and visualization was done within the tidyverse-suite for R (Wickham et al. 2019).

**Table 4.** Factors used in the statistical modelling.

| Factor | Abbreviation | Factor type | Levels | Units |
| --- | --- | --- | --- | --- |
| River identity | RIVER | Categorical | 2 (Ume R., Lule R.) |  |
| Impoundment identity | IMP | Categorical | 5 (Table 2) |  |
| Site in tributary | SITE | Categorical | 2 (OK, 1K) | - |
| Tributary identity | TRIB | Categorical | 20 levels (Table 2) | - |
| Fished area | FI.AR | Continuous | - | m<sup>2</sup> |
| Channel width | CH.WI | Continuous | - | m |
| Mean annual discharge | MQ | Continuous | - | m<sup>3</sup> · s<sup>-1</sup> |
| Altitude above main stem | ΔALT | Continuous | - | m |
| Aggradation at tributary mouth | AGGR | Categorical | 2 (yes, no) | - |
| Habitat; NMDS score, axis 1* | NMDS1 | Continuous | - | - |
| Habitat; NMDS score, axis 2* | NMDS2 | Continuous | - | - |

\* See section: *Statistical analyses: Dimension reduction of habitat variables, and Fig. 2.*

#### Dimension reduction of habitat variables in tributary mouths

To characterize and summarize habitat features in the tributary mouths, we conducted a non-metric multidimensional scaling (NMDS) analysis, using the vegan R-package (Oksanen et al. 2019). Habitat variables included: *i*) ordinal ranks of nine different sediment classes, *ii*) aggradation status (aggrading/non-aggrading); *iii*) presence/absence of protruding boulders, *iv*) presence/absence of shallow areas, *v*) binary classification of embayment, *vi*) binary classification of whether or not the mouth was sheltered, *vii*) ordinal rank of water turbidity, and *viii*) ordinal rank of flow velocity (see Fig. 2 for a key to the variables). We extracted two dimensions (*k* = 2) based on the Bray-Curtis dissimilarity index. Results were centred and half-scaled, variation was maximized in the first dimension by principal component rotation. Stress-value was derived based on type I-approach (weak ties).

**Figure 2.**
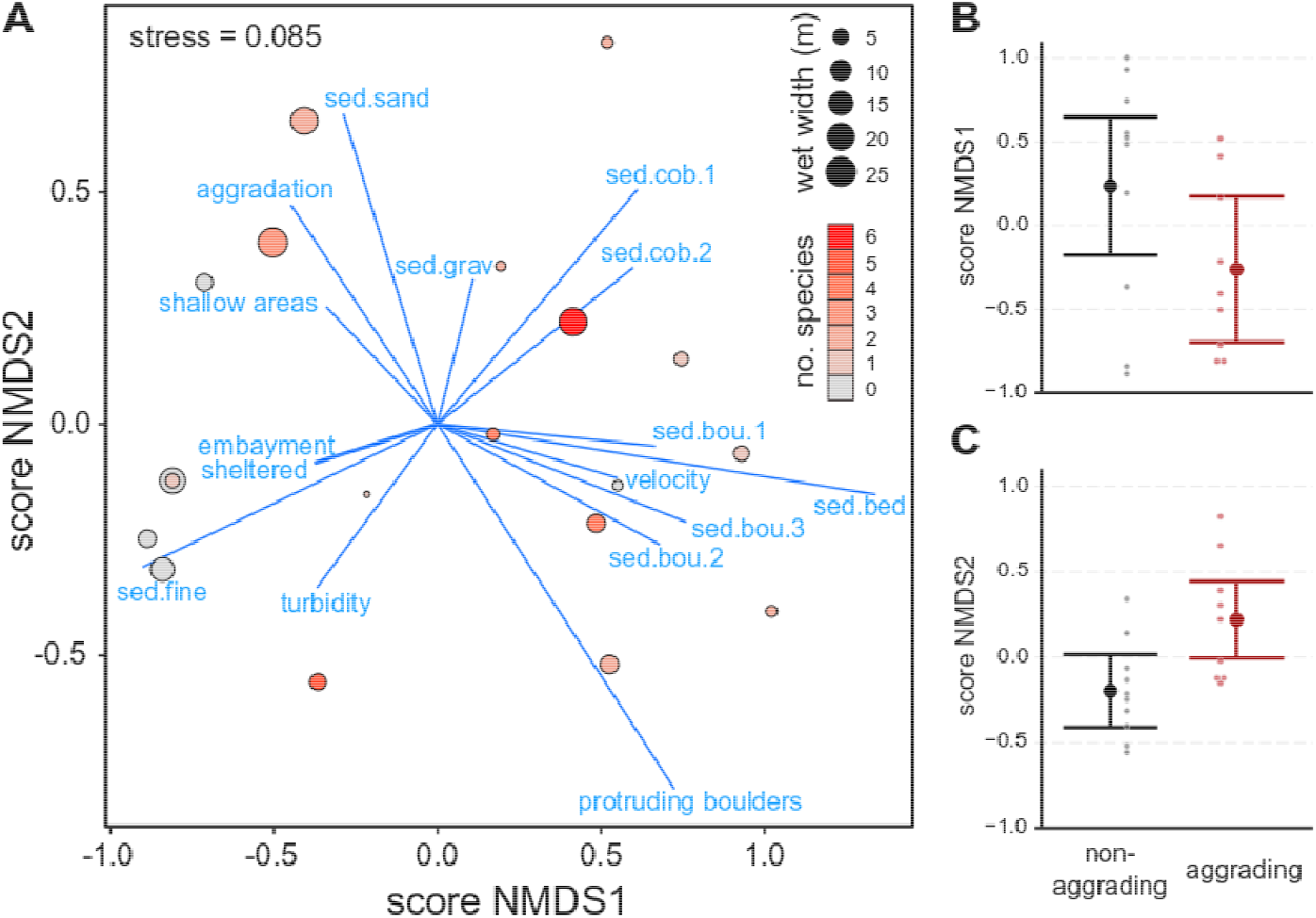
Non-metric multidimensional scaling (NMDS) and its relationship to classified aggradation status in tributary mouth areas. A) NMDS ordination in relation to number of species at each site. B) Relationship between mean score of NMDS1 and aggradation classification. C) Relationship between mean score of NMDS2 and aggradation classification. Error bars show 95% confidence interval. Key to the NMDS: ‘*sed*.*[class]*’ = sediment class on ordinal rank scale (ranks: 0 - absent, 0% coverage; 1 - scant, <5% coverage; 2 - moderate, 5-50% coverage; 3 - ample, >50% coverage; classes: fine = fine sediment <0.2mm ; sand = sand 0.2-2 mm; grav = gravel 0.2-2 cm; cob.1 = cobble 2-10 cm; com.2 = cobble 10-20 cm; bou.1 = boulders 20-30 cm; bou.2 = boulders 30-40 cm, bou.3 = boulders 40-200 cm, bed = boulders >200 cm and bedrock); ‘*aggradation*’ = binary aggradation classification (0 = non-aggrading; 1 = aggrading); ‘*protruding boulders*’ = presence of boulders protruding the surface (0 = no; 1 = yes); ‘*shallow areas*’ = presence of shallow areas (0 = no; 1 = yes); ‘*embayment*’ = tributary mouth located in a mainstem embayment (0 = no; 1 = yes); ‘*sheltered*’ = tributary mouth sheltered from the mainstem flow; ‘*turbidity*’ = ordinal rank of turbidity (0 = clear, <1 FNU; 1 = turbid, 1-2.5 FNU; 2 = very turbid, >2.5 FNU); ‘*velocity*’ = ordinal rank of flow velocity (0 = slow, <0.2 m · s^-1^ ; 1 = moderate, 0.2-0.7 m · s^-1^ ; 2 = rapid, >0.7 m · s^-1^).

#### Nuisance factors for species abundance

Initially, we ran a suite of simple one-factor Poisson-GLMs to investigate factors suspected, based on previous studies, to generally influence species abundance in the catches at the 0K and 1K sites, to make informed decisions on whether or not to include any of the factors in the construction of models specified below. Factors investigated were fished area FI.AR (Reynolds et al. 2003), channel width CH.WI (Trigal & Degerman 2015), mean annual discharge MQ (only at 0K) (Dunn & Paukert 2021), and altitude above the main stem ΔALT (only at 1K) (Lipsey et al. 2005); all but ΔALT were log_10_-transformed. No significant overdispersion was indicated in any model (all *p* > 0.064).

#### Environmental effects on fish diversity in tributary mouths

Differences in number of species and diversity (Shannon *H*’) between aggrading and non-aggrading tributary mouth areas were tested using Poisson-GLMM/GLM and LMM/LM, respectively. Global models included RIVER, AGGR, and log_10_(FI.AR) as fixed factors and IMP as a random factor (Table 4). Shannon *H*’ values were positively skewed and log_10_ transformed (Shannon *H*’ + 1) prior to analysis, which reduced, but did not eliminate the skew, due to a relatively high number of 0-values; hence interpretation of parameter estimates should be made with caution. The model reduction procedures led to exclusion of RIVER for the species-model (GLMM) and exclusion of RIVER, log_10_(FI.AR), and the random factor IMP for the diversity model (turning it from a LMM to a LM) (see supplementary material: Table S1-S2). The final species-model was not significantly overdispersed (dispersion = 0.77, *p* = 0.42).

Broader environmental habitat effects, as described by the two extracted NMDS axes, on number of species caught and diversity (Shannon *H*’) were modelled using Poisson-GLMM/GLM and LMM/LM, respectively. Global models included RIVER, NMDS1, NMDS2, and log_10_(FI.AR) as fixed factors and IMP as a random factor (Table 4). The factors NMDS1 and NMDS2 were fitted as second order polynomials and the interaction between NMDS-terms was included [in lme4-syntax: poly(NMDS1, 2)*poly(NMDS2, 2)]. As in previous analyses, Shannon *H*’ values were log_10_ transformed (Shannon *H*’ + 1). The purpose of the modelling was descriptive, rather than a test of a specific hypothesis; hence, all terms were allowed to be removed in the model reduction. The final reduced models for both number of species caught and Shannon diversity included poly(NMDS1, 2) and log_10_(FI.AR). No overdispersion of the reduced Poisson-model was indicated (dispersion = 0.65, *p* = 0.80). For model reduction details, see supplementary material: Table S3-S4.

Absence of a species or presence of only single species led to Pielou’s evenness (*J*’) not being applicable to a substantial proportion of the sites (*N* = 9). Therefore, no models were constructed for this index. Instead, Pielou data are graphed in relation to NMDS-scores, with tendencies evaluated based on loess regression and Spearman rank correlations.

#### Broader environmental effects on fish densities in tributary mouths

Environmental effects, as described by the two extracted NMDS axes, on densities of fish were modelled using LMM/LM. Fish densities were transformed as log_10_(density + 1) and analysed for six different groupings of species: i) all fish species combined, ii) tolerant species, iii) intolerant species, iv) benthic species, v) rheophilic species, and vi) species included in the Swedish Red List (see Table 3 for group classifications). Global models included RIVER, NMDS1, and NMDS2 as fixed factors and IMP as a random factor. NMDS1 and NMDS2 were fitted as second order polynomials, and their interaction was included, as described for previous analyses. All factors were allowed to be removed in the model reduction, see reduction procedure in supplementary material: Table S5-S10. The final reduced models were constructed as follows:

- All species: log_10_(density + 1) ∼ NMDS1 + NMDS2
- Tolerant species: log_10_(density + 1) ∼ 1 *(intercept-only)*
- Intolerant species: log_10_(density + 1) ∼ poly(NMDS1, 2) + RIVER
- Benthic species: log_10_(density + 1) ∼ poly(NMDS1, 2)
- Rheophilic species: log_10_(density + 1) ∼ poly(NMDS1, 2) + RIVER
- Red-listed species: log_10_(density + 1) ∼ poly(NMDS1, 2)

Final models were tested against intercept-only models using likelihood ratio tests, to assess their fit to the data. Densities were also investigated specifically in relation to NMDS1 using loess regression.

#### Differences between tributary mouth and upstream tributary sites

To compare average species richness between 0K and 1K sites within tributaries, a GLMM (Poisson, log-link) was used to model species count as dependent on SITE (fixed factor) and TRIB (random intercept); no overdispersion was indicated (dispersion = 1.08, *p* = 0.66). The same model structure was used in a LMM to model Shannon diversity (*H*’), using log_10_ transformed data (Shannon *H*’ + 1). Sign tests (two-sided) were used to compare changes (positive or negative) in species richness and Shannon *H*’ for 0K and 1K sites.

## RESULTS

### Captured fish fauna

In total, 12 species of fish (11 bony fishes and 1 lamprey) were recorded in the surveys (Table 5). In Ume River, 8 species were caught in total, 6 at the 0K-sites and 5 at the 1K-sites; the species caught at most sites were common bullhead *C. gobio* and burbot *L. lota*, each caught at 7 sites (Table 5). In Lule River, 11 species were caught, 9 at the 0K-sites and 8 at the 1K-sites; the species caught at most sites was brown trout *S. trutta*, found at 8 sites (Table 5). Non-native species (rainbow trout *O. mykiss* and brook charr *S. fontinalis*) were only caught in the Lule River system, and only at one 1K-site each; *O. mykiss* (*N* = 1; 204 mm TL) likely originated from an aquaculture net-pen in an upstream lake and *S. fontinalis* (*N* = 37) were naturalized, as indicated by the presence of age 0+ individuals (51-68 mm TL).

**Table 5.** Species caught in Ume- and Lule River, based on the number of 0K (tributary mouth) and 1K (1-km upstream) sites containing each species and the total number of individuals caught.

| Species | Ume River |  |  |  | Lule River |  |  |  |
| --- | --- | --- | --- | --- | --- | --- | --- | --- |
|  | OK sites<br>(*N*) | 1K sites<br>(*N*) | OK ind.<br>(*N*) | 1K ind.<br>(*N*) | OK sites<br>(*N*) | 1K sites<br>(*N*) | OK ind.<br>(*N*) | 1K ind.<br>(*N*) |
| *Cottus gobio* | 3 | 4 | 26 | 26 | 1 | 1 | 12 | 1 |
| *Cottus poecilopus* | 0 | 0 | 0 | 0 | 3 | 1 | 27 | 2 |
| *Esox lucius* | 1 | 0 | 1 | 0 | 1 | 1 | 1 | 1 |
| *Lampetra planeri* | 2 | 1 | 2 | 1 | 0 | 0 | 0 | 0 |
| *Lota lota* | 6 | 1 | 8 | 1 | 2 | 0 | 2 | 0 |
| *Oncorhynchus mykiss* | 0 | 0 | 0 | 0 | 0 | 1 | 0 | 1 |
| *Perca fluviatilis* | 3 | 0 | 7 | 0 | 1 | 0 | 1 | 0 |
| *Phoxinus phoxinus* | 3 | 0 | 12 | 0 | 3 | 2 | 24 | 21 |
| *Rutilus rutilus* | 0 | 0 | 0 | 0 | 1 | 0 | 5 | 0 |
| *Salmo trutta* | 0 | 2 | 0 | 18 | 4 | 4 | 90 | 129 |
| *Salvelinus fontinalis* | 0 | 0 | 0 | 0 | 0 | 1 | 0 | 37 |
| *Thymallus thymallus* | 0 | 1 | 0 | 1 | 2 | 2 | 44 | 56 |

### Dimension reduction of habitat variables in tributary mouths

A two-dimensional ordination of environmental variables in the tributary mouth areas resulted in a final stress value of 0.085, indicating a good ordination in combination with the Shepard plot derived from the analysis (Fig. S1; Clarke 1993). The first NMDS axis (NMDS1) largely ordered sites along a substrate size and water velocity gradient, with high values being associated with large substrate sizes and high velocity, and low values with small-sized substrate (Fig. 2A). Other features like turbidity, shallow areas, and embayment, to some extent loaded in the same direction as small substrates, as did the binary classification of aggradation status. Modelling NMDS1-scores as dependent on binary classified aggradation status revealed that NMDS1 did not differ significantly between sites classified as aggrading and non-aggrading, although a tendency for aggrading sites having lower values was seen (ANOVA: *F*_*1,17*_ = 3.06, *p* = 0.098; Fig. 2B). The second NMDS axis (NMDS2) was mainly described by the presence of protruding boulders loading in the negative direction of the axis, and sandy to small cobble substrates and aggradation status loading in the positive direction of the axis (Fig. 2A). Modelling NMDS2-scores as dependent on their binary classified aggradation status revealed that NMDS2 significantly differed between sites classified as aggrading and non-aggrading (ANOVA: *F*_*1,17*_ = 8.04, *p* = 0.011; Fig. 2C).

### Nuisance factors for species abundance

For 0K-sites, channel width did not have a significant effect on the number of species caught [log_10_(CH.WI): *z* = 1.494, *p* = 0.135], but mean annual discharge and fished area had positive effects [log_10_(MQ: *z* = 2.802, *p* = 0.005; log_10_(FI.AR): *z* = 1.987, *p* = 0.047] (Fig. S2A-D). For 1K-sites, channel width and fished area had significant effects on number of species caught [log_10_(CH.WI): z = 2.319, *p* = 0.020; log_10_(FI.AR): *z* = 3.162, *p* = 0.002], but altitude (in relation to the mainstem) did not (ΔALT: *z* = -0.924, *p* = 0.356) (Fig. S2E-H). Looking at all sites combined, channel width, mean annual discharge, and fished area were all strongly correlated with each other (all *r* > 0.7, all *p* < 0.05; Fig. S3). Hence, these three variables appear to largely represent the same thing (i.e. size of the tributary), and since fished area consistently had a positive effect on number of species caught, this variable was used in the more detailed modelling.

### Environmental effects on fish diversity in tributary mouths

Models investigating effects of binary aggradation status of the tributary mouth area indicated no statistically significant effects on number of species (ANODEV; AGGR: *χ*^2^ = 0.26, *p* = 0.61; log_10_(FI.AR): χ^2^ = 1.67, p = 0.20; Fig. S4a). Similarly, no significant effect on diversity, as indicated by Shannon *H*’, was found (ANOVA; AGGR: *F*_1,18_ = 0.45, *p* = 0.51; Fig. S4b).

The model for number of species indicated significant effects of the second-order polynomial term poly(NMDS1, 2) (*χ*^2^ = 15.8, *p* < 0.001) and log (FI.AR) (*χ*^2^ = 5.26, *p* = 0.022). Parameter estimates indicated a concave down relationship between number of species caught and NMDS1 (Fig. 3A; parameter estimates: *β*_intercept_ = -3.43, *β*_NMDS1:1_ = 2.26, *β*_NMDS1:2_ = -2.94, *β*_log10(FI.AR)_ = 1.56). As the first axis of the NMDS generally relates to substrate classes and their associated habitat characteristics, the pattern suggests that the intermediate substrate size is associated with the highest number of species. The number of species caught also increases with fished area, as also indicated in the previous analyses. The factor NMDS2 was not retained after model reduction, indicating that it is not influential on the number of species present (see relationship in Fig. S5A).

**Figure 3.**
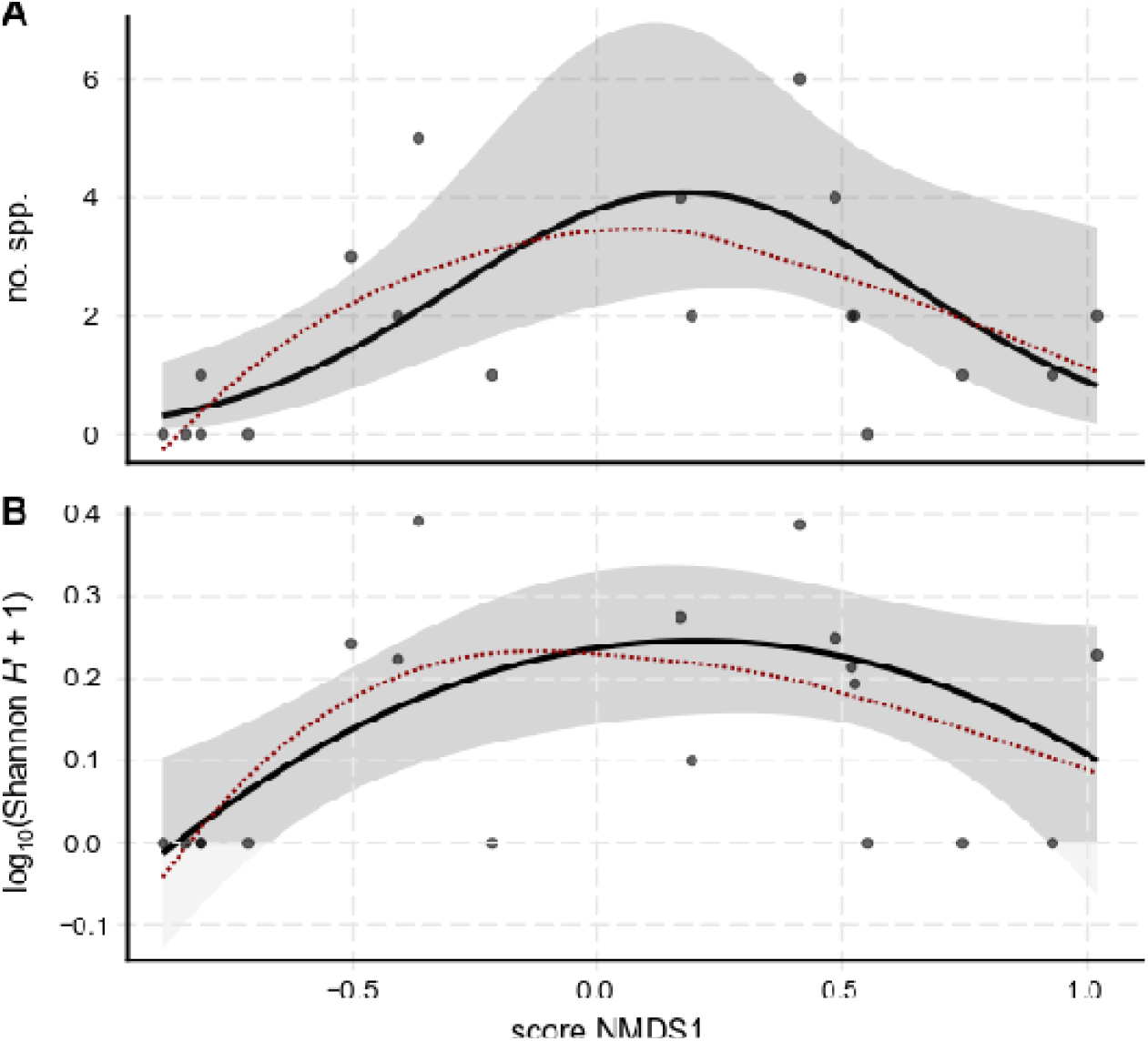
A) Effect of the first NMDS axis (NMDS1) scores on the number of species caught in the tributary mouth areas. Black line shows the modelled effect from a Poisson GLM, with 95% confidence bands in grey. Red dotted lines show a loess regression on raw data.

The model for Shannon diversity (*H*’) also indicated significant effects of the second-order polynomial term poly(NMDS1, 2) (*F*_2,15_ = 5.49, *p* = 0.016) and log_10_(FI.AR) (*F*_*1,15*_ = 5.71, *p* = 0.030). Parameter estimates indicated a concave down relationship with NMDS1 (Fig. 3B; parameter estimates: *β*_intercept_ = -0.47, *β*_NMDS1:1_ = 0.25, *β*_NMDS1:2_ = -0.28, *β*_log10(FI.AR)_ = 0.25). In accordance with number of species, the pattern suggests that intermediate substrate size is associated with higher diversity, and increased diversity with fished area. The factor NMDS2 was not retained after model reduction (see relationship in Fig. S5B). Pielou’s evenness (*J*’) was not significantly rank-correlated with any of the investigated environmental variables [NMDS1, NMDS 2, FI.AR, and CH.WI], with all Spearman rank correlations having *p* > 0.46 (see Fig. S6 for details and visualization of loess-regressions).

### Broader environmental effects on fish densities in tributary mouths

For the model of total fish density (all species combined), the factor *NMDS*1 was significant, indicating increasing density with increasing values of *NMDS*1 (Fig. 4A), while *NMDS*2 was non-significant (Table 6). Based on the loess regression fit, the significant positive relationship appears largely driven by low fish densities in sites dominated by fine sediment, with associated environmental features (i.e. sites with the lowest values on the *NMDS*1-scale). For all other sites, no apparent trend was indicated in the loess fit (Fig. 4A).

**Figure 4.**
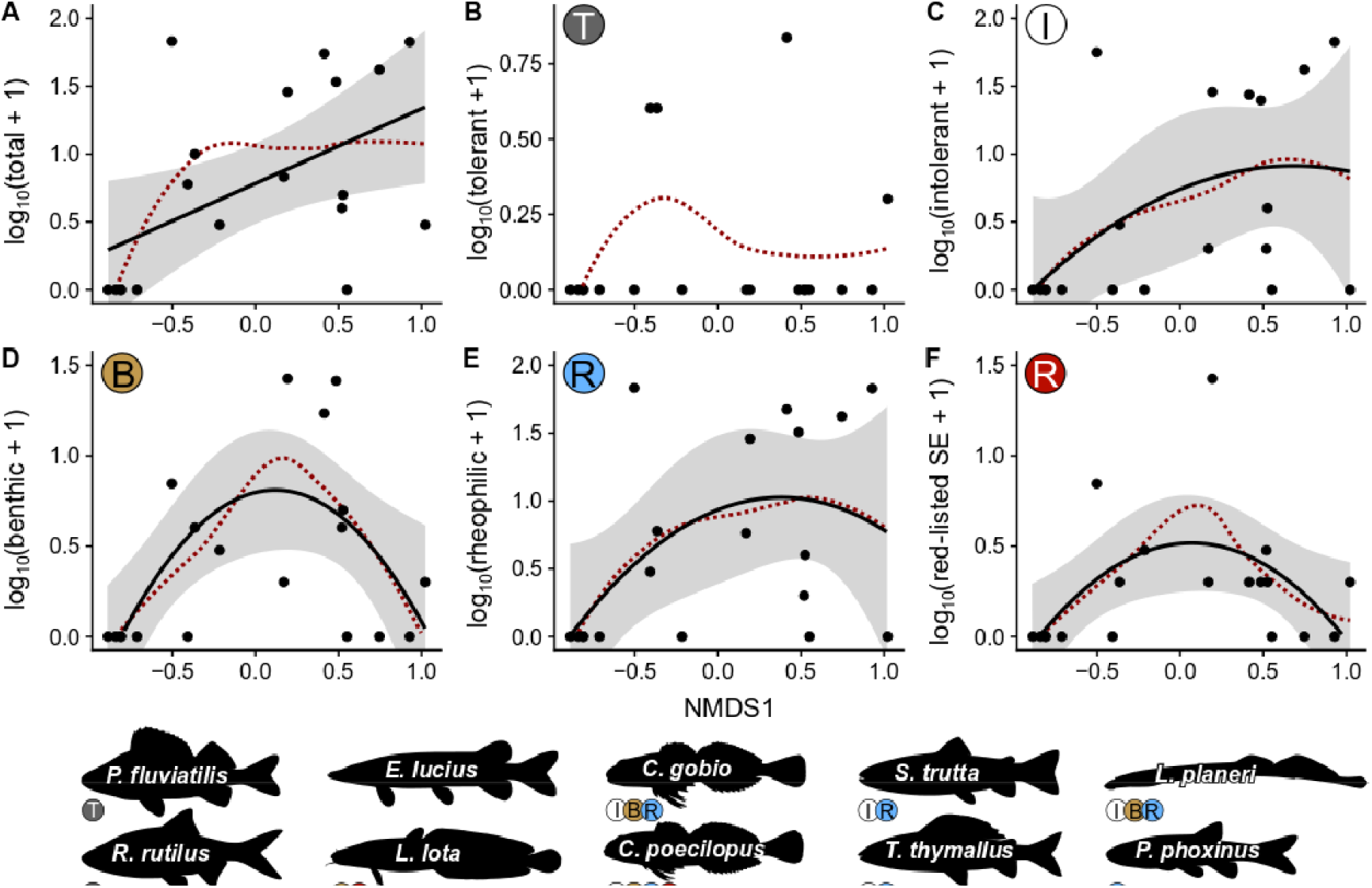
Densities of different fish guilds (including lampreys) along the first NMDS axis (finer to coarser substrate, from left to right): A) total density of fish, B) density of tolerant species, C) density of intolerant species, D) density of benthic species, E) density of rheophilic species, F) density of species listed in the Swedish Red List (Artdatabanken 2020). Black lines: regression lines from linear models, with grey-shaded areas representing the 95% confidence band (not provided in B; tolerant species were only present at four sites). Red dotted lines: loess regression lines based on raw data. Species present in the data illustrated with silhouettes, scientific name and the categories to which they belong (labels under the silhouettes matching labels in the graphs).

**Table 6.**
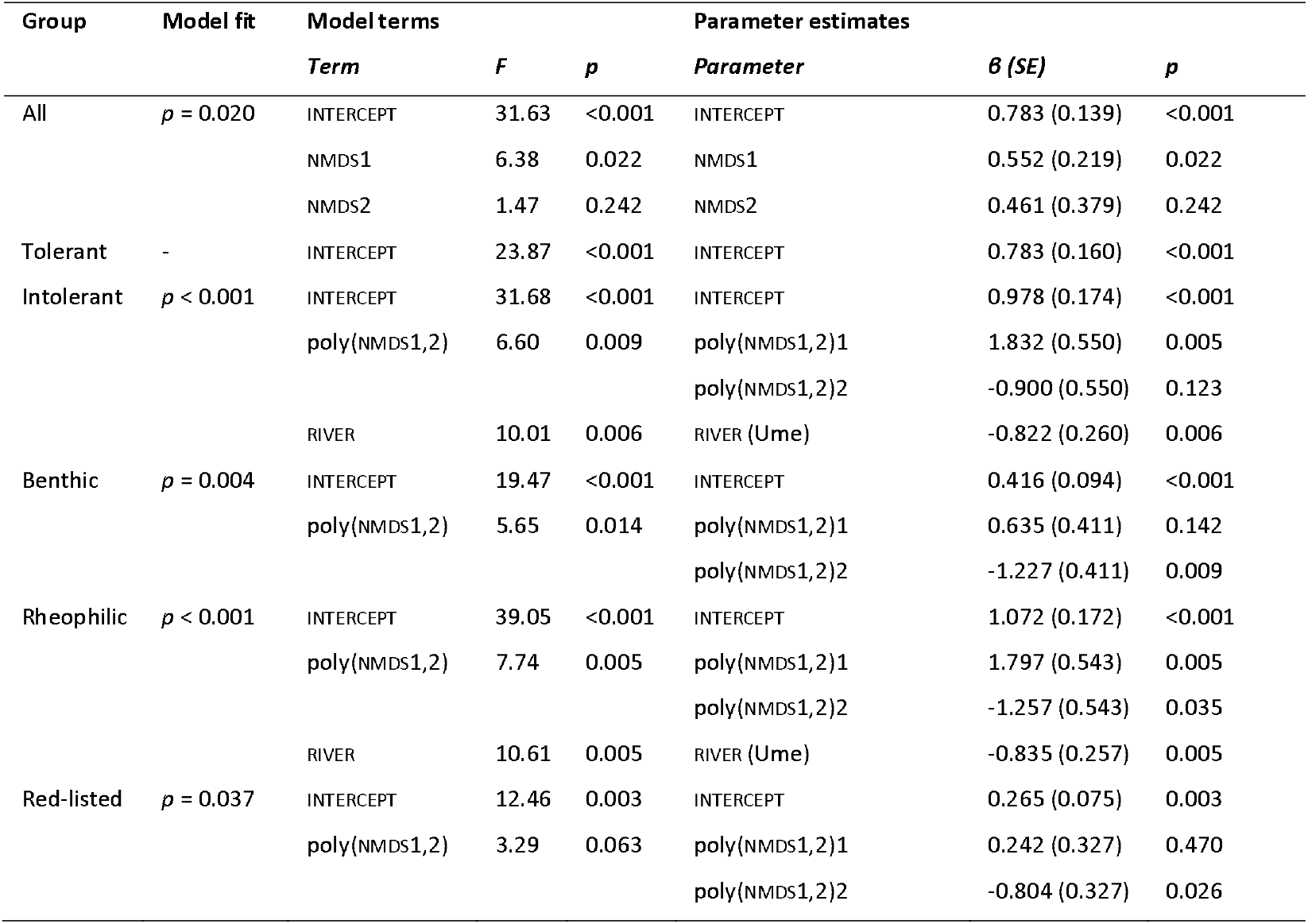
Summary of analyses of fish densities, with model fit (comparison with intercept-only model), significance of model terms, and parameter estimates from the linear models. Note that estimates for polynomial terms [poly()] relate to orthogonal polynomials.

For tolerant species, only four sites had tolerant species present (Fig. 4B), which is inadequate for modelling the influence of environmental factors. The model selection procedure ended with selection of the intercept-only model, and hence, no factors were specifically investigated (Table 6).

In the model of intolerant species density, the polynomial of *NMDS*1 and *RIVER* were retained in the model selection procedure (which was designed to allow for some model complexity; not always selecting the most parsimonious model) and both terms were significant. Parameter estimates, however, indicate that the polynomial relationship with NMDS1 is not significantly different from a linear fit (Table 6; Fig. 4C). This is supported by the fact that neither a likelihood ratio test (LRT) nor AIC_c_-comparisons indicate that the fit of the polynomial model is significantly better than a linear fit (LRT: *p* = 0.102; ΔAIC_c_ = 0.64). Loess regression suggests that the fit may be driven largely by a lack of intolerant species in the sites with the lowest NMDS1-scores. For sites with higher NMDS1-scores (> -0.5), there is no trend and intolerant species’ densities vary substantially throughout this range (Fig. 4C). Ume River had a lower fish density in general (Table 5).

Density of benthic species showed a concave down relationship with NMDS1 (Table 6; Fig. 4D). This pattern was also supported by the loess regression.

Density of rheophilic species was modelled using a polynomial function of NMDS1, and this model indicated a potential concave down relationship (Table 6; Fig. 4E). While the polynomial fit is not obviously different from a positive linear fit, both LRT and AIC_c_ comparisons of the models indicate that the fit of the polynomial model is significantly better (LRT: *p* = 0.021; ΔAIC_c_ = 2.04). Similar to density of intolerant species (which constitute a subset of the rheophilic species), assessment of the loess regression suggests that the fit can be driven largely by the lack of rheophilic species in the sites with the lowest NMDS1-scores. An effect of RIVER was also detected, with Ume River having a lower density of rheophilic species (Table 6).

Density of red-listed species followed the same concave down relationship with NMDS1 as the density of benthic species (Table 6; Fig. 4F). This result is consistent with the fact that red-listed species constitute a subset of the benthic species.

From a local conservation perspective, presence of specific species in relation to habitat characteristics can be of importance. Hence, we present a graphical representation of each species caught within this study in the Appendix (Fig. A1).

### Differences between tributary mouth-and upstream sites

Neither species richness nor Shannon diversity differed significantly between 0K-and 1K sites (species richness: *z* = -1.542, *p* = 0.123; Shannon *H*’: *t* = -1.546; *p* = 0.139; Fig. 5). Based on preliminary analyses of possible nuisance factors (see above), altitude was not considered in the models and further support for this decision was in the lack of a correlation between the change in species richness or Shannon *H*’ and the change in altitude ΔALT between 0K-and 1K sites (species richness: *r* = -0.267, *p* = 0.269; Shannon *H*’: *r* = -0.173, *p* = 0.479). Sign tests comparing general patterns of increase or decrease between 0K-and 1K sites did not indicate any significant differences (species richness: *z* = 1.508, *p* = 0.132; Shannon *H*’: *z* = 1.155; *p* = 0.248; Fig. S7), in line with the GLMM/LMM analyses above.

**Figure 5.**
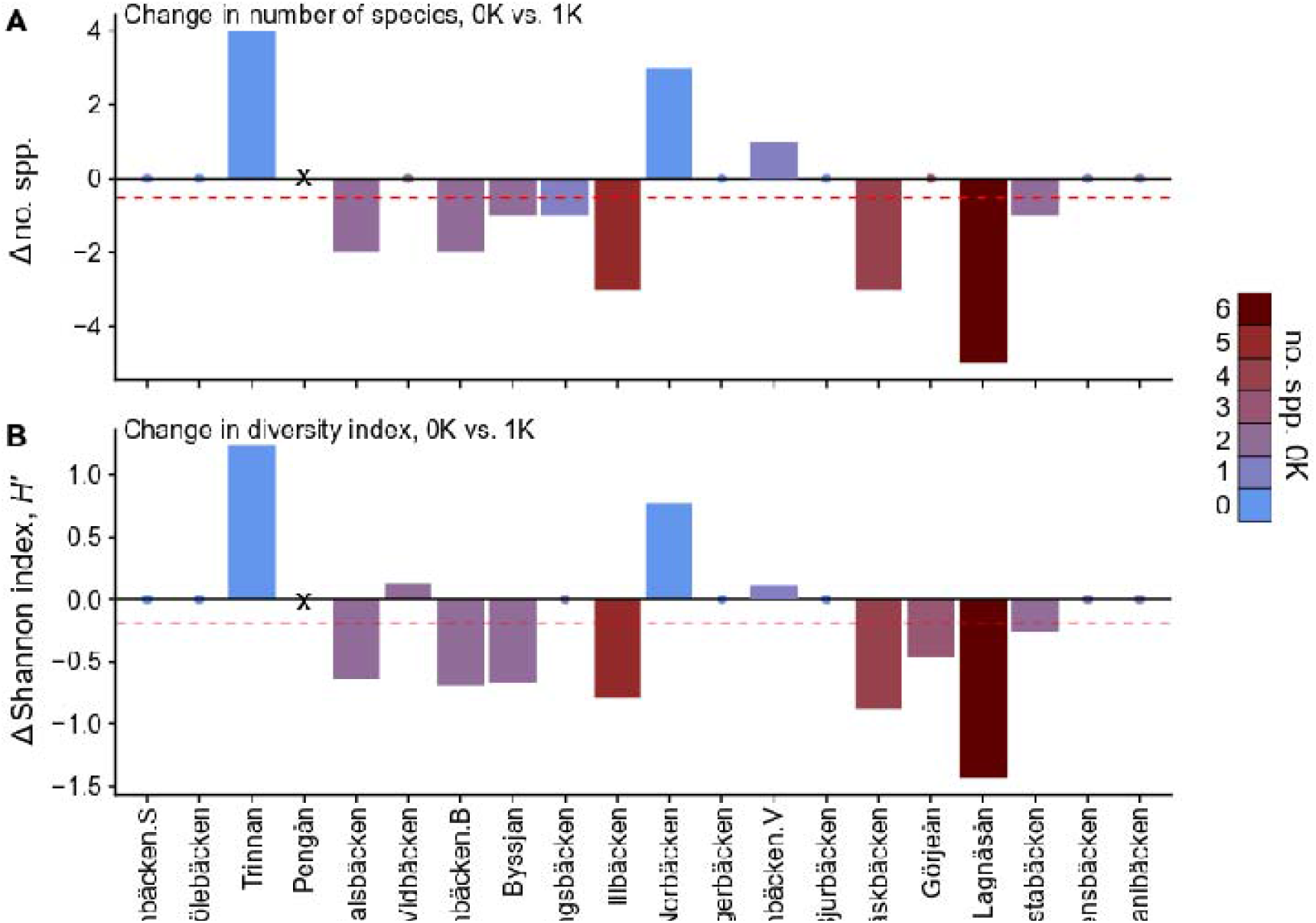
Comparison between sites located at the mouth of tributaries (0K) and sites located ca. 1 km upstream these tributaries (1K). A) Difference in species richness. B) Difference in Shannon index (diversity). Positive values indicate an increase from 0K-to 1K-sites and negative values indicate a decrease. Color scale represents number of species at the 0K sites. Red dashed lines show the arithmetic mean difference. In Pengån, no 1K site could be sampled (marked ‘X’).

Generally, the species present at the mouth of the tributary were not the same species as encountered upstream (Fig. A2). Regressing change in species richness as dependent on the species richness at the 0K site gives a slope of -0.87 (95% CI: -0.50 – -1.23) which suggests a strong regression-to-the-mean effect, which could be due to more or less random distribution of data for 0K and 1K sites (e.g. Barnett et al. 2005).

## DISCUSSION

### Fish biodiversity in relation to habitat characteristics

In the investigated sections of the impounded boreal Ume- and Lule Rivers, tributary mouths characterized by intermediate sediment-sizes (gravel-cobble) had the highest α-diversity (species richness) of fish. The Shannon diversity index, which incorporates also the relative abundance of the species present, showed a similar pattern, although not statistically significant. The same areas tended to harbor higher densities of benthic species, which generally may be favored by a high environmental complexity (possibly with the exception of brook lamprey *L. planeri*, which bury in finer sediments). In line with this observation, the two red-listed species (Artdatabanken 2020), burbot *L. lota* (VU) and alpine bullhead *C. poecilopus* (NT), both benthic species, showed higher densities at intermediate substrate sizes. While abiotic effects often dictate the ecological community in freshwater ecosystems (Jackson et al. 2001), a similar study from a more species-rich river system in central Europe found no clear detectable effects of habitat structure on species richness (Czeglédi et al. 2015). The discrepancy in effects suggests that the importance of local habitat may differ depending on biogeographical factors (Grenouillet et al. 2004) and extrapolation of results outside of the investigated area may be difficult.

Our initial classification based on aggradation status, based on presence/absence of sediment plumes at the tributary mouth, did not relate to any clear statistical differences in fish diversity. This classification mainly predicted the pattern on the second NMDS axis (NMDS2) and analyses including NMDS2 instead of the binary aggradation class did not suggest any relationship between this axis and species richness or Shannon *H*’.

### Fish abundance

Densities of fish in generally increased with coarser substrate, but it is not clear whether densities increase continuously with increasing sediment size (NMDS1 value), or if there is a non-linear saturating or step-wise effect where densities increase from fine sediments to gravel substrate, and then remain at a stable level with increasing substrate size (as indicated by the non-linear loess regression in Fig. 4A). In many cases, non-linear effects in response to habitat complexity are expected (Soukup et al. 2022), but more detailed investigations are needed to resolve this question. Species contributing to high densities in coarser-substrate tributary mouth areas tended to be rheophilic and intolerant to anthropogenic impacts (these two guilds have largely overlapping species composition; Pont et al. 2006; Schmidt-Kloiber & Hering 2015). Where large-sized substrates dominates, stream power (streamflow × slope) is typically higher as compared to sites where smaller substrates dominate (Lane 1955), which can explain why occurrence of rheophilic species is higher at sites with higher values of NMDS1 (i.e. characterized by larger substrate).

With regards to the two rheophilic taxa of national management concern, the bullheads (*Cottus spp*.) and brown trout (*S. trutta*), we noted that the former are found with their highest densities at intermediate NMDS1 values (cobble to gravel, or even sandy habitats), while the latter have its highest densities at high NDMS1 values (i.e. boulder habitats). These patterns fit with previous observations in Norwegian subalpine rivers (Hesthagen et al. 2004). The differences in habitat preference for these two taxa, which both are important for environmental management, illustrate the importance of maintaining environmental variation across sites.

Predominantly limnophilic species were not common in this study, not even in slow-flowing tributary mouths dominated by finer substrates. These latter types of sites were generally shallow over substantial areas so, for limnophilic species, the main stems or lakes may be more suitable as habitats within the river systems. Moreover, it was not possible to electrofish deep areas when they occurred, and this may be biasing our results as indicated previously (Cooke et al. 2012). Limnophilic species may also be more mobile, only inhabiting tributary mouth areas temporarily, which decreases the probability of them being caught during single electrofishing surveys. Seasonal and diel changes in habitat-specific species composition have been described in several river systems (Copp & Jurajda 1999; Nunn et al. 2010; Salas & Snyder 2010). Hence, sampling over a broader period of time might have produced a different picture of fish biodiversity in tributary mouths.

#### Fish α-and γ-diversity in broader context

The number of fish species found in each tributary mouth ranged from 0 to 6, indicating a generally low α-diversity. At sites located 1 km upstream of the tributary mouths, species richness ranged from 0 to 4, but without any clear decline within a given tributary. Furthermore, the species present generally differed between the mouth and the upstream site in the tributaries. Hence, the species found in the upstream areas are not typically a subset of the species present in the mouth-area, but may constitute a different type of community. To some extent, these findings contrast with results from central Europe, where a decline in tributary fish α-diversity could be detected from the mouth to sites located 1 km upstream (Czeglédi et al. 2015).

Low species richness likely reflects the overall γ-diversity in these boreal rivers. Only 24 species in total are known from the two investigated river systems (Table 3), out of which four are not expected in the survey areas due to downstream migration barriers (*A. anguilla, S. salar, L. fluviatilis*) or alpine distributional limits (*S. alpinus*). Considering that the α-diversity at a given tributary mouth area constitutes only 0-30% of the expected γ-diversity and that no systematic differences were found between mouth and upstream sites, it is questionable whether the tributary mouth areas can be considered biodiversity hot-spots for fish in these systems.

#### Caveats and future research requirements

The present study focuses on tributary mouth fish diversity in impounded rivers, as assessed from electrofishing surveys. As such, the study constitutes a first comparative insight into the fish diversity in these areas, but some key information is still missing – especially for designing appropriate management actions. For instance, the study does not provide information about the biodiversity in unimpacted reference systems. To gain this knowledge future studies could survey the tributaries in the few remaining unimpacted boreal rivers (or river sections) in Europe. Surveys in the present study did not cover the biodiversity in the main stems, due to lack of methods comparable to wading electrofishing being possible to perform in these deeper habitats. Boat electrofishing could be considered in future studies, but differences in species-specific capture bias between wading-and boat electrofishing present a problem for comparisons. Furthermore, efficient boat electrofishing is only possible to a depth of a couple of meters, making it difficult to survey the main stem fish fauna representatively.

Surveys conducted at different points in time is another focus area that could be investigated in future studies to improve our knowledge about the importance of tributary mouth areas as fish habitats. Many juvenile species use these shallower habitats mainly at night (Copp & Jurajda 1999), and given that we only have survey data from daytime here, information about nocturnal behavior and community composition through the diel cycle is currently lacking in these systems. These studies would likely require a different survey method since wading electrofishing at night could be hazardous. Environmental DNA do not have high enough spatial resolution for detailed studies, but quantitative metabarcoding may give insights into short-term changes in species presence. Other survey methods like e.g. snorkeling transects, trapping, or possibly seining may be better suited; all of which, however, may be primarily applicable in deeper, slow-flowing, areas. Similarly, we do not have information about usage of these habitats during spring, summer, or winter, and seasonal differences have at least been detected in a similar study in from a more species-rich system in central Europe (Czeglédi et al. 2015).

### Management considerations

In terms of the value of tributaries for the biodiversity in larger impounded rivers, the mouth areas of the tributaries are not particularly rich in fish species in these northern rivers. However, rheophilic species are disfavored in impounded rivers since their typical habitats, riffles and rapids, are often either inundated, or completely or partially dried out when bypassed by hydropower infrastructure (Malm Renöfält et al. 2010; Göthe et al. 2019; Widén et al. 2021). To maintain as high ecological potential as possible, natural tributary mouths characterized by flowing habitats and medium to large sediment substrate could be protected from further anthropogenic impacts, as they may constitute near-main stem refuges for rheophilic species and as migration areas to upstream suitable habitats. Where degraded (e.g. by dams or culverts near the confluence), these types of habitats could also be rehabilitated or restored.

To promote the currently red-listed species present in the area, *L. lota* and *C. poecilopus*, habitat measures may consist of ensuring high enough availability of medium-sized sediment habitat. Habitat restoration efforts in Swedish boreal rivers typically promote brown trout habitats (i.e. coarser-sediment habitats) (Degerman & Näslund 2021), which might disfavor bullhead, given the apparent competition between trout and bullheads (Hesthagen et al. 2004). Tributary mouth areas could possibly be appropriate target areas for (re-)creating intermediate sediment-size habitats and trout may instead be targeted in the upstream areas of the tributaries. Also the burbot might find a refuge from negative thermal-, flow-and pollution impacts in the main stem (Stapanian et al. 2010; Dugdale et al. 2013; Koizumi et al. 2013; Artdatabanken 2020; Wang et al. 2020).

To promote γ-diversity in the river systems at large, we need more knowledge about the diversity in the rivers, e.g. how the different species are distributed through the river networks. It could, in fact, be important to maintain a range of environmentally different types of tributary mouths to not only benefit a subset of the species present, due to competitive differences associated to habitat characteristics. Management activities must also consider other taxa than fish, including invertebrates, plants, semi-aquatic vertebrates, etc., in both the aquatic environment and the riparian zone.

Restorative action gains cumulative value if considered on a catchment scale rather than focusing on local efforts (Gann et al. 2019; Cid et al. 2021). Therefore, when a native reference state is unachievable, a combination of efforts ca be implemented. For example, reconstruction of road culverts at the tributary mouth will likely lead to a more natural flow regime (Widén et al. 2016), and recreating natural stream morphology with natural riparian vegetation in currently straightened and channelized tributaries will likely reduce the erosion of fine sediment in the tributary (Beschta & Platts 1986). Both measures may also help retaining the water in the tributary to avoid draught.

### Conclusions

With this study, we have gained information about the diversity of fish in tributary mouth areas within large impounded boreal river systems. We find that fish species richness and diversity is relatively low but variable, with variation being primarily explained by habitat features related to sediment grain sizes. Highest diversity was found in mouths with intermediate grain size (gravel-cobble). Fine sediment habitats often contain few, if any, fish species during the time of our surveys. The species composition at the mouth was generally not the same as upstream in the tributary and the species present upstream was not a subset of the species at the mouth. Overall, we find no clear evidence supporting the statement that tributary mouth areas are biodiversity hot-spot for fishes. To understand the most suitable management actions needed in these areas, further studies on the native natural state of these environments are needed, and so is a more detailed understanding of the temporal dynamics of fish communities in relation to spatial distribution of fish in tributaries and their mouths..

## Supporting information

Supplementary information: analyses

Supplementary information: Photos of study sites

## Data Availability Statement

Data from the electrofishing surveys are deposited in the Swedish Electrofishing Register (http://www.slu.se/elfiskeregistret). Collated data sets used for analyses and R code are deposited in the figshare database (link for review purpose: https://figshare.com/s/1c373beb5b0ba497f9bc).

## Acknowledgements

We thank Birgitta Malm-Renöfält and Roland Jansson (Umeå University) for collaboration regarding selection of field sites and assistance with field assessment the selected sites. Electrofishing was carried out by Fiskmiljö Nord AB (Lule River) and EKOM AB (Ume River).

## Supplementary Materials

**Supplement 1:** Supporting analyses and figures.

**Supplement2:** Photographs of the electrofished sites.

## APPENDIX

**Figure A1.**
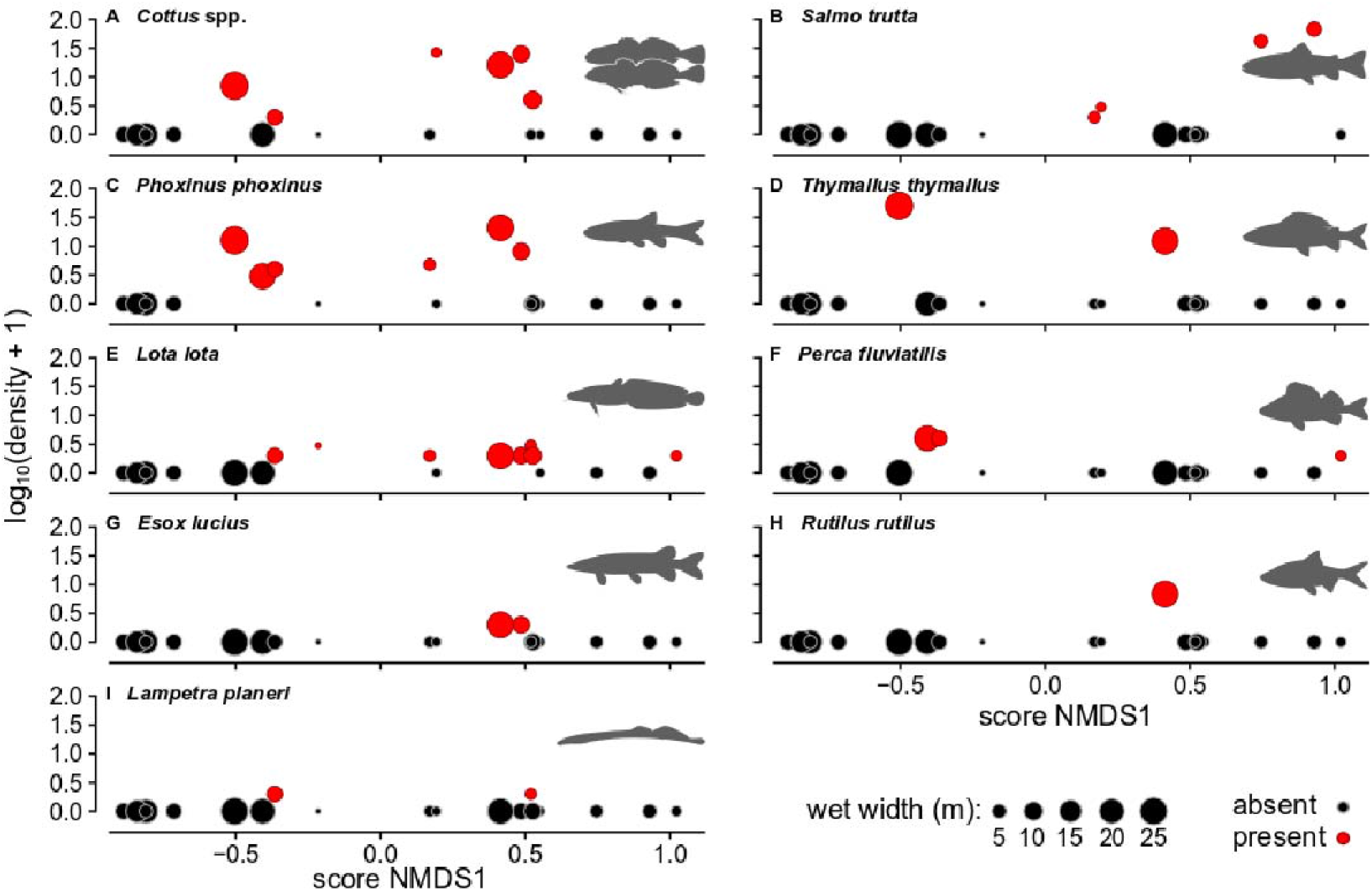
Density estimates of the different fish species caught in the tributary mouth areas, in relation to the first NMDS axis (x-axis) and wet width of the tributary at the mouth (size of the dots). Densities for *Cottus gobio* and *C. poecilopus* were pooled due to similar autecology, but only partially overlapping distribution.

**Figure A2.**
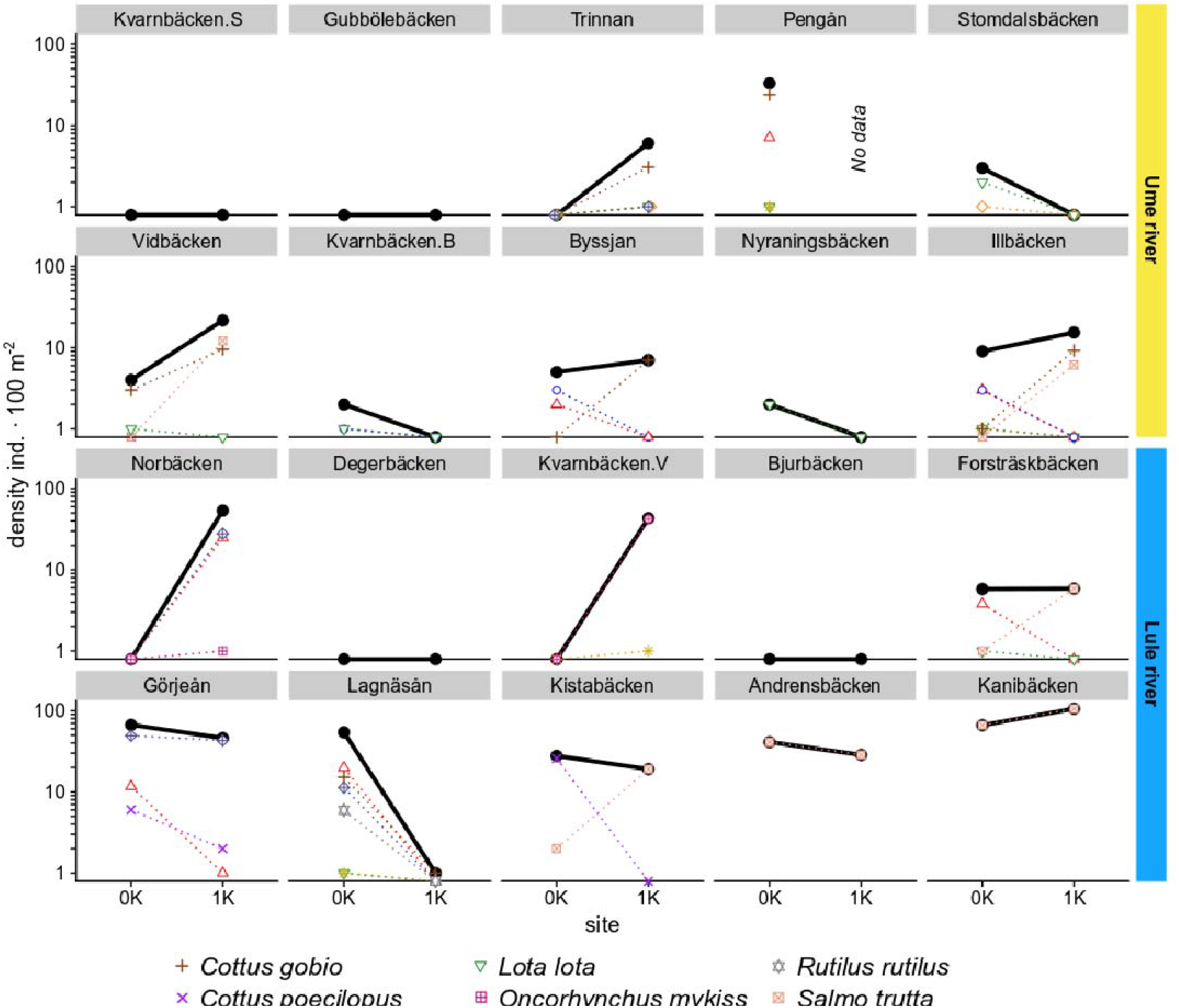
Densities of fish in the surveyed tributaries, at the tributary mouth (0K) and 1 km upstream the mouth (1K). Black symbols show total fish density and colored symbols show each individual species caught. Species with no catches at either site in a given tributary are not shown. Note that the y-axes are logarithmic.

## REFERENCES

Allan, J.D., Castillo, M.M., 2007. Stream ecology – structure and function of running waters, second edition. Springer, Dordrecht. https://doi.org/10.1007/978-1-4020-5583-6

Anderson, D., Moggridge, H., Warren, P., Shucksmith, J., 2015. The impacts of ‘run-of-river’ hydropower on the physical and ecological condition of rivers. Water Environm. J. 29, 268–276. https://doi.org/10.1111/wej.12101

Artdatabanken, 2020. Rödlistade arter i Sverige 2020. Artdatabanken, Swedish University of Agricultural Sciences, Uppsala. https://www.artdatabanken.se/publikationer/bestall-publikationer/bestall-rodlista-2020/

Barnett, A.G., van der Pols, J.C., Dobson, A.J. 2005. Regression to the mean: what it is and how to deal with it. Int. J. Epidemiol. 34, 215–220. https://doi.org/10.1093/ije/dyh299

Bartoń, K., 2020. MuMIn: Multi-Model Inference. R-package. https://CRAN.R-project.org/package=MuMIn.

Bates, D., Maechler, M., Bolker, B., Walker, S., Christensen, R.H.B., 2020. lme4: linear mixed models using ‘Eigen’ and S4. R package, version 1.1-26, https://cran.r-project.org/web/packages/lme4/index.html

Baxter, R.M., 1977. Environmental effects of dams and impoundments. Annu. Rev. Ecol. Syst. 8, 255–283.

Benda, L., Poff, N.L., Miller, D., Dunne, T., Reeves, G., Pess, G., Pollock, M., 2004. The network dynamics hypothesis: how channel networks structure riverine habitats. BioScience 54, 413–427. https://doi.org/10.1641/0006-3568(2004)054[0413:TNDHHC]2.0.CO;2

Bergquist, B., Degerman, E., Petersson, E., Sers, B., Stridsman, S., Winberg, S., 2014. Standardiserat elfiske I vattendrag – en manual med praktiska råd. Aqua reports 2014:15, 165 p. Swedish University of Agricultural Sciences, Drottningholm.

Bergstrand, M., Asp, S.-S., Lindström, G. 2014. Nationwide hydrological statistics for Sweden with high resolution using the hydrological model S-HYPE. Hydrol. Res. 45, 349–356. https://doi.org/10.2166/nh.2013.010

Betschta, R.L., Platts, W.S., 1986. Morphological features of small streams: significance and function. Water Resour. Bull. 22, 369–379. https://doi.org/10.1111/j.1752-1688.1986.tb01891.x

Cid, N., Erős, T., Heino, J., Singer, G., Jähnig, S.C., Cañedo-Argüelles, M., Bonada, N., Sarremejane, R., Mykrä, H., Sandin, L., Paloniemi, R., Varumo, L., Datry, T., 2022. From meta-system theory to the sustainable management of rivers in the Anthropocene. Front. Ecol. Environm. 20, 49–57. https://doi.org/10.1002/fee.2417

CEC (The Council of the European Communities), 1992. Council Directive 92/43/EEC of 21 May 1992 on the conservation of natural habitats and of wild fauna and flora. Official Journal of the European Communities L 206/7.

Clarke, K.R., 1993. Non-parametric multivariate analyses of changes in community structure. Aust. J. Ecol. 18, 117–143. https://doi.org/10.1111/j.1442-9993.1993.tb00438.x

Cooke, S.J., Paukert, C., Hogan, Z., 2012. Endangered river fish: factors hindering conservation and restoration. Endangered Species Res. 17, 179–191. https://doi.org/10.3354/esr00426

Copp, G.H., Jurajda, P., 1999. Size-structured diel use of river banks by fish. Aquat. Sci. 61:75–91. https://doi.org/10.1007/s000270050053

Czeglédi, I., Sály, P., Takács, P., Dolezsai, A., Nagy, S.A., Erős, T., 2016. The scales of variability of stream fish assemblages at tributary confluences. Aquat. Sci. 78, 641–654. https://doi.org/10.1007/s00027-015-0454-z

Degerman, E., Näslund, I., 2021. Fysisk restaurering av akvatiska miljöer. GRIP on LIFE:s rapportserie 2021:03, 381 p.

Dugdale, S.J., Bergeron, N.E., St-Hilaire, A., 2013. Temporal variability of thermal refuges and water temperature patterns in an Atlantic salmon river. Remote Sens. Environm. 136, 358–373. https://doi.org/10.1016/j.rse.2013.05.018

Dunn, C.G., Paukert, C.P., 2021. Accounting for dispersal and local habitat when evaluating tributary use by riverine fishes. Ecosphere 12, e03711. https://doi.org/10.1002/ecs2.3711

Dynesius, M., Nilsson, C., 1994. Fragmentation and flow regulation of river systems in the northern third of the world. Science 266, 753–762. https://doi.org/10.1126/science.266.5186.753

EC (European Commission), 2000. Directive 2000/60/EC of the European Parliament and of the Council of 23 October 2000 establishing a framework for community action in the field of water policy. Official Journal of the European Communities 43:L 327/1.

EC (European Commission), 2020. EU Biodiversity Strategy for 2030. Communication from the Commission to the European Parliament, the Council, the European Economic Social Committee and the Committee of the Regions COM/2020/380.

Ekman, S., 1922. Fiskarnas geografiska utbredning. In: Nordqvist O. (Ed.), Sötvattensfiske och fiskodling. Albert Bonniers Förlag, Stockholm, pp. 171–194.

Erkinaro, J., Erkinaro, H., Niemelä, E., 2017. Road culvert restoration expands the habitat connectivity and production area of juvenile Atlantic salmon in a large subarctic river system. Fish. Managem. Ecol. 24, 73–81. https://doi.org/10.1111/fme.12203

FAME Consortium, 2004. Manual for the application of the European Fish Index – EFI. A fish-based method to assess the ecological status of European rivers in support of the Water Framework Directive. Version 1.1. Ministrie van de Vlaamse Gemeenschap, Brussels.

Fausch, K.D., Torgersen, C.E., Baxter, C.V., Li, H.W., 2002. Landscapes to riverscapes: bridging the gap between research and conservation of stream fishes. BioScience 52, 483–498. https://doi.org/10.1641/0006-3568(2002)052[0483:LTRBTG]2.0.CO;2

Gann, G.D., McDonald, T., Walder, B., Aronson, J., Nelson, C.R., Jonson, J., Hallett, J.G., Eisenberg, C., Guariguata, M.R., Liu, J., Hua, F., Echeverría, C., Gonzales, E., Shaw, N., Decleer, K., Dixon, K.W., 2019. International principles and standards for the practice of ecological restoration. Second edition. Restor. Ecol. 27, S1–S46.

GBIF, 2020. GBIF – the Global Biodiversity Information Facility. GBIF Sectretariat, Copenhagen. https://www.gbif.org/ [Accessed: 2020-11-05]

GOS (Government Offices of Sweden), 2020. Nationell plan för moderna miljövillkor. Regeringsbeslut M2019/01769/Nm. Regeringen, Miljödepartementet, Stockholm.

Göthe, E., Degerman, E., Sandin, L., Segersten, J., Tamario, C., Mckie, B.G., 2019. Flow restoration and the impacts of multiple stressors on fish communities in regulated rivers. J. Appl. Ecol. 56, 1687–1702.

Grill, G., Lehner, B., Thieme, M., Geenen, B., Tickner, D., Antonelli, F., Babu, S., Borrelli, P., Cheng, L., Crochetiere, H., Ehalt Macedo, H., Filgueiras, R., Gopichot, M., Higgins, J., Hogan, Z., Lip, B., McClain, M.E., Meng, J., Mulligan, M., Nilsson, C., Olden, J.D., Opperman, J.J., Petry, P., Reidy Liermann, C., Sáenz, L., Salinas-Rodríguez, S., Schelle, P., Schmitt, R.J.P., Snider, J., Tan, F., Tockner, K., Valdujo, P.H., van Soesbergen, A., Zarfl, C., 2019. Mapping the world’s free-flowing rivers. Nature 569, 215–221.

Greenwood, M.T., Bickerton, M.A., Gurnell, A.M., Petts, G.E., 1999. Channel changes and invertebrate faunas below Nant-Y-Môch Dam, River Rheidol, Wales, UK: 35 years on. Regul. Rivers Res. Managem. 15, 99–112. https://doi.org/10.1002/(SICI)1099-1646(199901/06)15:1/3<99::AID-RRR530>3.0.CO;2-I

Grenoulliet, G., Pont, D., Hérissé, C., 2004. Within-basin fish assemblage structure: the relative influence of habitat versus stream spatial position on local species richness. Can. J. Fish. Aquat. Sci. 61, 93–102. https://doi.org//10.1139/F03-145

Harrison, X.A., 2014. Using observation-level random effects to model overdispersion in count data in ecology and evolution. PeerJ 2, e616. https://doi.org/10.7717/peerj.616

Hartig, F., 2021. DHARMa: Residual Diagnostics for Hierarchical (Multi-Level/Mixed) Regression Models. R-package, version 0.4.3. https://cran.r-project.org/web/packages/DHARMa/

Hesthagen, T., Saksgård, R., Hegge, O., Dervo, B.K., Skurdal, J., 2004. Niche overlap between young brown trout (Salmo trutta) and Siberian sculpin (Cottus poecilopus) in a subalpine Norwegian river. Hydrobiologia 521, 117–125. https://doi.org/10.1023/B:HYDR.0000026354.22430.17

IUCN (International Union for Conservation of Nature), 2020. IUCN Red List of Threatened Species. Version 2020-2. IUCN, Cambridge. https://www.iucnredlist.org/

Jackson, D.A., Peres-Neto, P.R., Olden, J.D., 2001. What controls who is where in freshwater fish communities – the roles of biotic, abiotic, and spatial factors. Can. J. Fish. Aquat. Sci. 58, 157–170. https://doi.org/10.1139/f00-239

Jansson, R., Nilsson, C., Malmqvist, B., 2007. Restoring freshwater ecosystems in riverine landscapes: the roles of connectivity and recovery processes. Freshw. Biol. 52, 589–596. https://doi.org/10.1111/j.1365-2427.2007.01737.x

Kiffney, P.M., Greene, C.M., Hall, J.E., Davies, J.R., 2006. Tributary streams create spatial discontinuities in habitat, biological productivity, and diversity in mainstem rivers. Can. J. Fish. Aquat. Sci. 63, 2518–2530. https://doi.org/10.1139/f06-138

Koizumi, I., Kanazawa, Y., Tanaka, Y., 2013. The fishermen were right: experimental evidence for tributary refuge hypothesis during floods. Zool. Sci. 30, 375–379.

Kullander, S.O., Nyman, L., Jilg, K., Delling, B., 2012. Nationalnyckeln till Sveriges flora och fauna. Strålfeniga fiskar. Actinopterygii. Artdatabanken, Swedish University of Agricultural Sciences, Uppsala.

Lane, E.W., 1955. The importance of fluvial morphology in hydraulic engineering. Proc. Am. Soc. Civ. Eng. 81, 1–17.

Laub, B.G., Thiede, G.P., Macfarlane, W.W., Budy, P., 2018. Evaluating the conservation potential of tributaries for native fishes in the upper Colorado River basin. Fisheries 43, 194–206. https://doi.org/10.1002/fsh.10054

Lenth, R.V., 2021. emmeans: estimated marginal means, aka least-squares means. R package, version 1.5.5-1, https://cran.r-project.org/web/packages/emmeans/index.html

Ligon, F.K., Dietrich, W.E., Trush, W.J., 1995. Downstream ecological effects of dams. BioScience 45, 183–192. https://doi.org/10.2307/1312557

Lipsey, T.S.B., Hubert, W.A., Rahel, F.J., 2005. Relationships of elevation, channel slope, and stream width to occurrences of native fishes at the Great Plains-Rocky Mountains interface. J. Freshw. Ecol. 20, 695–705.

Lundberg, R., 1899. Om svenska insjöfiskarnas utbredning (On the distribution of Swedish freshwaterfishes). Meddelanden från Kongl. Landtbruksakademien 58, 1–87.

Maceda-Veiga A, Baselga A, Sousa R, Vilà M, Doadrio I, de Sostoa A. 2017. Fine-scale determinants of conservation value of river reaches in a hotspot of native and non-native species diversity. Sci. Tot. Environm. 574, 455–466. https://doi.org/10.1016/j.scitotenv.2016.09.097

Malm Renöfält, B., Jansson, R., Nilsson, C., 2010. Effects of hydropower generation and opportunities for environmental flow management in Swedish riverine ecosystems. Freshw. Biol. 55, 49–67. https://doi.org/10.1111/j.1365-2427.2009.02241.x

Meyer, J.L., Strayer, D.L., Wallace, J.B., Eggert, S.L., Helfman, G.S., Leonard, N.E., 2007. The contribution of headwater streams to biodiversity in river networks. J. Am. Water Res. Assoc. 43, 86–103.

Milner, V.S., Yarnell, S.M., Peek, R.A., 2019. The ecological importance of unregulated tributaries to macroinvertebrate diversity and community composition in a regulated river. Hydrobiologia 829, 291–305. https://doi.org/10.1007/s10750-018-3840-4

Miranda, L.E., Killgore, K.J., Slack, W.T., 2019. Spatial organization of fish diversity in a species-rich basin. River Res. Appl. 35:188–196. https://doi.org/10.1002/rra.3392

Nunn, A.D., Copp, G.H., Vilizzi, L., Carter, M.G., 2010. Seasonal and diel patterns in the migrations of fishes between a river and a floodplain tributary. Ecol. Freshw. Fish 19, 153–162.

Oberdorff, T., Tedesco, P.A., Hugueny, B., Leprieur, F., Beauchard, O., Brosse, S., Dürr, H.H., 2011. Global and regional patterns in riverine fish species richness: a review. Int. J. Ecol. 2011, 967631. https://doi.org/10.1155/2011/967631

Ödmann, E., Bucht, E., Nordström, M., 1982. Naturskyddet och utnyttjandet av vattnet. In: Ödmann, E., Bucht, E., Nordström, M. (Eds.), Vildmarken och välfärden, pp. 66–78. Liberförlag, Lund.

Ogle, D., Wheeler, P., Dinno, A., 2020. Package ‘FSA’ – Simple fisheries stock assessment methods’, version 0.8.31. https://github.com/droglenc/FSA

Oksanen, J., Blanchet, F.G., Friendly, M., Kindt, R., Legendre, P., McGlinn, D., Minchin, P.R., O’Hara, R.B., Simpson, G.L., Solymos, P., Stevens, M.H.H., Szoecs, E., Wagner, H., 2019. Package ‘vegan’ – Community ecology package, version 2.5-6. https://github.com/vegandevs/vegan

Pielou, E.C., 1966. The measurement of diversity in different types of biological collections. J. Theor. Biol. 13, 131–144. https://doi.org/10.1016/0022-5193(66)90013-0

Poff, N.L., Olden, J.D., Merritt, D.M., Pepin, D.M., 2007. Homogenization of regional river dynamics by dams and global biodiversity implications. Proc. Natl. Acad. Sci. 104, 5732–5737. https://doi.org/10.1073/pnas.0609812104

Pont, D., Hugueny, B., Beier, U., Goffaux, D., Melcher, A., Noble, R., Rogers, C., Roset, N., Schmutz. S., 2006. Assessing river biotic condition at a continental scale: a European approach using functional metrics and fish assemblages. J. Appl. Ecol. 43, 70–80. https://doi.org/10.1111/j.1365-2664.2005.01126.x

Power, M.E., Dietrich, W.E., 2002. Food webs in river networks. Ecol. Res. 17, 451–471. https://doi.org/10.1046/j.1440-1703.2002.00503.x

Pracheil, B.M., McIntyre, P.B., Lyons, J.D., 2013. Enhancing conservation of large-river biodiversity by accounting for tributaries. Front. Ecol. Environm. 11, 124–128. https://doi.org/10.1890/120179.

QGIS Development Team, 2021. QGIS Geographic Information System. Open Source Geospatial Foundation Project. https://qgis.org/

Rex, W., Foster, V., Lyon, K., Bucknall, J., Liden, R., 2014. Supporting Hydropower: An Overview of the World Bank Group’s Engagement. World Bank Group, Washington D.C. http://hdl.handle.net/10986/20351

Reynolds, L., Herlihy, A.T., Kaufmann, P.R., Gregory, S.V., Hughes, R.M., 2003. Electrofishing effort requirements for assessing species richness and biotic integrity in western Oregon streams. N. Am. J. Fish. Managem. 23, 450–461. https://doi.org/10.1577/1548-8675(2003)023<0450:EERFAS>2.0.CO;2

Rice, S.P., Ferguson, R.I., Hoey, T.B., 2006. Tributary control of physical heterogeneity and biological diversity at river confluences. Can. J. Fish. Aquat. Sci. 63, 2553–2566. https://doi.org/10.1139/f06-145

Rice, S.P., Kiffney, P., Greene, C., Pess, G.R., 2008. The ecological importance of tributaries and confluences. In: Rice, S.P., Roy, A.G., Rhoads, B.L. (Eds.), River confluences, tributaries and the fluvial network. John Wiley & Sons Inc., Hoboken, pp. 209–242.

Salas, A.K., Snyder, E.B., 2010. Diel fish habitat selection in a tributary stream. Am. Midl. Nat. 163, 33–43.

Sandin, L., Degerman, E., Bergengren, J., Gren, I.-M., Carlson, P., Donadi, S., Andersson, M., Drakare, S., Göthe, E., Johnson, R.K., Kahlert, M., Segersten, J., McKie, B., Spjut, D., Tirkaso, W.T., Tamario, C., Trigal, C., von Wachtenfeldt, E., 2017. Ekologiska och ekonomiska strategier för optimering av vattenkraftsrelaterade miljöåtgärder, EKOLIV. Energiforsk rapport 2017:450. Energiforsk, Stockholm, 156 p.

Schäfer, T., 2021. Legal protection schemes for free-flowing rivers in Europe: an overview. Sustainability 13:6423. https://doi.org/10.3390/su13116423

Schmidt-Kloiber, A., Hering, D., 2015. http://www.freshwaterecology.info - an online tool that unifies, standardises and codifies more than 20,000 European freshwater organisms and their ecological preferences. Ecol. Ind. 53, 271–282. https://doi.org/10.1016/j.ecolind.2015.02.007

Scott, W.B., Crossman, E.J. 1973. Freshwater fishes of Canada. Bull. Fish. Res. Board Can. 184, 1–966. https://publications.gc.ca/site/eng/9.870340/publication.html

Seber, G.A.F., 1982. The estimation of animal abundance and related methods, 2nd edition. Macmillan Publishing Co., Inc., New York.

SEA (Swedish Energy Agency), 2021. Energy in Sweden 2021 – an overview. ET 2021:11. Swedish Energy Agency, Stockholm, 18 p.

SERS, 2020. Swedish Electrofishing RegiSter – SERS. Swedish University of Agricultural Sciences (SLU), Department of Aquatic Resources, Drottningholm. http://www.slu.se/elfiskeregistret. [2020-10-14]

Shannon, C.E., 1948. A mathematical theory of communication. Bell Syst. Techn. J. 27, 623–656. https://doi.org/10.1002/j.1538-7305.1948.tb01338.x

SMHI (Swedish Meteorological and Hydrological Institute), 2010. Sveriges vattendrag. SMHI Faktablad 44-2010, 1–4. SMHI, Norrköping.

Soukup, P.R., Näslund, J., Höjesjö, J., Boukal, D.S., 2022. From individuals to communities: habitat complexity affects all levels of organization in aquatic environments. WIREs Water 9, e1575. https://doi.org/10.1002/wat2.1575

Stapanian, M.A., Paragamian, V.L., Madenjian, C.P., Jackson, J.R., Lappalainen, J., Evenson, M.J., Neufeld, M.D., 2010. Worldwide status of burbot and conservation measures. Fish Fish. 11, 34–56. https://doi.org/10.1111/j.1467-2979.2009.00340.x

Trigal, C., Degerman, E., 2015. Multiple factors and thresholds explaining fish species distributions in lowland streams. Global Ecol. Conserv. 4, 589–601. https://doi.org/10.1016/j.gecco.2015.10.009

Thornbrugh, D.J., Gido, K.B., 2009. Influence of spatial positioning within stream networks on fish assemblage structure in the Kansas River basin, USA. Can. J. Fish. Aquat. Sci. 67, 143–156. https://doi.org/10.1139/F09-169

Vinson, M.R., 2001. Long-term dynamics of an invertebrate assemblage downstream from a large dam. Ecol. Appl. 11, 711–730. https://doi.org/10.1890/1051-0761(2001)011[0711:LTDOAI]2.0.CO;2

Ver Hoef, J.M., Boveng, P.L., 2007. Quasi-Poisson vs. negative binomial regression: how should we model overdispersed count data? Ecology 88, 2766–2772.

Vörösmarty, C.J., McIntyre, P.B., Gessner, M.O., Dudgeon, D., Prusevich, A., Green, P., Glidden, S., Bunn, S.E., Sullivan, C.A., Reidy Liermann, C., Davies, P.M., 2010. Global threats to human water security and river biodiversity. Nature 465, 555–561. https://doi.org/10.1038/nature09440

Wang, T., Kelson, S.J., Greer, G., Thompson, S.E., Carlson, S.M., 2020. Tributary confluences are dynamic thermal refuges for a juvenile salmonid in a warming river network. River Res. Appl. 36, 1076–1086.

Wickham, H., Averick, M., Bryan, J., Chang, W., McGowan, L.D., François, R., Grolemund, G., Hayes, A., Henry, L., Hester, J., Kuhn, M., Pedersen, T.L., Miller, E., Bache, S.M., Müller, K., Ooms, J., Robinson, D., Seidel, D.P., Spinu, V., Takahashi, K., Vaughan, D., Wilke, C., Woo, K., Yutani, H., 2019. Welcome to the Tidyverse. J. Open Source Software 4, 1686.

Widén, Å., Jansson, R., Johansson, M., Lindström, M., Sandin, L., Wisaeus, D., 2016. Maximal ekologisk potential i Umeälven. Final project report for Projekt Umeälven, https://umealven.se/rapporter/

Widén, Å., Malm Renöfält, B., Degerman, E., Wisaeus, D., Jansson, R., 2021. Let it flow: modeling ecological benefits and hydropower production impacts of banning zero-flow events in a large regulated river system. Sci. Tot. Environm. 783, 147010.

Wipfli, M., Gregovich, D.P., 2002. Export of invertebrates and detritus from fishless headwater streams in southeastern Alaska: implications for downstream salmonid production. Freshw. Biol. 47, 957–969. https://doi.org/10.1046/j.1365-2427.2002.00826.x

Wohl, E., 2017. The significance of small streams. Front. Earth Sci. 11, 447–456. https://doi.org/10.1007/s11707-017-0647-y

Ziv, G., Baran, E., Nam, S., Rodriguez-Iturbe, I., 2012. Trading-off fish biodiversity, food security, and hydropower in the Mekong River Basin. Proc. Natl. Acad. Sci. 109, 5609–5614. https://doi.org/10.1073/pnas.1201423109

