## Supplementary information: analyses for "Sediment substrate size influences fish diversity in tributary mouth areas in impounded boreal rivers in Sweden"

**Table S1.** Model reduction procedure for analysis of number of species as dependent on aggradation status in the tributary mouth area.

| Number of species (no.spp) | | |
| --- | --- | --- |
| **Global Poisson-GLMM model**: no.spp ~ river + aggr + log_10_(fi.ar) + (1\|imp) | | |
| *Step 1: compare all subordinate GLMMs using the dredge()-command from the MuMIn-package; factor* aggr = *fixed (i.e. cannot be removed)* | | |
| **Models (ranked by AICc)** | **AICc** | **ΔAICc** |
| M1: no.spp ~ aggr + (1\|imp) | 76.0 | 0 |
| M2: no.spp ~ aggr + log_10_(fi.ar) + (1\|imp) | 77.5 | 1.5 |
| *No other models with ΔAICc ≤ 2* | | |
| *Step 2: select M2 (most complex within ΔAICc ≤ 2) for comparison with GLM without* (1\| imp) | | |
| **Models** | **AICc** | **ΔAICc** |
| M2: no.spp ~ aggr + log_10_(fi.ar) + (1\|imp) | 77.5 | 0 |
| GLM: no.spp ~ aggr + log_10_(fi.ar) | 78.7 | 1.1 |
| ***Decision: use M2 for analysis*** | | |

**Table S2.** Model reduction procedure for analysis of Shannon diversity *H*’ as dependent on aggradation status in the tributary mouth area.

| Shannon diversity *H*’ (shannon) | | |
| --- | --- | --- |
| **Global LMM model**: log_10_(shannon+1) ~ river + aggr + log_10_(fi.ar) + (1\|imp) | | |
| *Step 1: compare all subordinate LMMs using the dredge()-command from the MuMIn-package; factor* aggr = *fixed (i.e. cannot be removed)* | | |
| **Models (ranked by AICc)** | **AICc** | **ΔAICc** |
| M1: ~ aggr + (1\|imp) | -10.2 | 0 |
| *No other models with ΔAICc ≤ 2* | | |
| *Step 2: select M1 (only model within ΔAICc ≤ 2) for comparison with LM without* (1\| imp) | | |
| **Models** | **AICc** | **ΔAICc** |
| M2: ~ aggr + (1\|imp) | -10.2 | 0 |
| LM: ~ aggr | -15.4 | -5.2 |
| ***Decision: use LM for analysis*** | | |

**Table S3.** Model reduction procedure for analysis of number of species as dependent on NMDS-axes describing the environment in the tributary mouth area.

| Number of species (no.spp) | | |
| --- | --- | --- |
| **Global Poisson-GLMM model**: no.spp ~ poly(nmds1, 2) × poly(nmds2, 2) + river + log_10_(fi.ar) + (1\|imp) | | |
| *Step 1: compare all subordinate GLMMs using the* dredge()*-command from the* MuMIn*-package* | | |
| **Models (ranked by AICc)** | **AICc** | **ΔAICc** |
| M1: no.spp ~ poly(nmds1, 2) + log_10_(fi.ar) + (1\|imp) | 68.1 | 0 |
| M2: no.spp ~ poly(nmds1, 2) + (1\|imp) | 69.6 | 1.5 |
| *No other models with ΔAICc ≤ 2* | | |
| *Step 2: check AICc of models without quadratic terms of* nmds1 and nmds2 | | |
| *All these models have AICc > 71* | | |
| *Step 3: select M1 (most complex within ΔAICc ≤ 2) for comparison with GLMs without* (1\| imp) | | |
| **Models** | **AICc** | **ΔAICc** |
| M1: no.spp ~ poly(nmds1, 2) + log_10_(fi.ar) + (1\|imp) | 68.1 | 0 |
| GLM1: no.spp ~ poly(nmds1, 2) + log_10_(fi.ar) | 64.3 | -3.8 |
| GLM2: no.spp ~ poly(nmds1, 2) | 66.3 | -1.8 |
| ***Decision: use GLM1 for analysis*** | | |

**Table S4.** Model reduction procedure for analysis of Shannon diversity *H*’ as dependent on NMDS-axes describing the environment in the tributary mouth area.

| Number of species (no.spp) | | |
| --- | --- | --- |
| **Global LMM model**: log_10_(shannon+1) ~ poly(nmds1, 2) × poly(nmds2, 2) + river + log_10_(fi.ar) + (1\|imp) | | |
| *Step 1: compare all subordinate LMMs using the* dredge()*-command from the* MuMIn*-package* | | |
| **Models (ranked by AICc)** | **AICc** | **ΔAICc** |
| M1: log_10_(shannon+1) ~ (1\|imp) | -14.5 | 0 |
| *No other models with ΔAICc ≤ 2* | | |
| *Step 2: check AICc of models without quadratic term of* nmds1 and nmds2 | | |
| *All these models have ΔAICc > 2, as compared to M1* | | |
| *Step 3: select M1 (only model within ΔAICc ≤ 2) for comparison with LM without* (1\| imp) | | |
| **Models** | **AICc** | **ΔAICc** |
| M1: log_10_(shannon+1) ~ (1\|imp) | -14.5 | 0 |
| LM1: log_10_(shannon+1) ~ poly(nmds1, 2) + log_10_(fi.ar) | -19.3 | -4.8 |
| *No other LM-models within ΔAICc < 2 from the model with the lowest AIC_c_-value* | | |
| ***Decision: use LM1 for analysis*** | | |

**Table S5.** Model reduction procedure for analysis of **total fish density** as dependent on NMDS-axes describing the environment in the tributary mouth area.

| Total fish density (all species combined) | | |
| --- | --- | --- |
| **Global LMM model**: log_10_(density+1) ~ poly(nmds1, 2) × poly(nmds2, 2) + river + (1\|imp) | | |
| *Step 1: compare all subordinate LMMs using the* dredge()*-command from the* MuMIn*-package.* | | |
| **Models (ranked by AICc)** | **AICc** | **ΔAICc** |
| M1: log_10_(density+1) ~ (1\|imp) | 39.3 | 0 |
| M2: log_10_(density+1) ~ river + (1\|imp) | 41.1 | 1.85 |
| *No other models with ΔAICc ≤ 2* | | |
| *Step 2: check AICc of models without quadratic term of* nmds1 and nmds2 | | |
| *All these models have ΔAICc > 2, as compared to M1* | | |
| *Step 3: select M2 (most complex model within ΔAICc ≤ 2) for comparison with LM without* (1\| imp) | | |
| **Models** | **AICc** | **ΔAICc** |
| M2: log_10_(density+1) ~ river + (1\|imp) | 41.1 | 0 |
| LM1: log_10_(density+1) ~ nmds1 | 41.0 | -0.1 |
| LM2: log_10_(density+1) ~ nmds1 + nmds2 | 42.5 | 1.4 |
| *No other LM-models within ΔAICc < 2 from the model with the lowest AIC_c_-value* | | |
| ***Decision: use LM2 for analysis*** | | |

**Table S6.** Model reduction procedure for analysis of density of **tolerant** species as dependent on NMDS-axes describing the environment in the tributary mouth area.

| Tolerant fish density | | |
| --- | --- | --- |
| **Global LMM model**: log_10_(density+1) ~ poly(nmds1, 2) × poly(nmds2, 2) + river + (1\|imp) | | |
| *Step 1: compare all subordinate LMMs using the* dredge()*-command from the* MuMIn*-package.* | | |
| **Models (ranked by AICc)** | **AICc** | **ΔAICc** |
| M1: log_10_(density+1) ~ (1\|imp) | 13.1 | 0 |
| *No other models with ΔAICc ≤ 2* | | |
| *Step 2: check AICc of models without quadratic term of* nmds1 and nmds2 | | |
| *All these models have ΔAICc > 2, as compared to M1* | | |
| *Step 3: select M1 (most complex model within ΔAICc ≤ 2) for comparison with LM without* (1\| imp) | | |
| **Models** | **AICc** | **ΔAICc** |
| M1: log_10_(density+1) ~ (1\|imp) | 13.1 | 0 |
| LM1: log_10_(density+1) ~ 1 | 6.6 | -6.5 |
| *No other LM-models within ΔAICc < 2 from the model with the lowest AIC_c_-value* | | |
| ***Decision: use LM1 for analysis*** | | |

**Table S7.** Model reduction procedure for analysis of density of **intolerant** species as dependent on NMDS-axes describing the environment in the tributary mouth area.

| Intolerant fish density | | |
| --- | --- | --- |
| **Global LMM model**: log_10_(density+1) ~ poly(nmds1, 2) × poly(nmds2, 2) + river + (1\|imp) | | |
| *Step 1: compare all subordinate LMMs using the* dredge()*-command from the* MuMIn*-package.* | | |
| **Models (ranked by AICc)** | **AICc** | **ΔAICc** |
| M1: log_10_(density+1) ~ (1\|imp) | 41.6 | 0 |
| M2: log_10_(density+1) ~ river + (1\|imp) | 43.1 | 1.5 |
| *No other models with ΔAICc ≤ 2* | | |
| *Step 2: check AICc of models without quadratic term of* nmds1 and nmds2 | | |
| *All these models have ΔAICc > 2, as compared to M1* | | |
| *Step 3: select M2 (most complex model within ΔAICc ≤ 2) for comparison with LM without* (1\| imp) | | |
| **Models** | **AICc** | **ΔAICc** |
| M2: log_10_(density+1) ~ river + (1\|imp) | 43.1 | 0 |
| LM1: log_10_(density+1) ~ poly(nmds1, 2) + river | 40.2 | -2.9 |
| *No other LM-models within ΔAICc < 2 from the model with the lowest AIC_c_-value* | | |
| ***Decision: use LM1 for analysis*** | | |

**Table S8.** Model reduction procedure for analysis of density of **benthic** species as dependent on NMDS-axes describing the environment in the tributary mouth area.

| Benthic fish density | | |
| --- | --- | --- |
| **Global LMM model**: log_10_(density+1) ~ poly(nmds1, 2) × poly(nmds2, 2) + river + (1\|imp) | | |
| *Step 1: compare all subordinate LMMs using the* dredge()*-command from the* MuMIn*-package.* | | |
| **Models (ranked by AICc)** | **AICc** | **ΔAICc** |
| M1: log_10_(density+1) ~ poly(nmds1, 2) + (1\|imp) | 34.5 | 0 |
| *No other models with ΔAICc ≤ 2* | | |
| *Step 2: check AICc of models without quadratic term of* nmds1 and nmds2 | | |
| *All these models have ΔAICc > 2, as compared to M1* | | |
| *Step 3: select M1 (most complex model within ΔAICc ≤ 2) for comparison with LM without* (1\| imp) | | |
| **Models** | **AICc** | **ΔAICc** |
| M1: log_10_(density+1) ~ poly(nmds1, 2) + (1\|imp) | 34.5 | 0 |
| LM1: log_10_(density+1) ~ poly(nmds1, 2) | 27.7 | -6.8 |
| *No other LM-models within ΔAICc < 2 from the model with the lowest AIC_c_-value* | | |
| ***Decision: use LM1 for analysis*** | | |

**Table S9.** Model reduction procedure for analysis of density of **rheophilic** species as dependent on NMDS-axes describing the environment in the tributary mouth area.

| Rheophilic fish density | | |
| --- | --- | --- |
| **Global LMM model**: log_10_(density+1) ~ poly(nmds1, 2) × poly(nmds2, 2) + river + (1\|imp) | | |
| *Step 1: compare all subordinate LMMs using the* dredge()*-command from the* MuMIn*-package.* | | |
| **Models (ranked by AICc)** | **AICc** | **ΔAICc** |
| M1: log_10_(density+1) ~ (1\|imp) | 41.3 | 0 |
| M2: log_10_(density+1) ~ river + (1\|imp) | 42.7 | 1.48 |
| *No other models with ΔAICc ≤ 2* | | |
| *Step 2: check AICc of models without quadratic term of* nmds1 and nmds2 | | |
| *All these models have ΔAICc > 2, as compared to M1* | | |
| *Step 3: select M2 (most complex model within ΔAICc ≤ 2) for comparison with LM without* (1\| imp) | | |
| **Models** | **AICc** | **ΔAICc** |
| M2: log_10_(density+1) ~ river + (1\|imp) | 42.7 | 0 |
| LM1: log_10_(density+1) ~ poly(nmds1, 2) + river | 39.7 | -3.0 |
| *No other LM-models within ΔAICc < 2 from the model with the lowest AIC_c_-value* | | |
| ***Decision: use LM1 for analysis*** | | |

**Table S10.** Model reduction procedure for analysis of density of **red-listed** species as dependent on NMDS-axes describing the environment in the tributary mouth area.

| Red-listed fish density | | |
| --- | --- | --- |
| **Global LMM model**: log_10_(density+1) ~ poly(nmds1, 2) × poly(nmds2, 2) + river + (1\|imp) | | |
| *Step 1: compare all subordinate LMMs using the* dredge()*-command from the* MuMIn*-package.* | | |
| **Models (ranked by AICc)** | **AICc** | **ΔAICc** |
| M1: log_10_(density+1) ~ (1\|imp) | 25.4 | 0 |
| M2: log_10_(density+1) ~ poly(nmds1, 2) +(1\|imp) | 27.2 | 1.8 |
| *No other models with ΔAICc ≤ 2* | | |
| *Step 2: check AICc of models without quadratic term of* nmds1 and nmds2 | | |
| *All these models have ΔAICc > 2, as compared to M1* | | |
| *Step 3: select M2 (most complex model within ΔAICc ≤ 2) for comparison with LM without* (1\| imp) | | |
| **Models** | **AICc** | **ΔAICc** |
| M2: log_10_(density+1) ~ poly(nmds1, 2) + (1\|imp) | 27.2 | 0 |
| LM1: log_10_(density+1) ~ poly(nmds1, 2) | 19.1 | -8.1 |
| LM2: log_10_(density+1) ~ 1 | 19.5 | -7.7 |
| LM3: log_10_(density+1) ~ nmds2 | 20.8 | -6.4 |
| *No other LM-models within ΔAICc < 2 from the model with the lowest AIC_c_-value* | | |
| ***Decision: use LM1 for analysis*** | | |

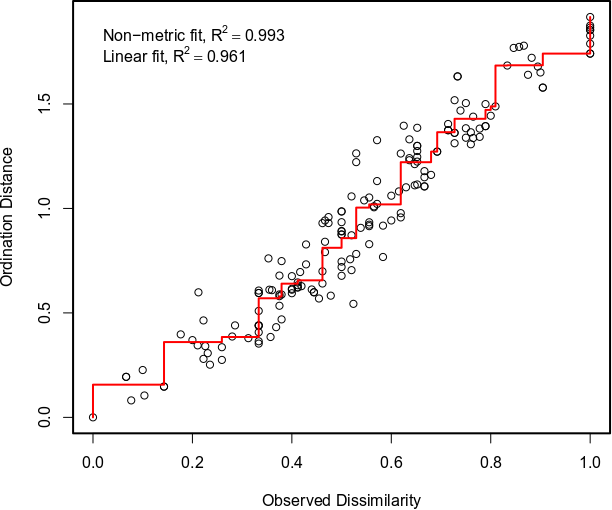

**Figure S1**. Shepard plot derived from the non-metric multidimensional scaling (NMDS) analysis.

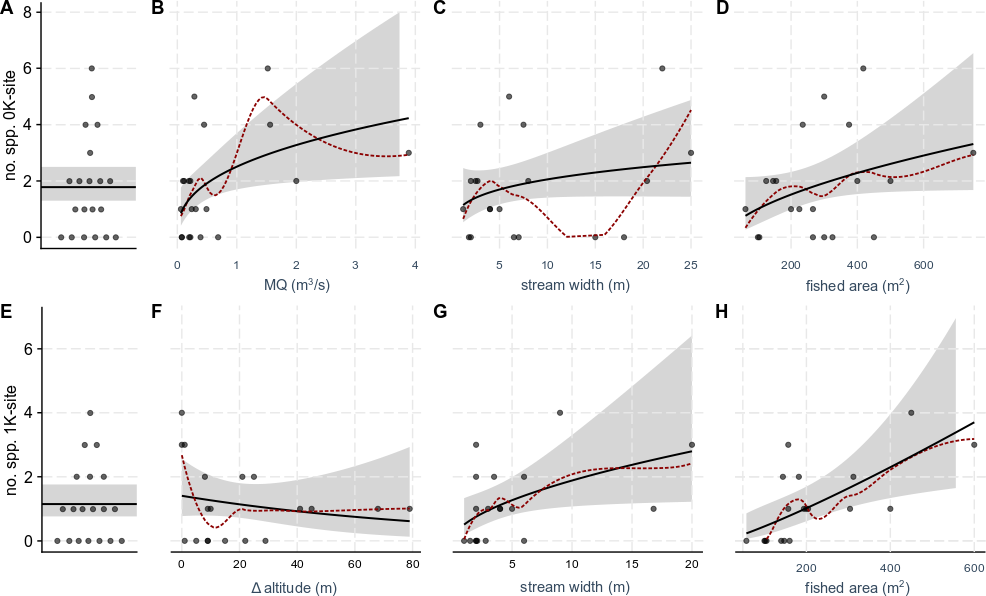

**Figure S2**. Effects of some different environmental variables related to overall size of the tributary. A) Number of species at the tributary mouth sites (0K-sites). B-D) Effects of B) mean annual discharge (MQ), C) stream channel width, and D) fished area on number of species caught at the 0K-sites. E) Number of species at the sites located 1 km upstream the tributary mouth (1K-sites). F-H) Effects of F) site altitude (in relation to the altitude of the mainstem; Δ altitude), G) stream channel width, and H) fished area on number of species caught at the 0K-sites. Black lines show the regression lines, surrounded by the 95% confidence interval/-band in grey. Red dashed lines show loess regression lines.

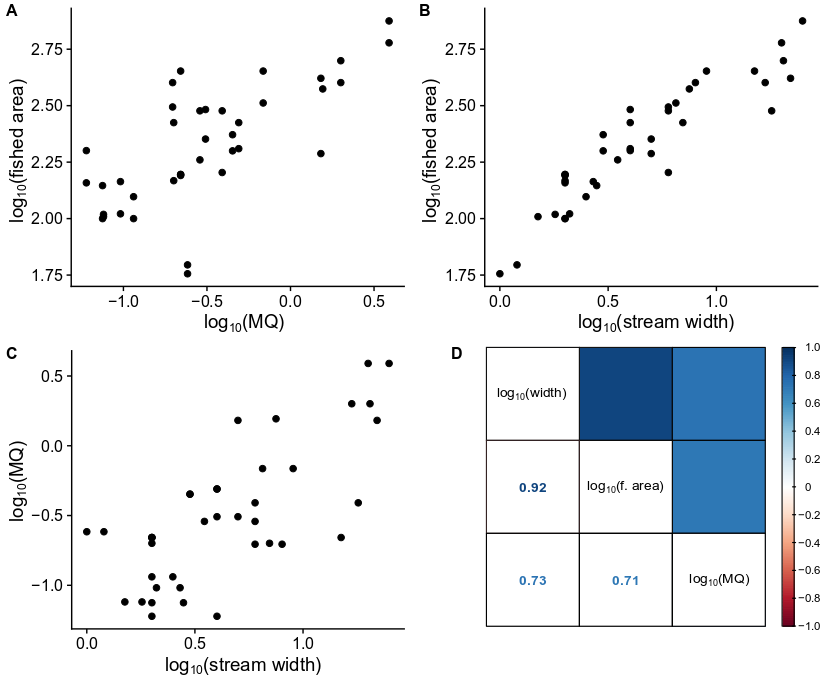

**Figure S3**. Correlations among the variables fished area (m^2^), mean annual discharge (MQ; m^3^ ∙ s^-1^), and stream channel width (m). A-C) Scatterplots showing the overall relationships. D) Correlation matrix with Pearson correlation coefficients (*r*).

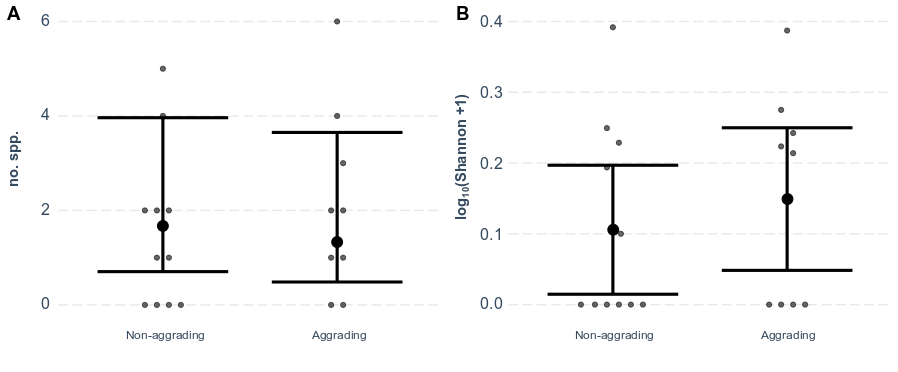

**Figure S4**. Effects of binary aggradation status in the tributary mouth area on A) number of species (no. spp.) and B) diversity as indicated by Shannon H’ (log transformed). Contrast ratio for A): non-aggrading:aggrading = 1.26 (SE: 0.566; *p* = 0.061). Contrast estimate for B): non-aggrading – aggrading = -0.043 (SE: 0.064; *p* = 0.51). Error bars show the 95% confidence intervals around the means.

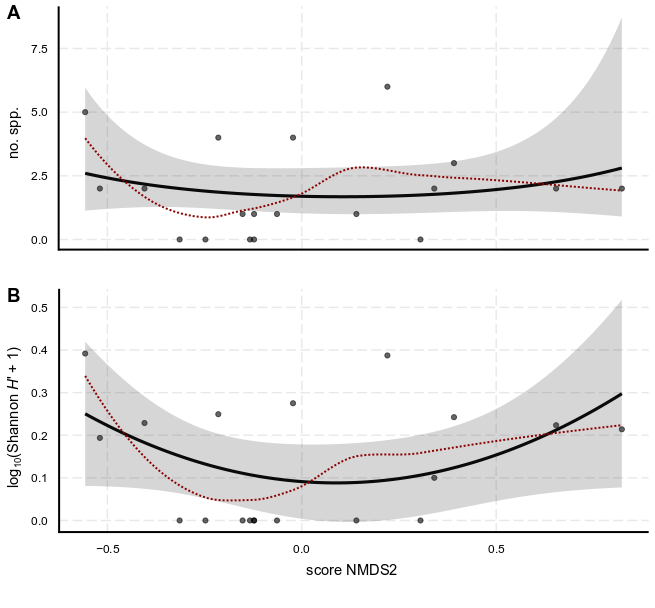

**Figure S5**. Effect of the second NMDS axis (NMDS2) scores on A) number of species caught and B) Shannon diversity *H*’. Black line show the modelled effect from A) a Poisson GLM including a polynomial of NMDS2 and the logarithm of fished area, with the model structure ~ poly(nmds2, 2) + log_10_(fi.ar), and B) a LM with the same factors as in A). Grey bands around the predicted lines (in black) show 95% confidence bands. Red dotted lines shows a loess regressions based on raw data.

Results, model in A) – poly(nmds2, 2): χ^2^ = 0.89, p = 0.64; log_10_(fi.ar): χ^2^ = 2.61, p = 0.11.

Results, model in B) – poly(nmds2, 2): F_2,15_ = 1.99, p = 0.34; log_10_(fi.ar): F_1,15_ = 2.11, p = 0.17.

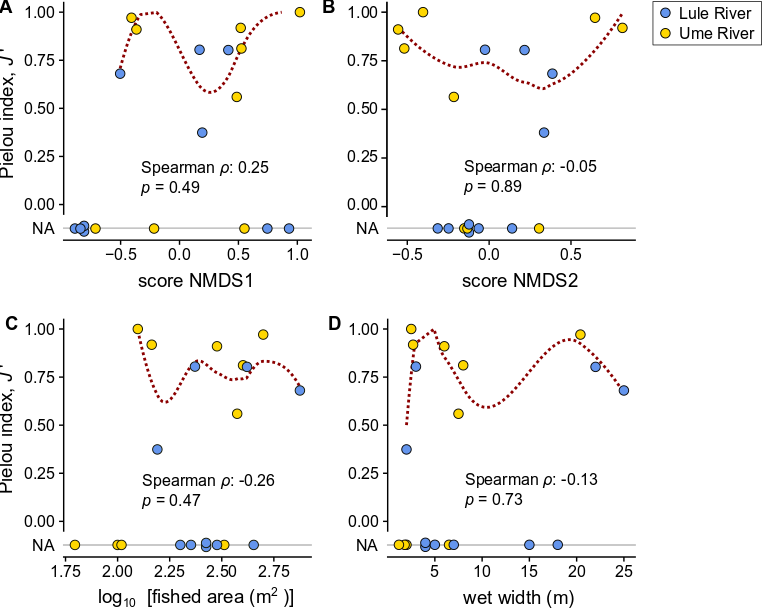

**Figure S6**. Visualization of the relationship between Pielou evenness (*J*’) and A) the first axis of the NMDS (NMDS1), B) the second axis of the NMDS (NMDS2), C) fished area, and D) stream width. Results from Spearman rank correlations are reported in each figure A-D. Red dotted lines shows a loess regressions based on raw data. Note the presence of ‘NA’ (missing values), which is due to the absence of species or the presence of only a single species (i.e. situations where Shannon *H*’ = 0).

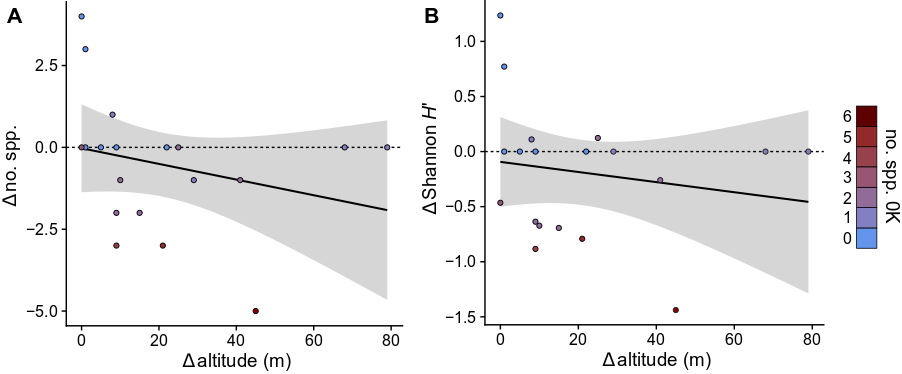

**Figure S7**. Relationship between difference in altitude and difference in A) species richness and B) Shannon diversity, for paired 0K- and 1K sites in the investigated tributaries. Color scale represents number of species at the 0K sites. Linear regression line is shown with 95% confidence bands. No significant correlation was detected.
