## Supplementary information: Photos of study sites for "Sediment substrate size influences fish diversity in tributary mouth areas in impounded boreal rivers in Sweden"

### Ume River: Kvarnbäcken.1

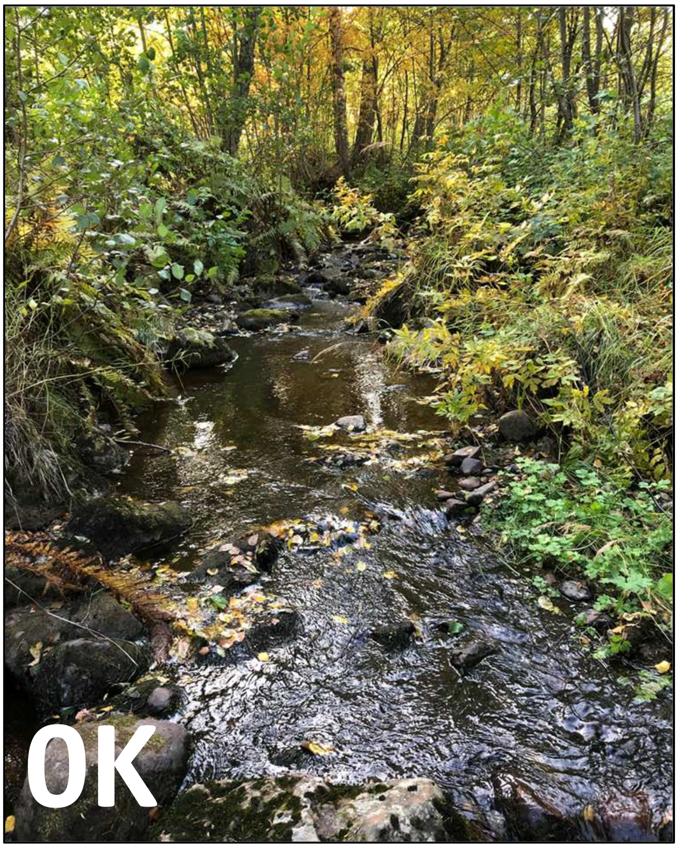


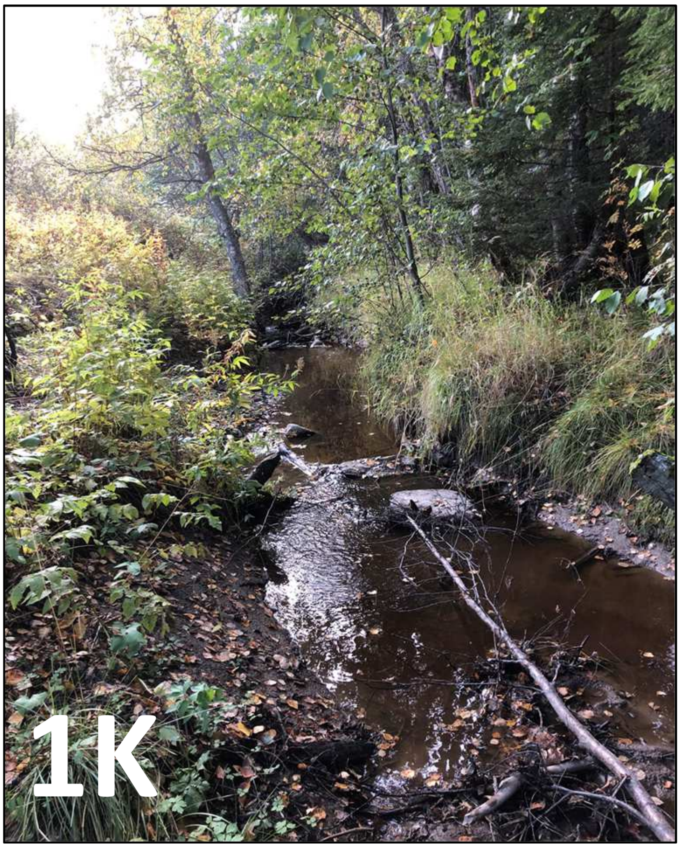

### Ume River: Gubbölebäcken

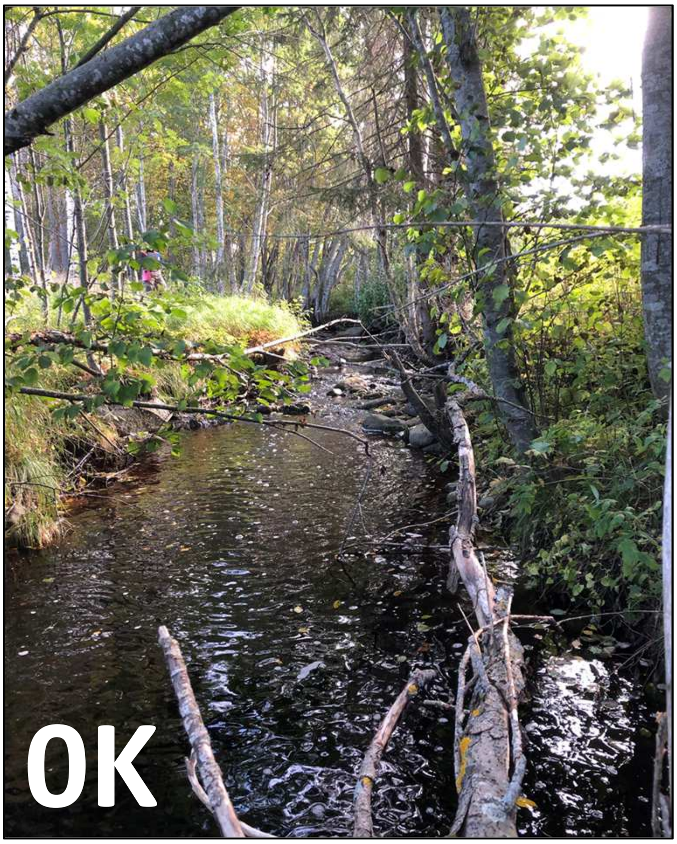


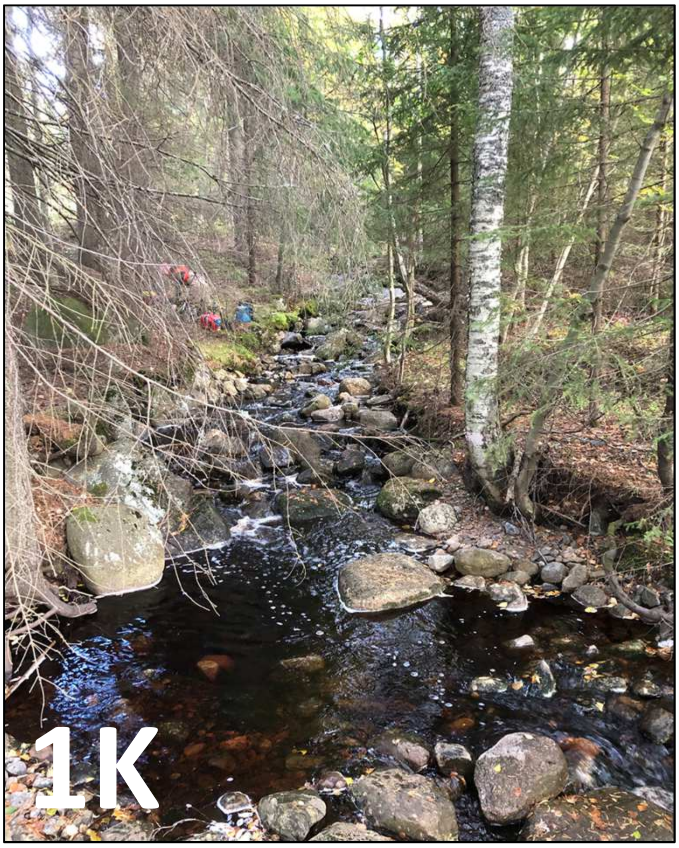

### Ume River: Trinnan

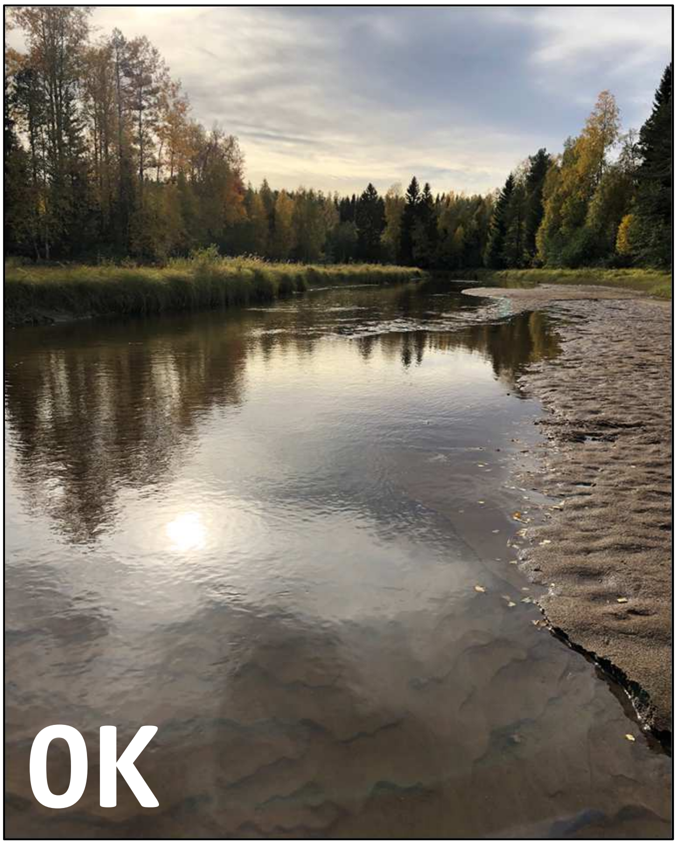


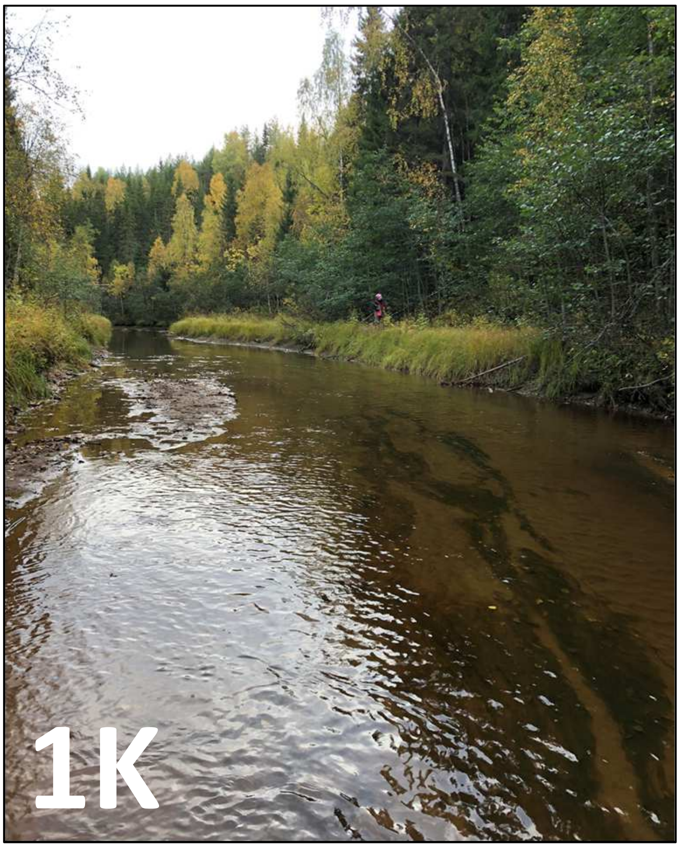

### Ume River: Pengån

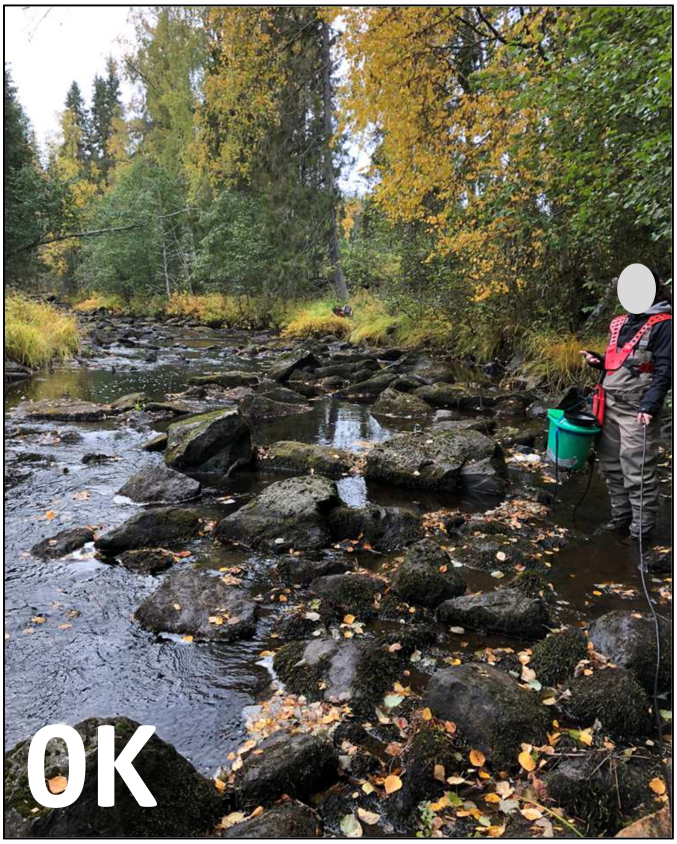


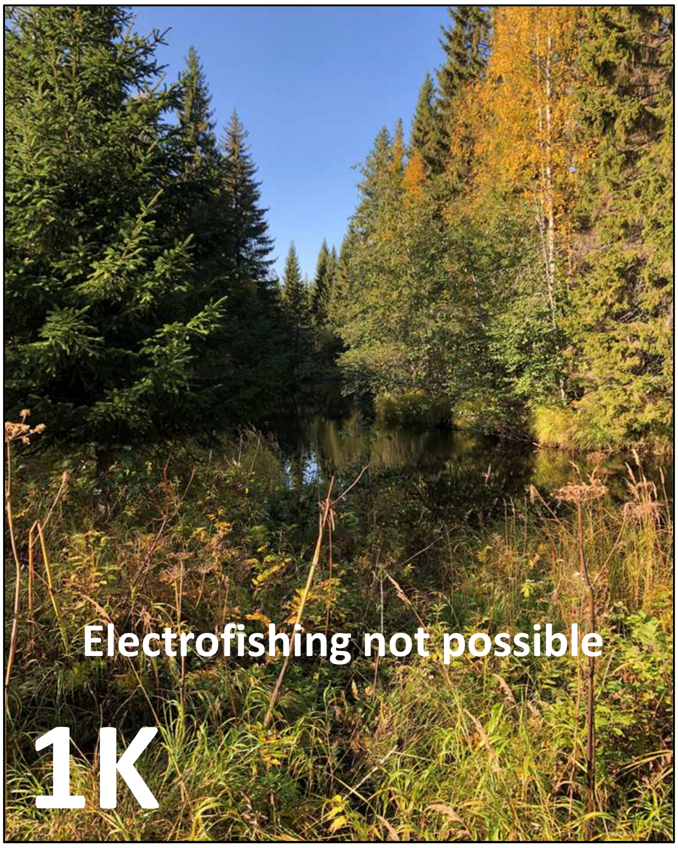

### Ume River: Stomdalsbäcken

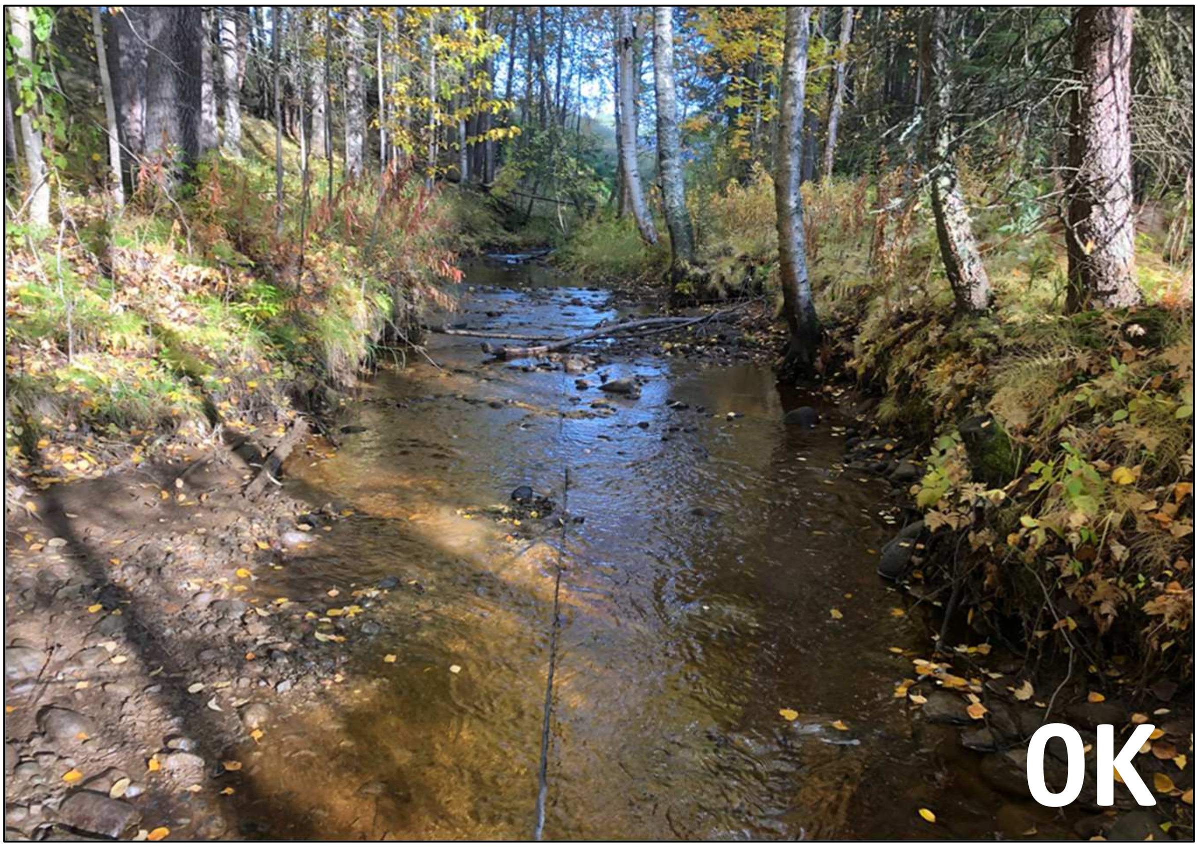


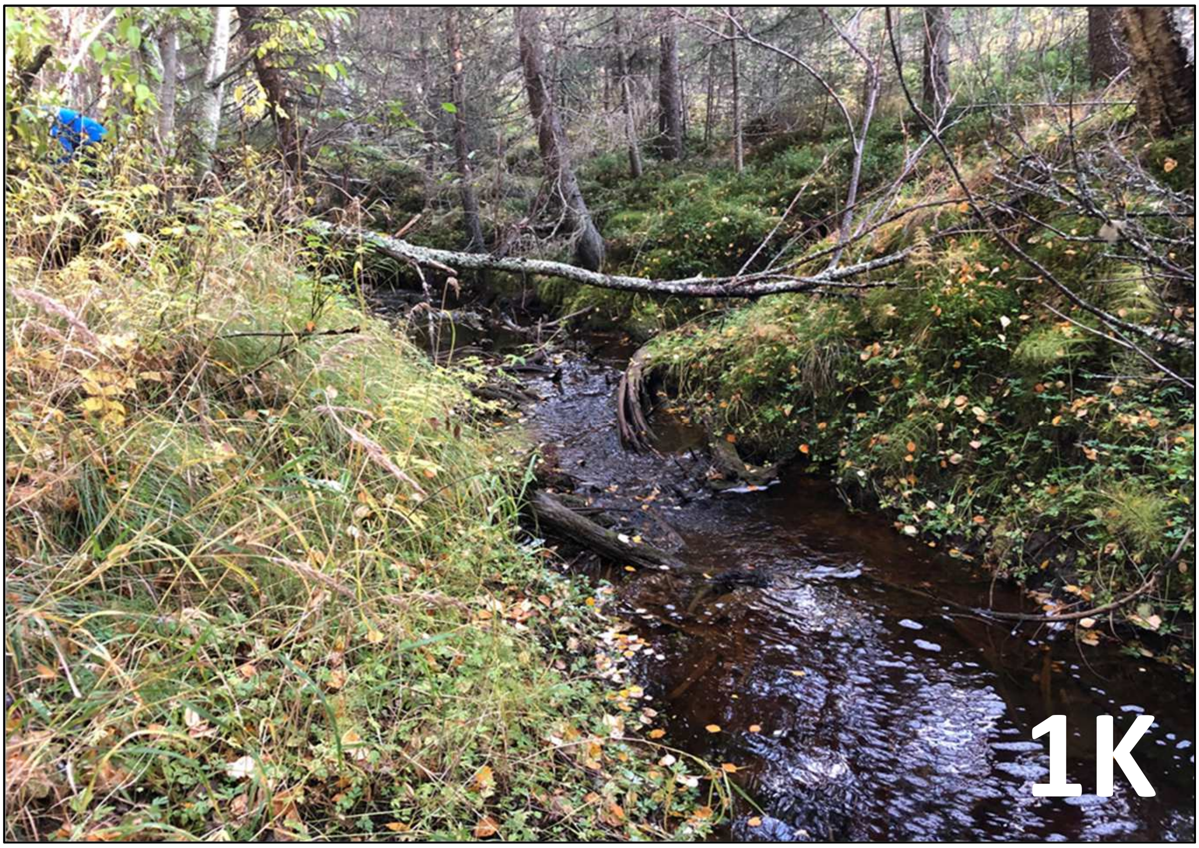

### Ume River: Vidbäcken

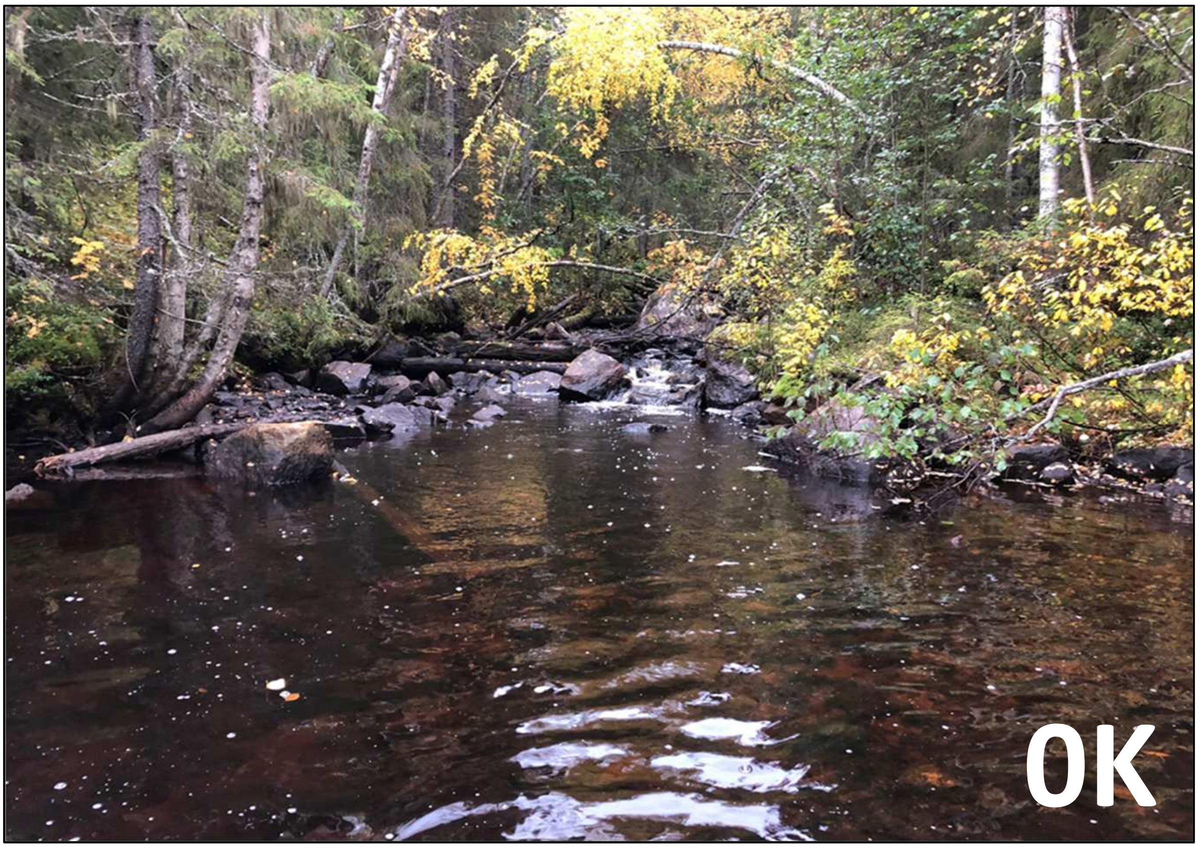


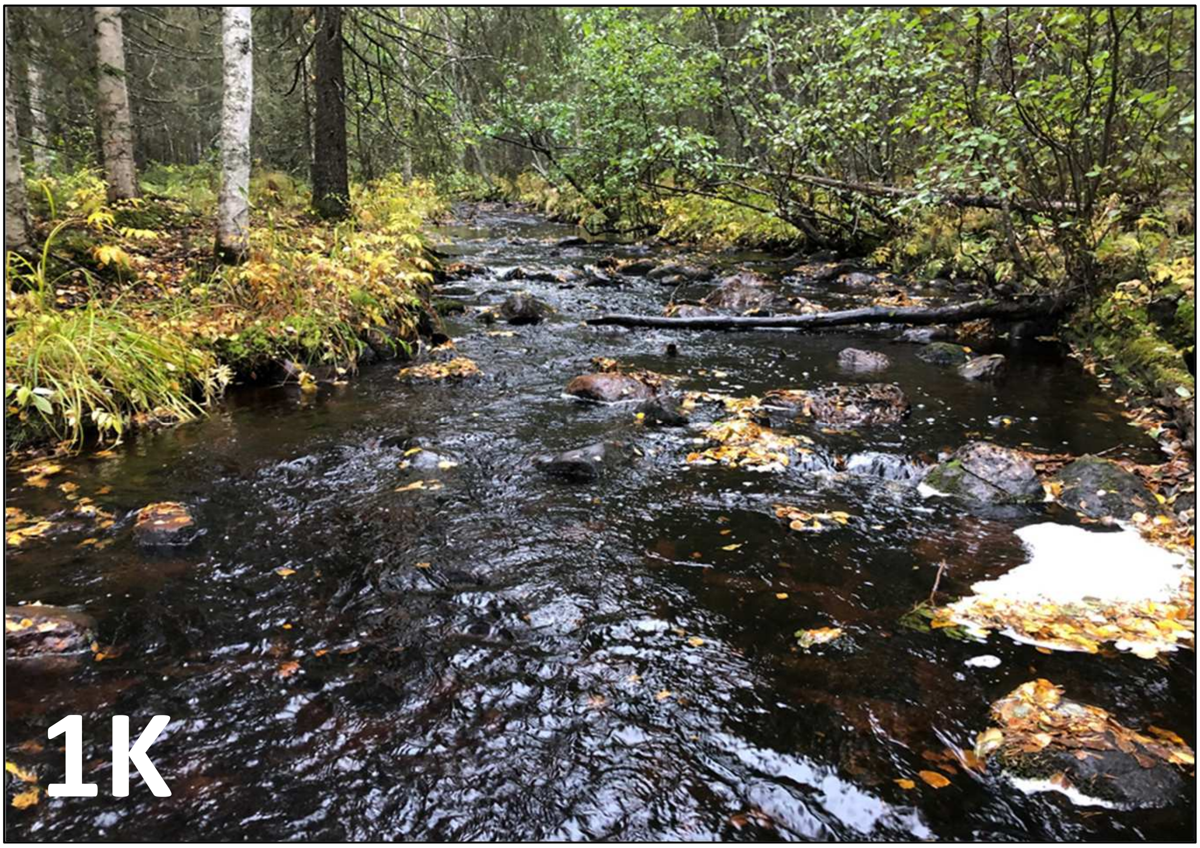

### Ume River: Kvarnbäcken.2

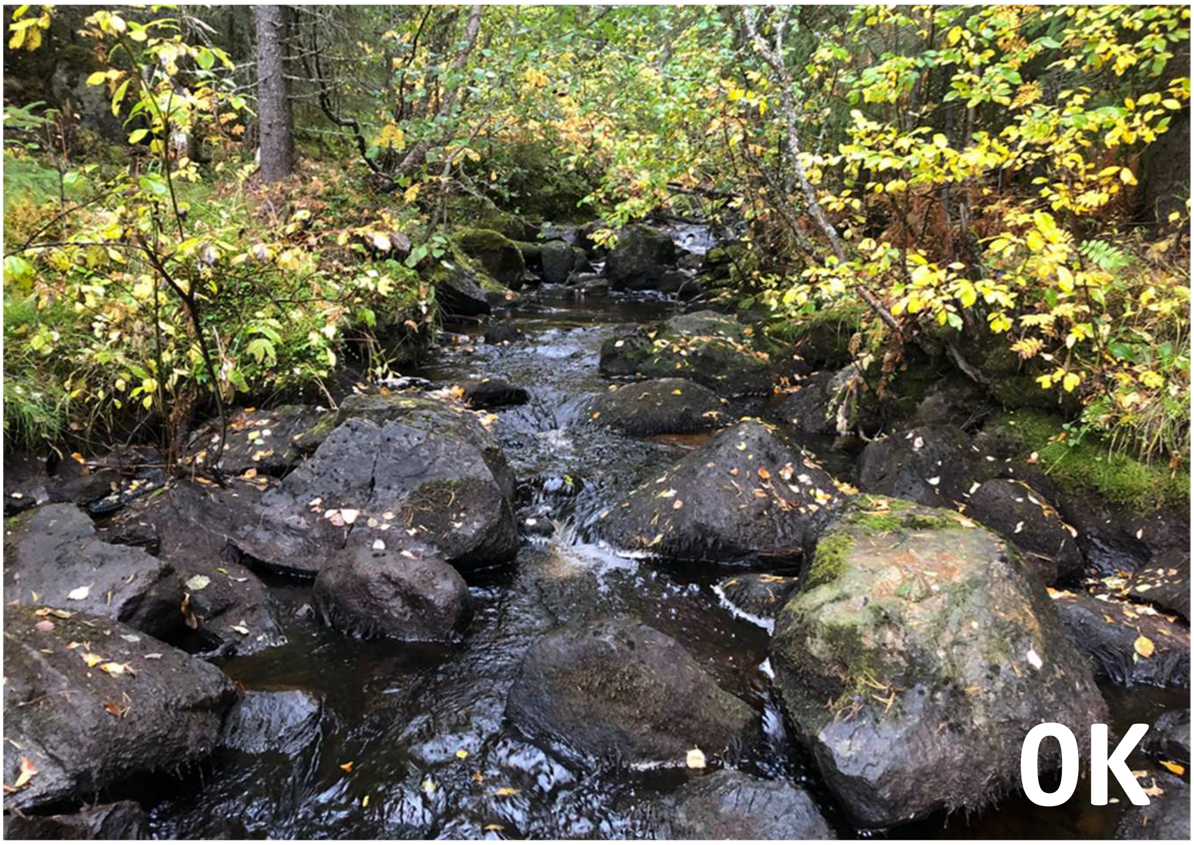


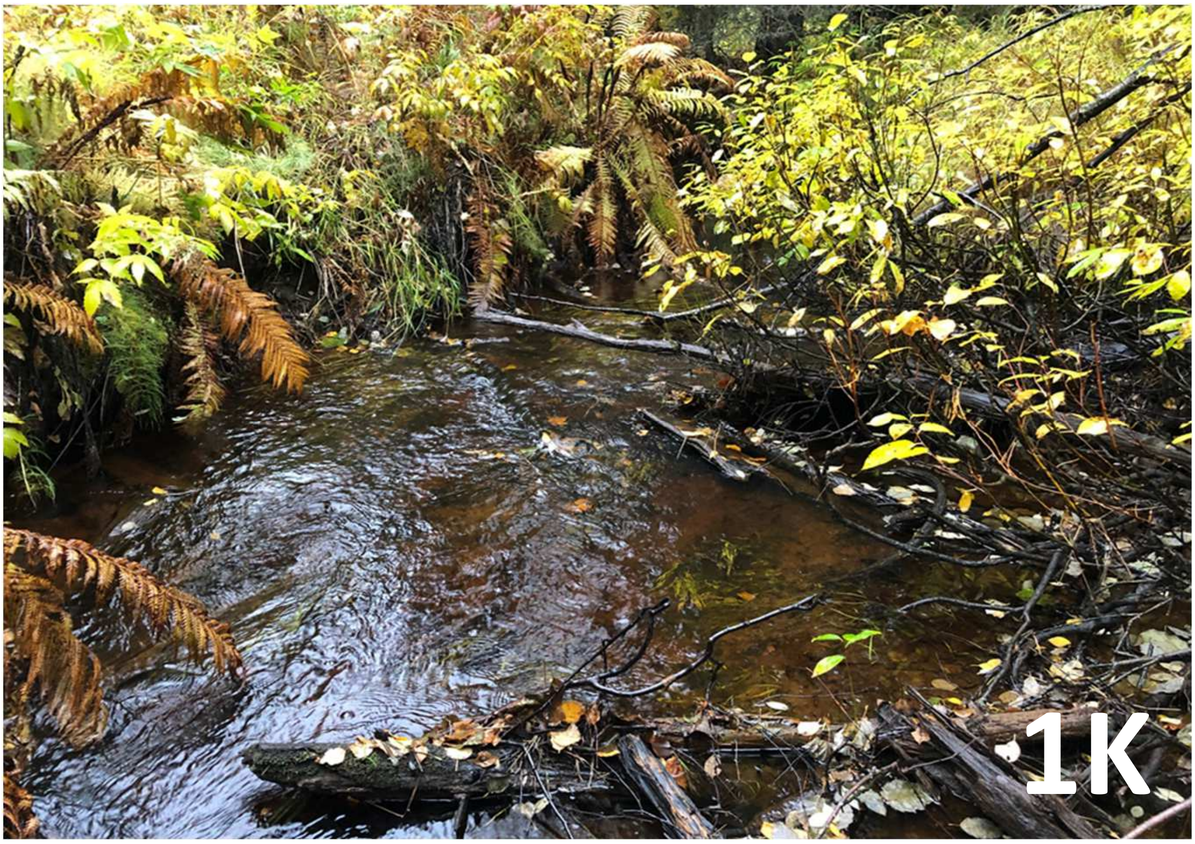

### Ume River: Byssjan

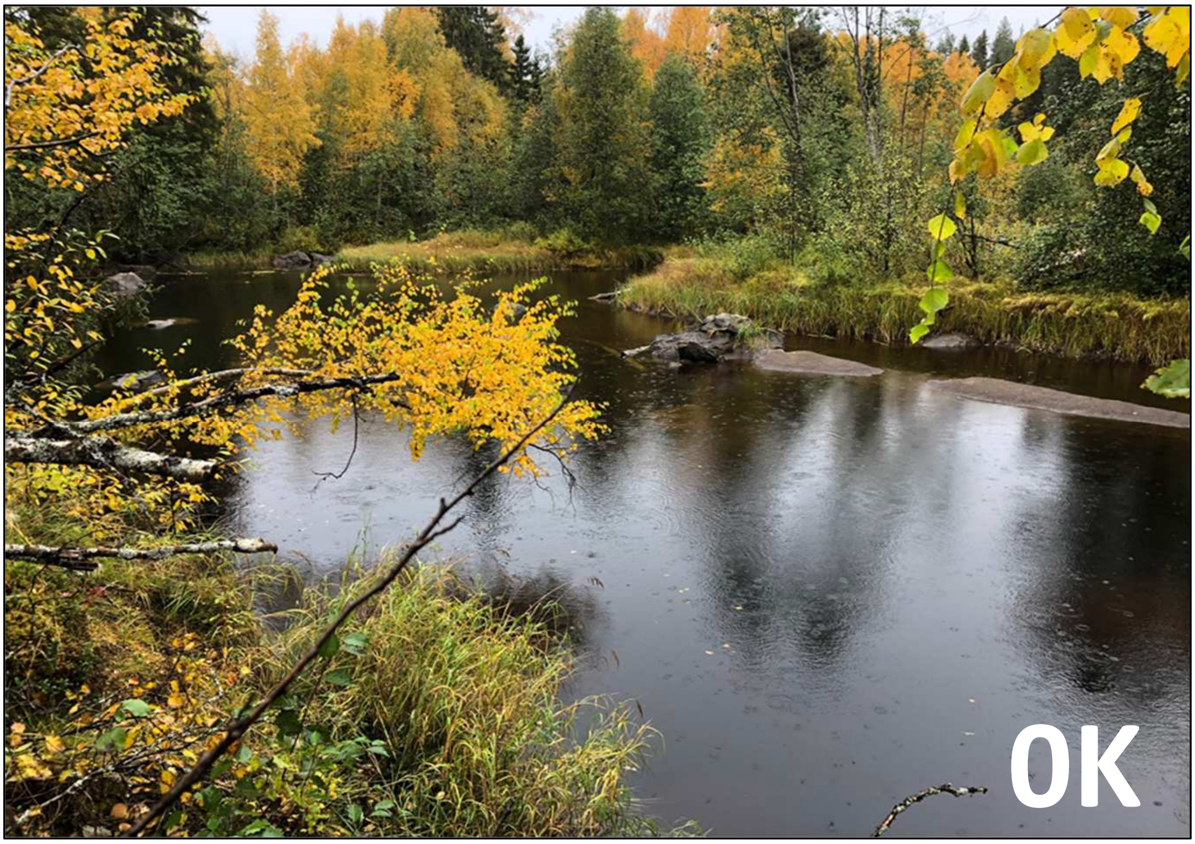


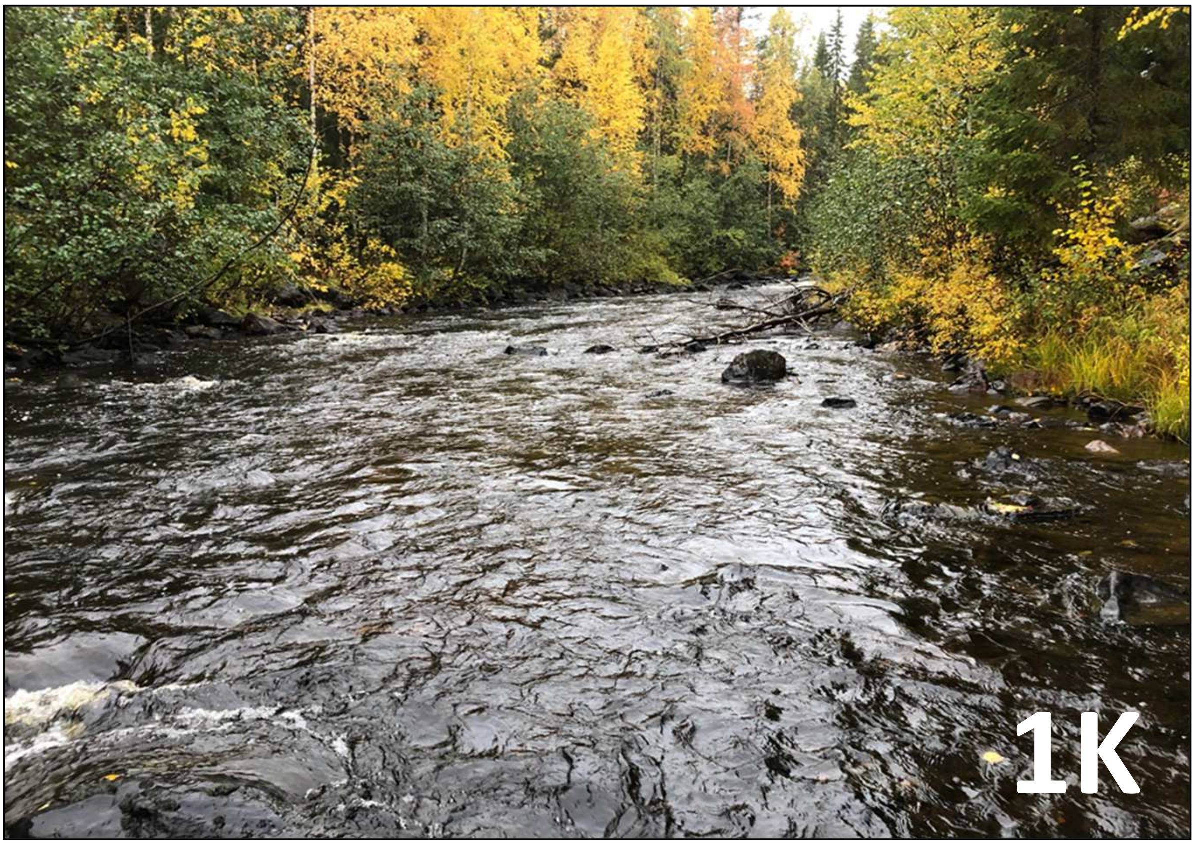

### Ume River: Nyraningsbäcken

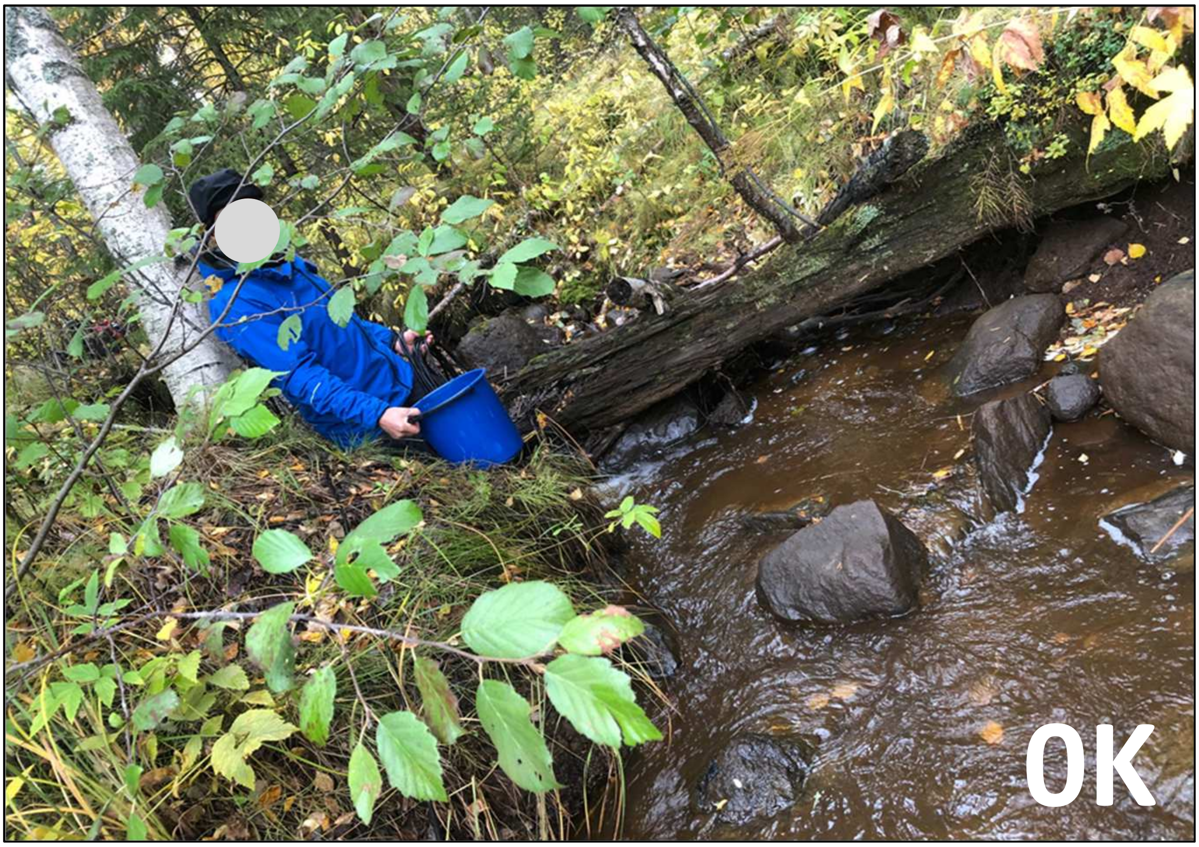


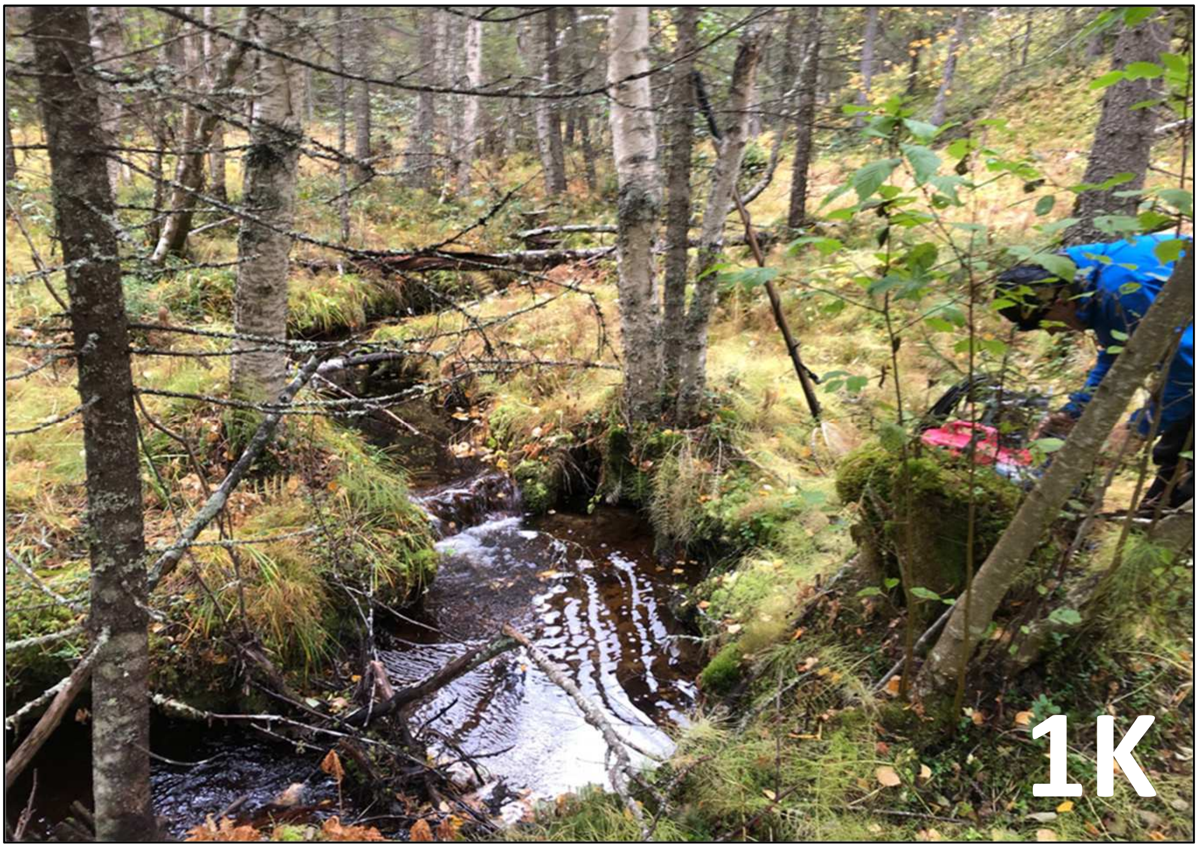

### Ume River: Illbäcken

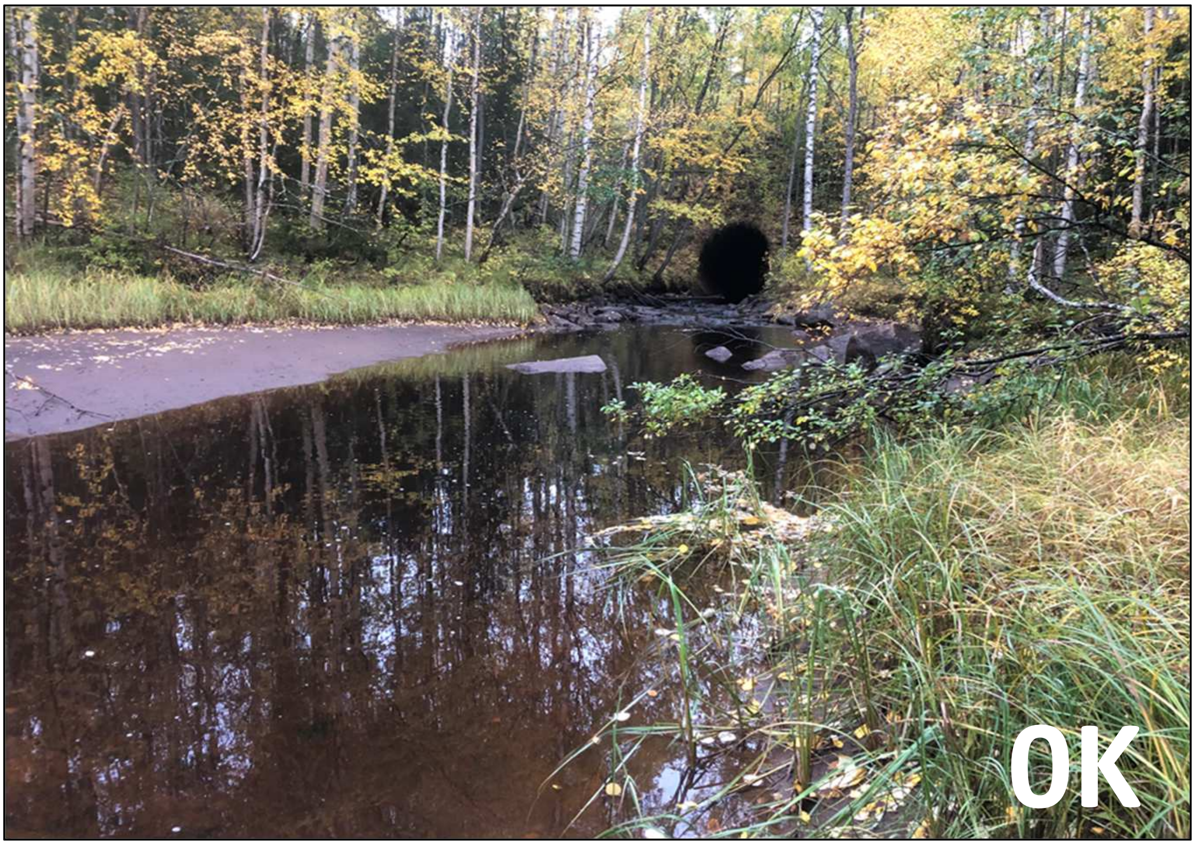


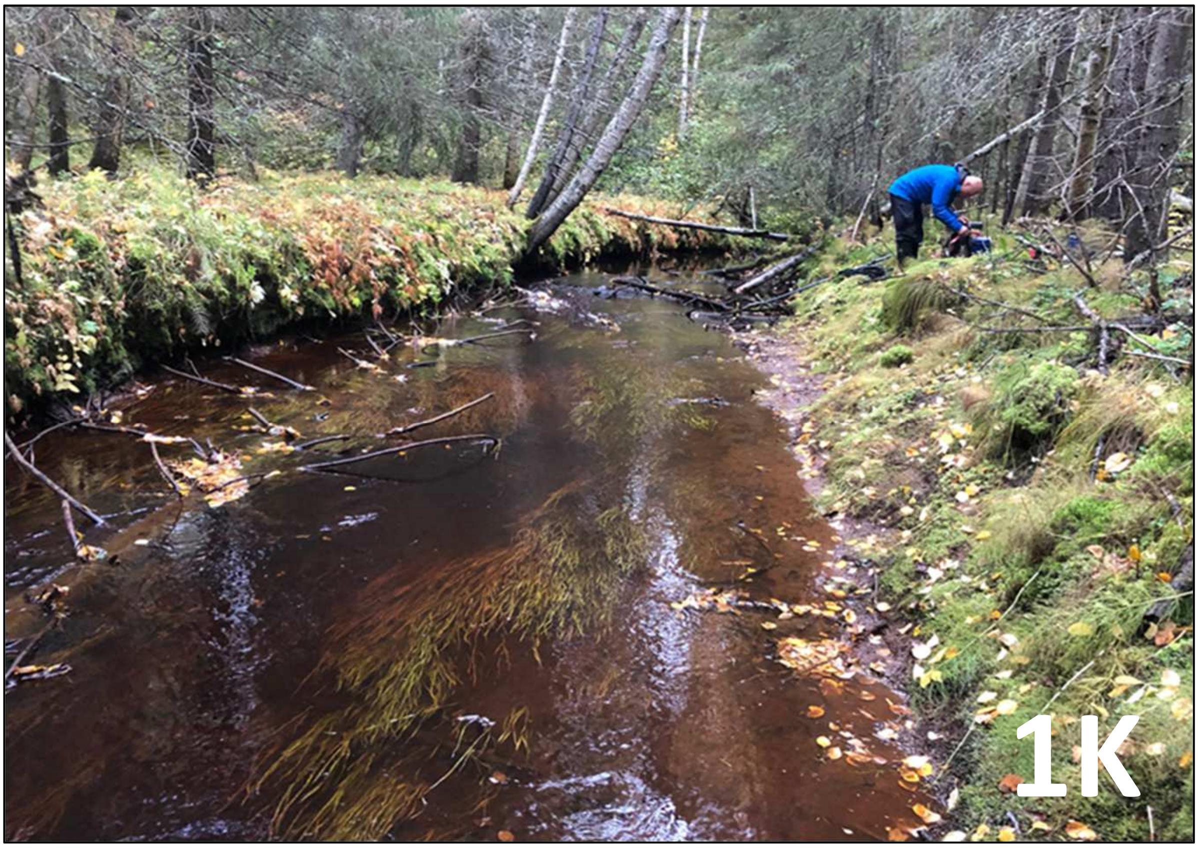

### Lule River: Norbäcken

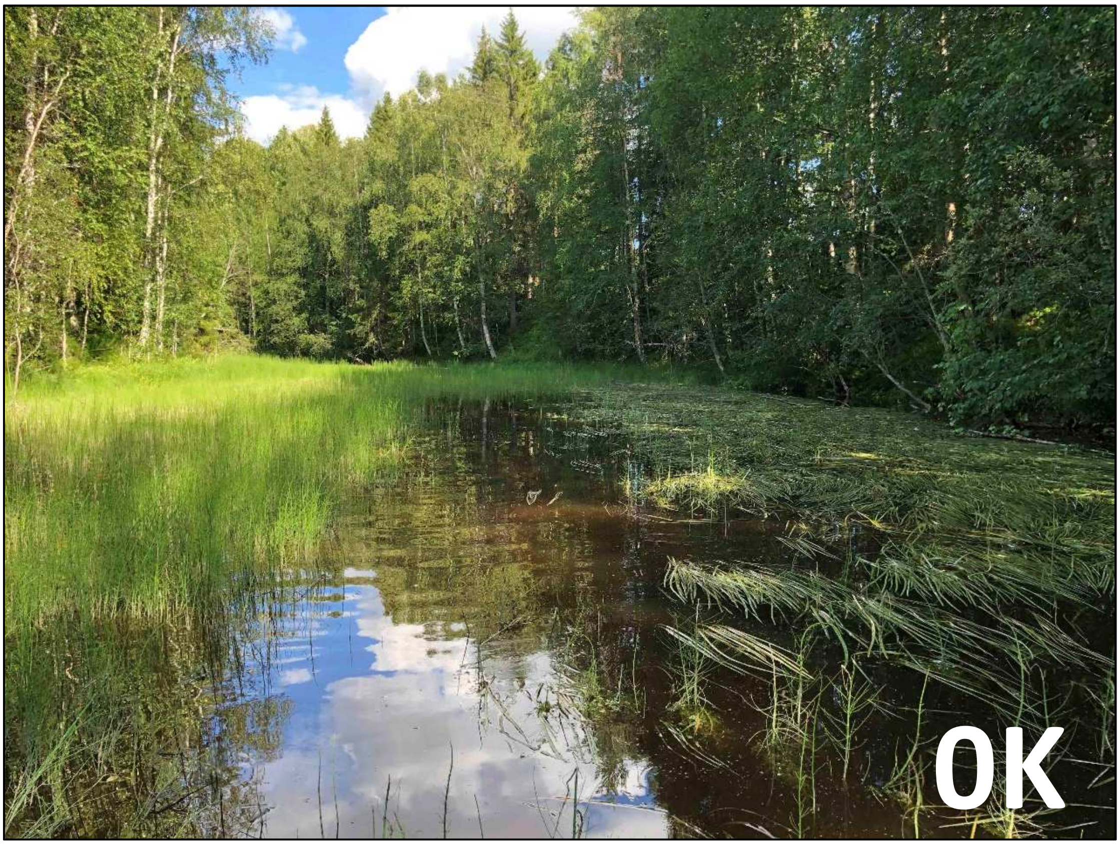


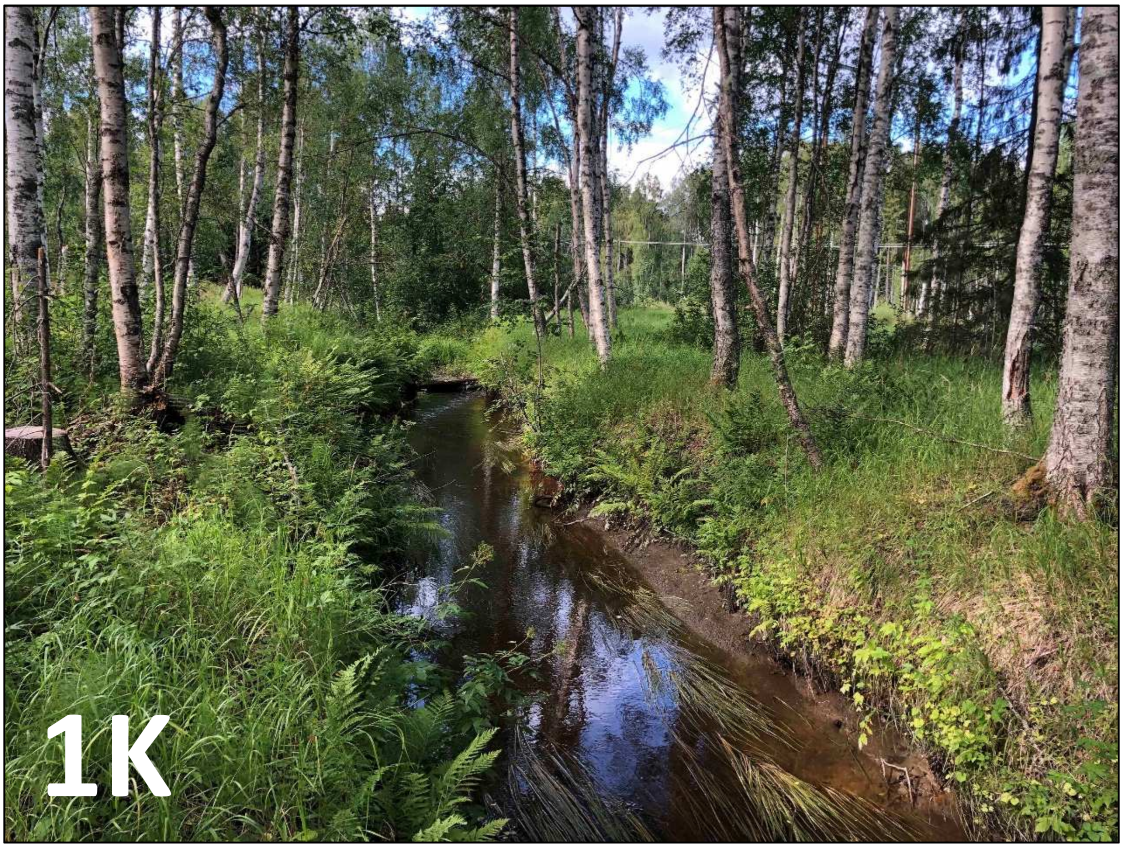

### Lule River: Degerbäcken

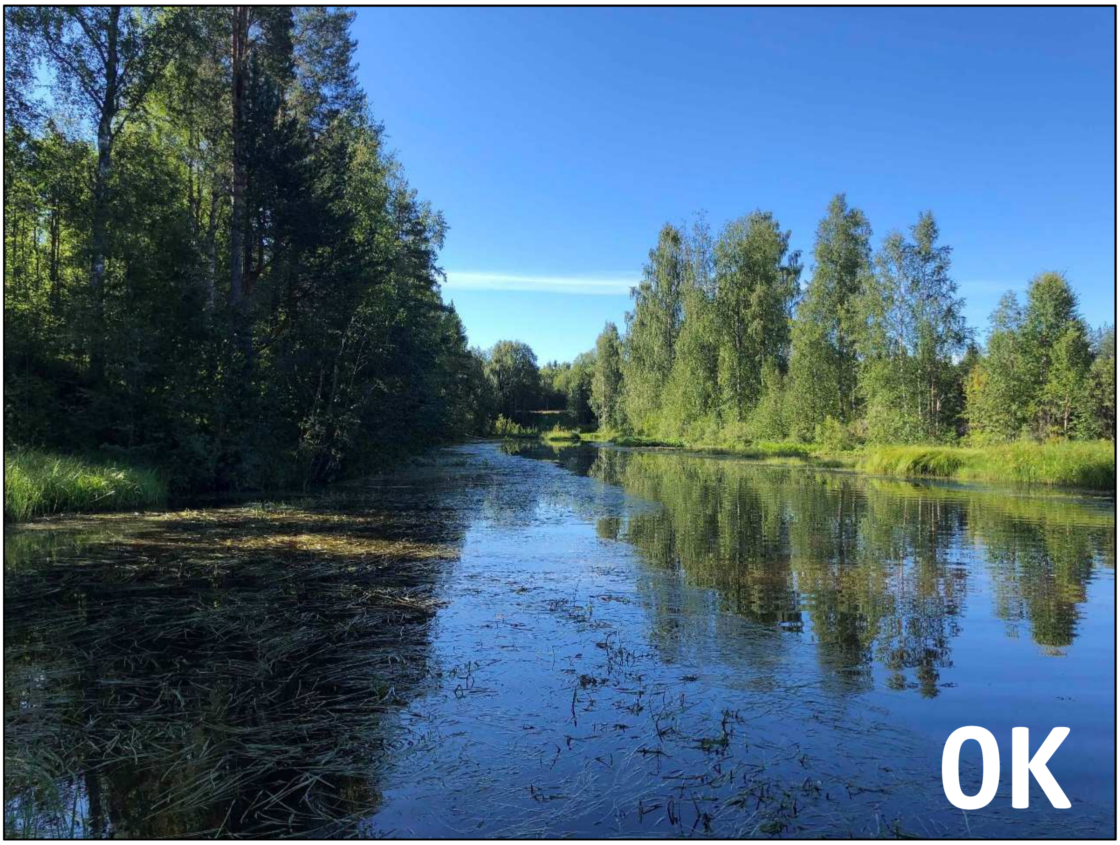


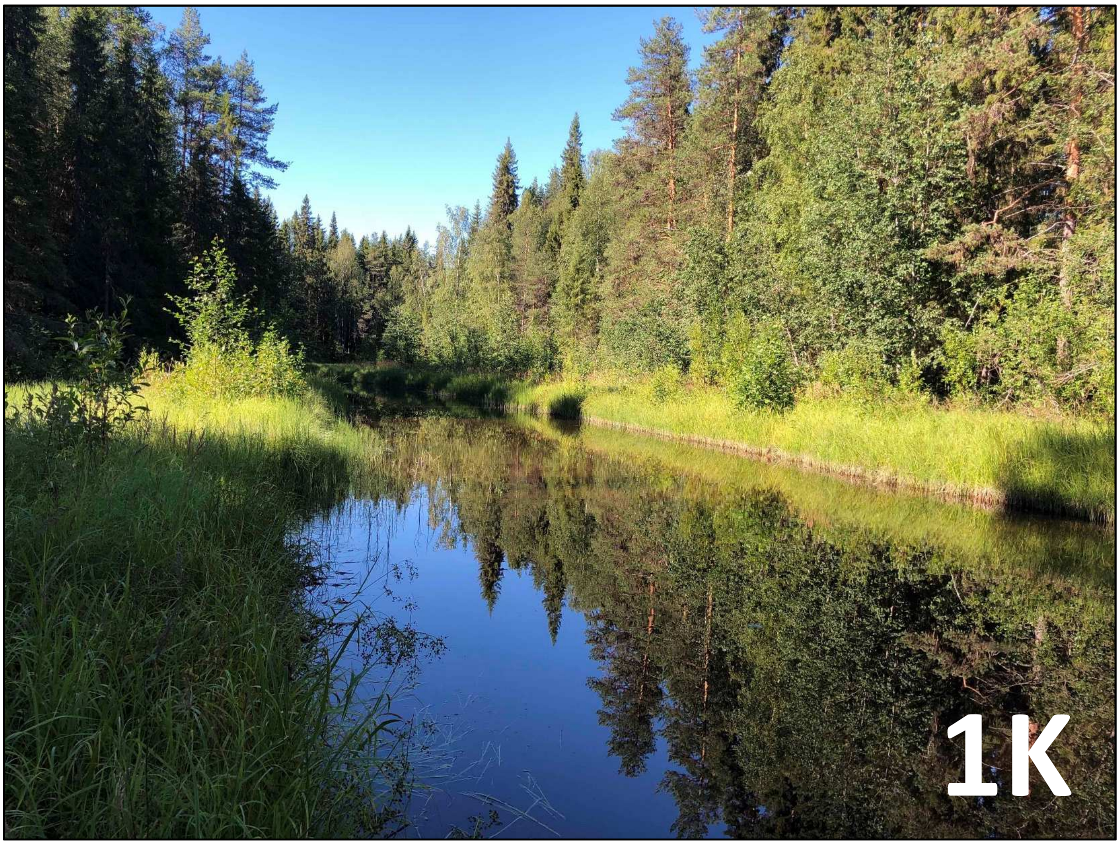

### Lule River: Kvarnbäcken.4

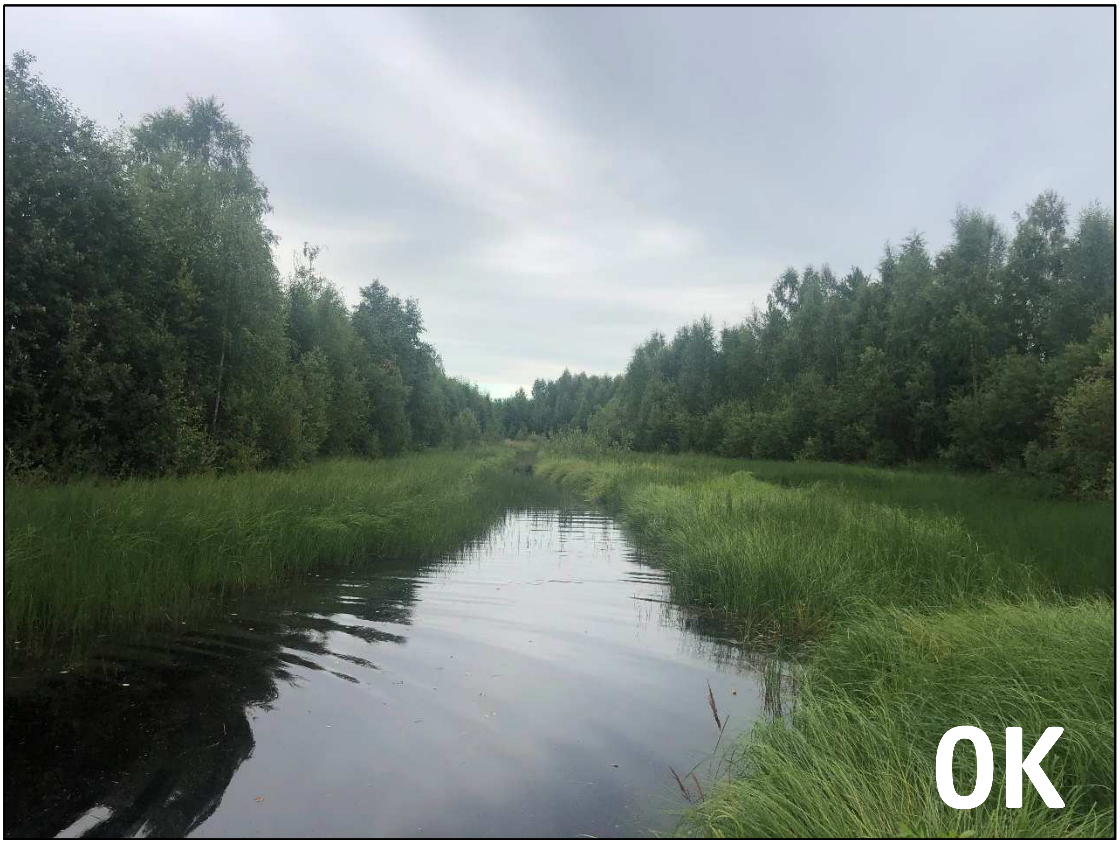


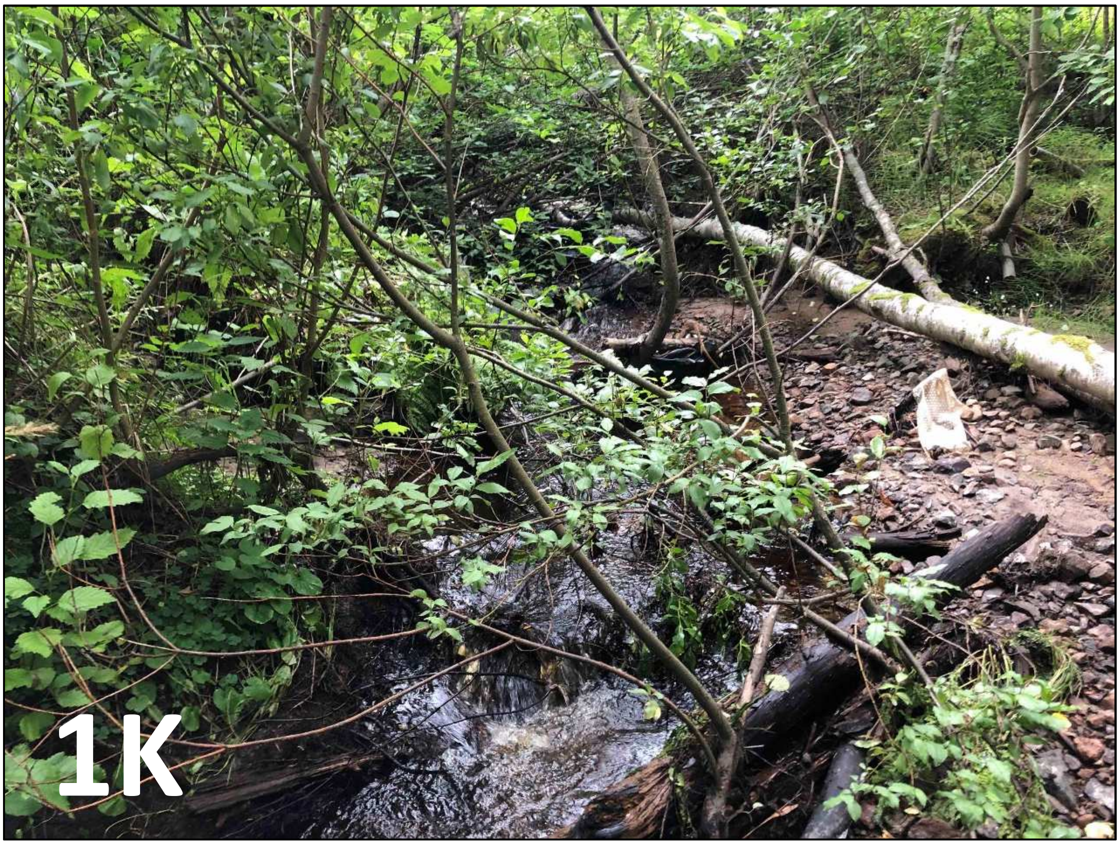

### Lule River: Bjurbäcken

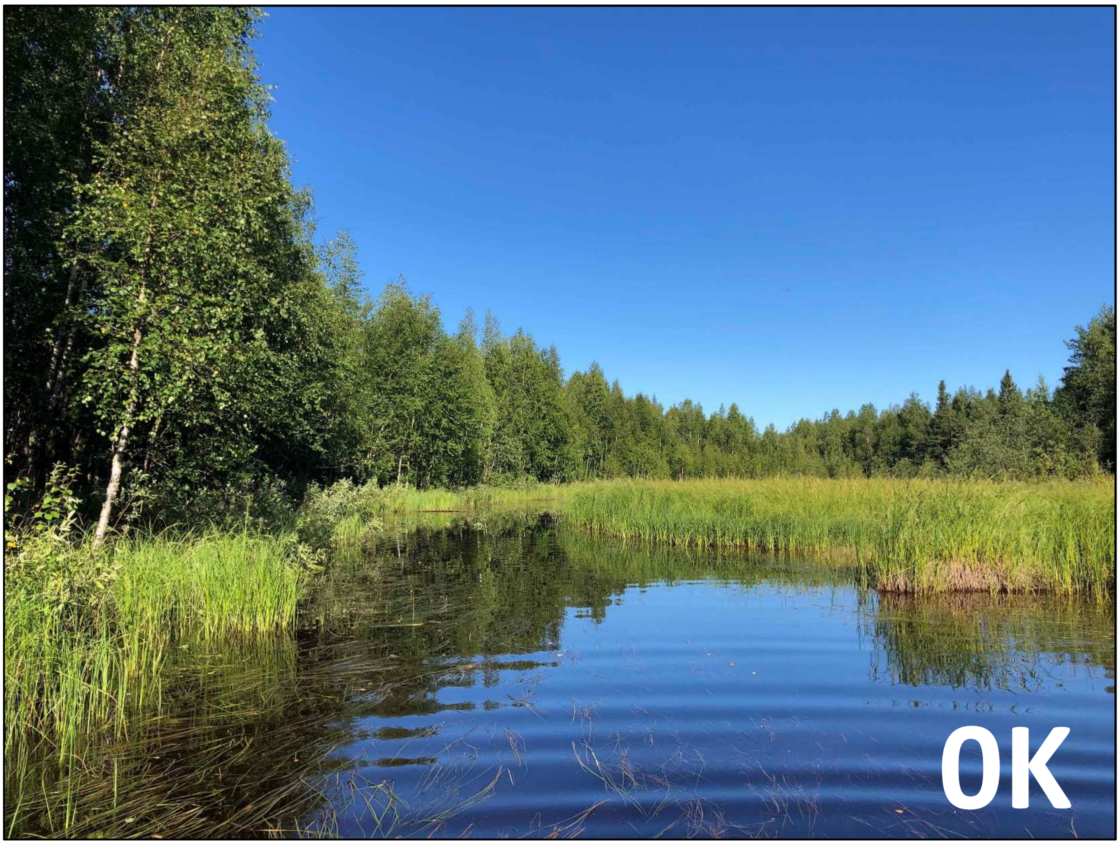


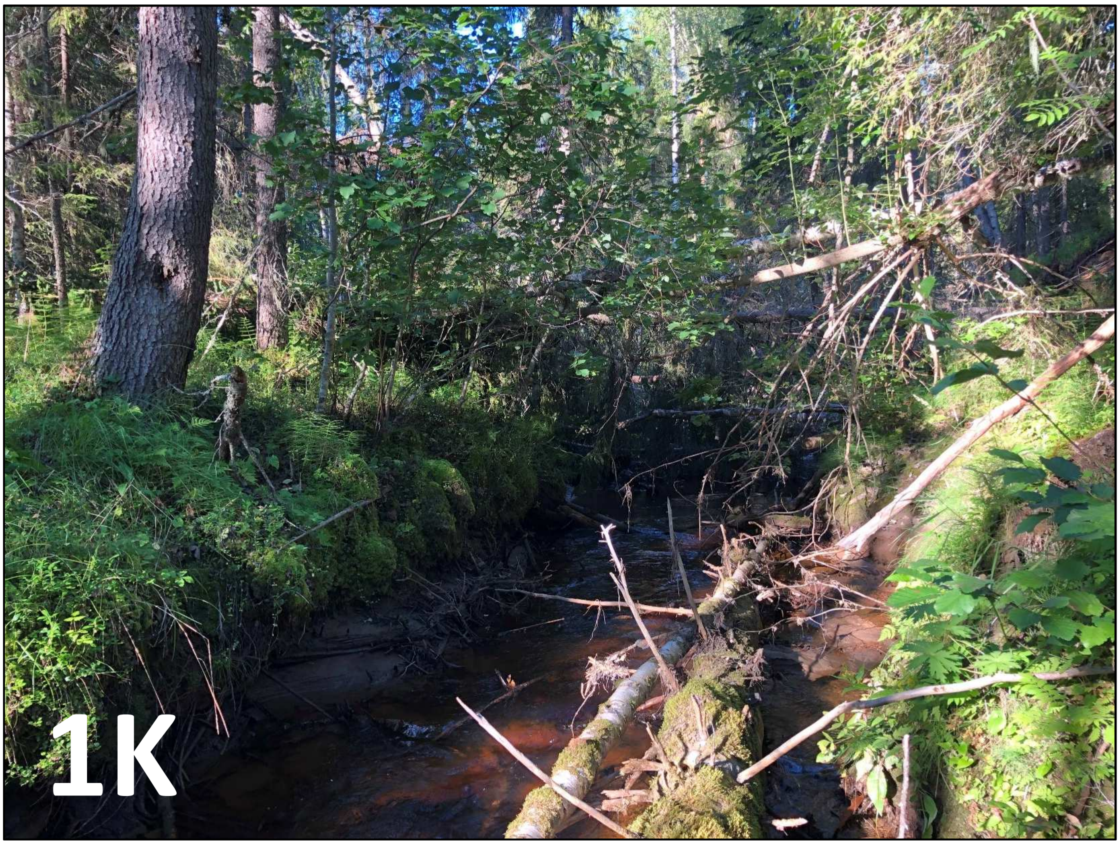

### Lule River: Forsträskbäcken

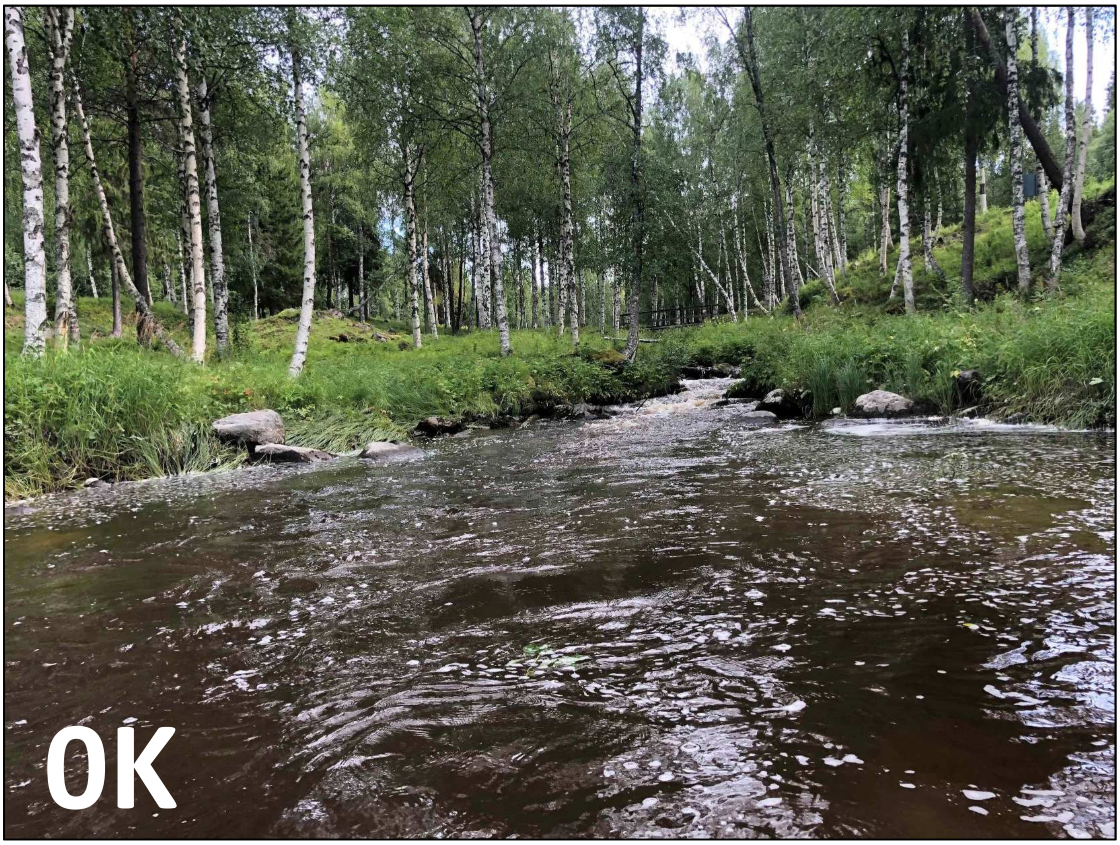


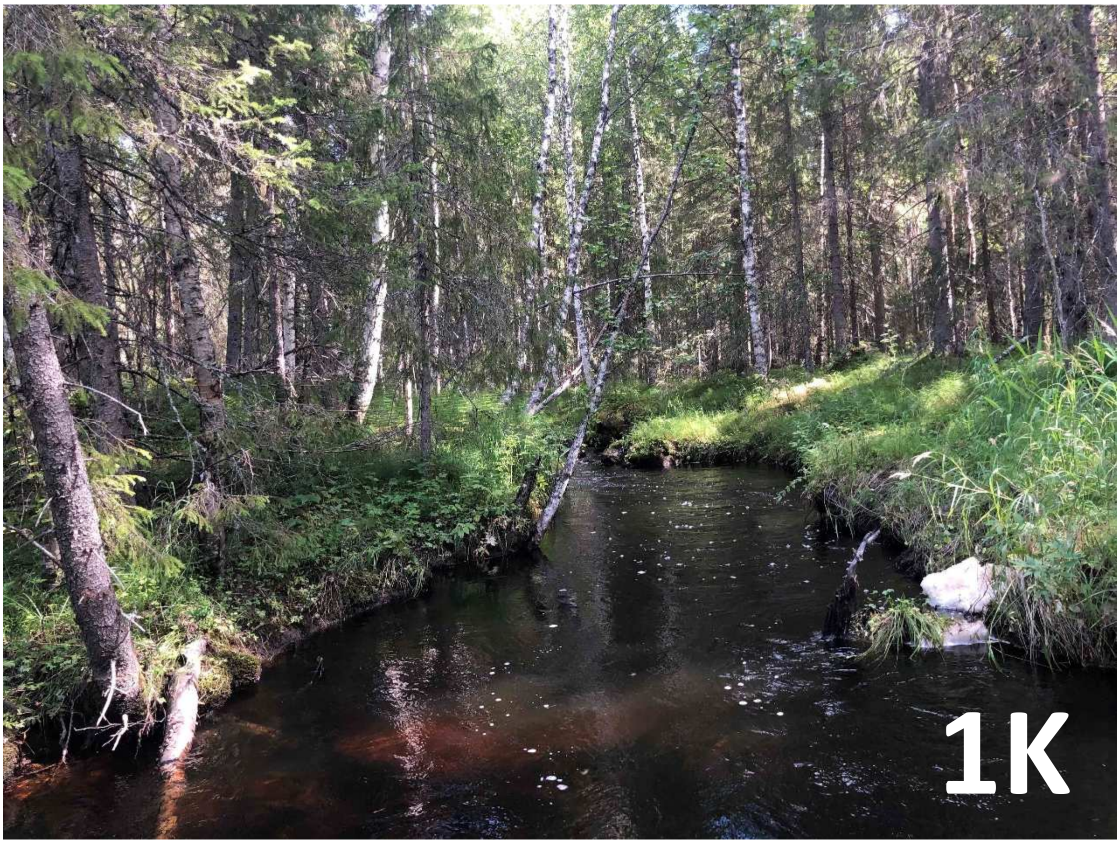

### Lule River: Görjeån

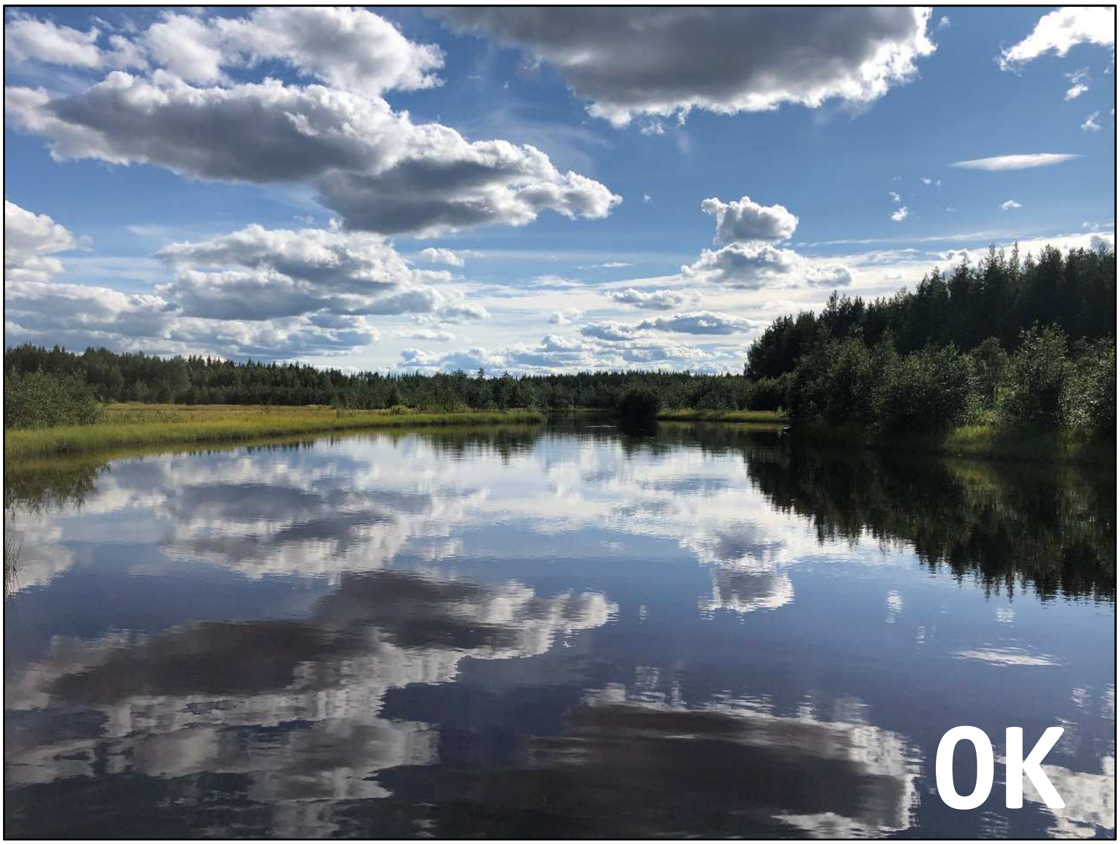


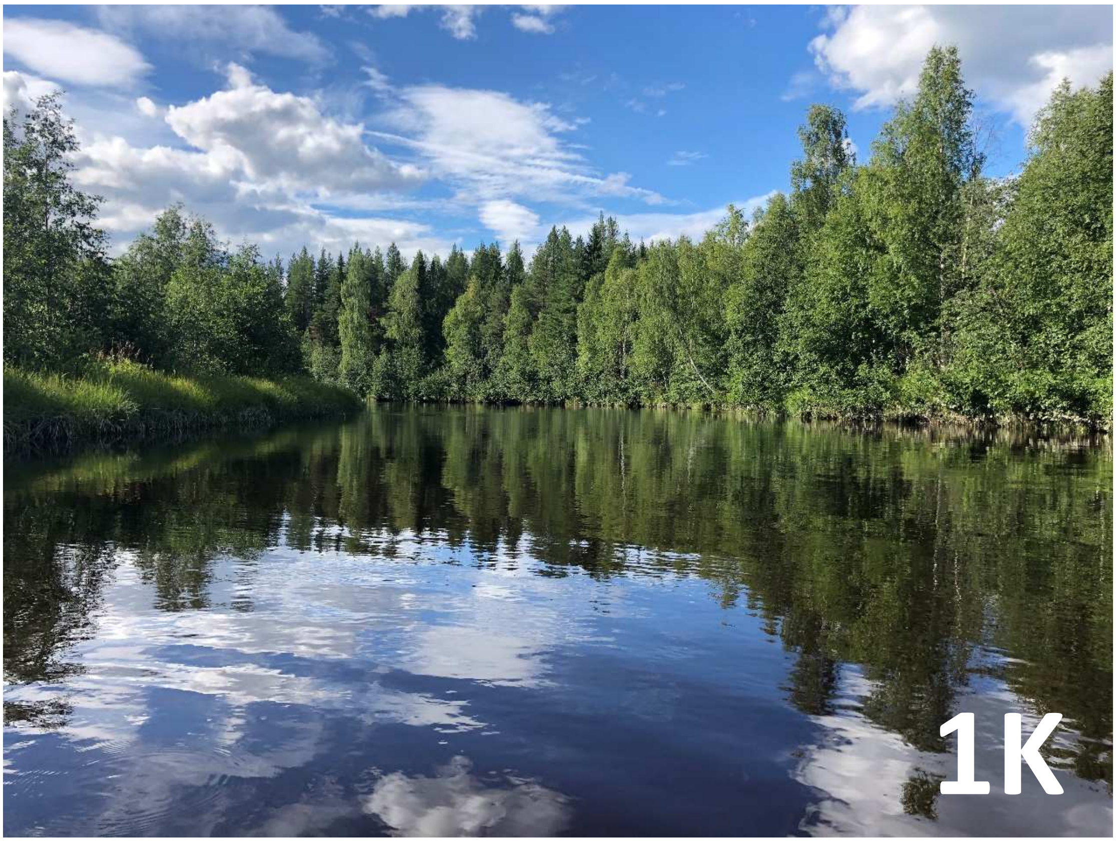

### Lule River: Lagnäsån

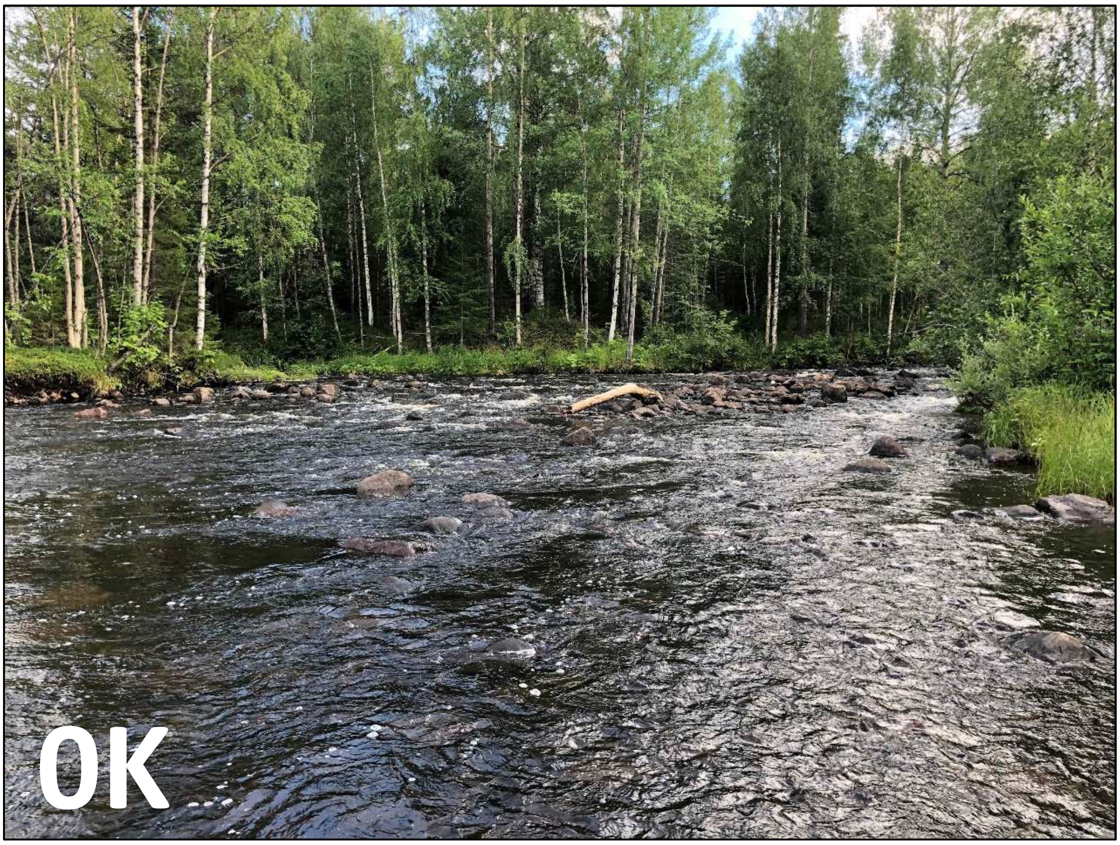


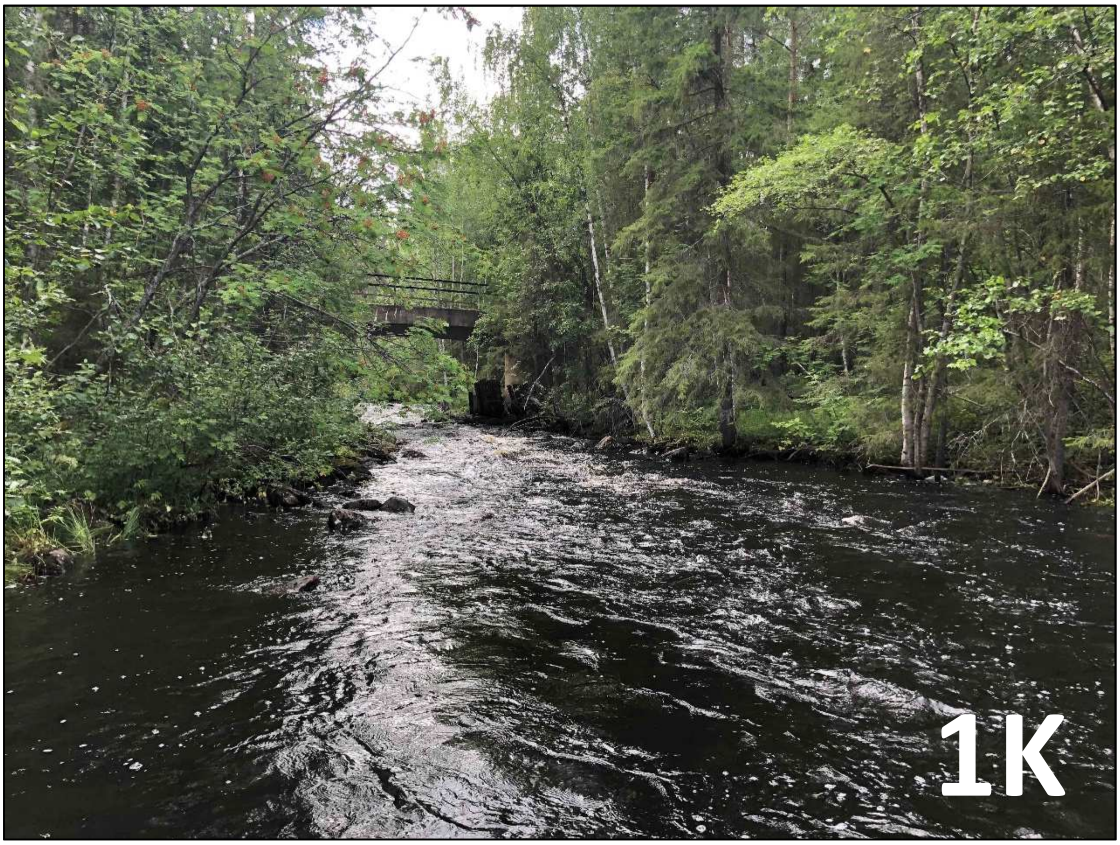

### Lule River: Kistabäcken

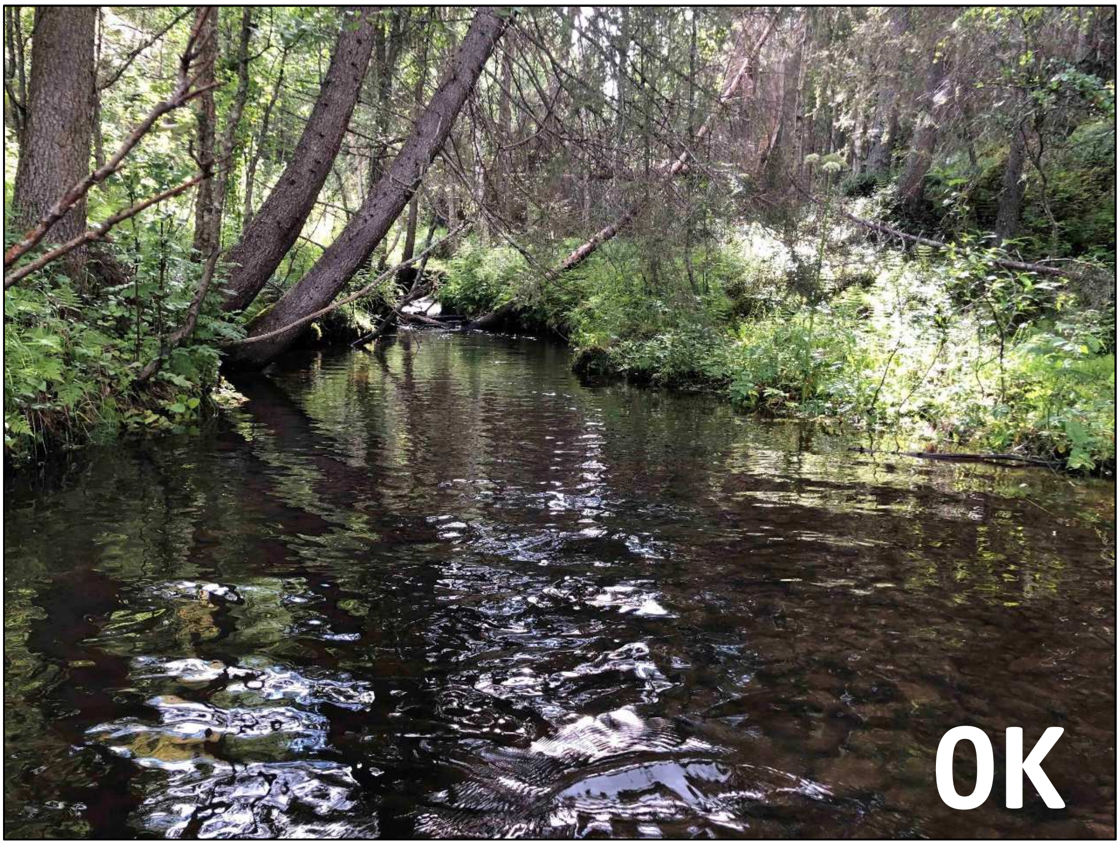


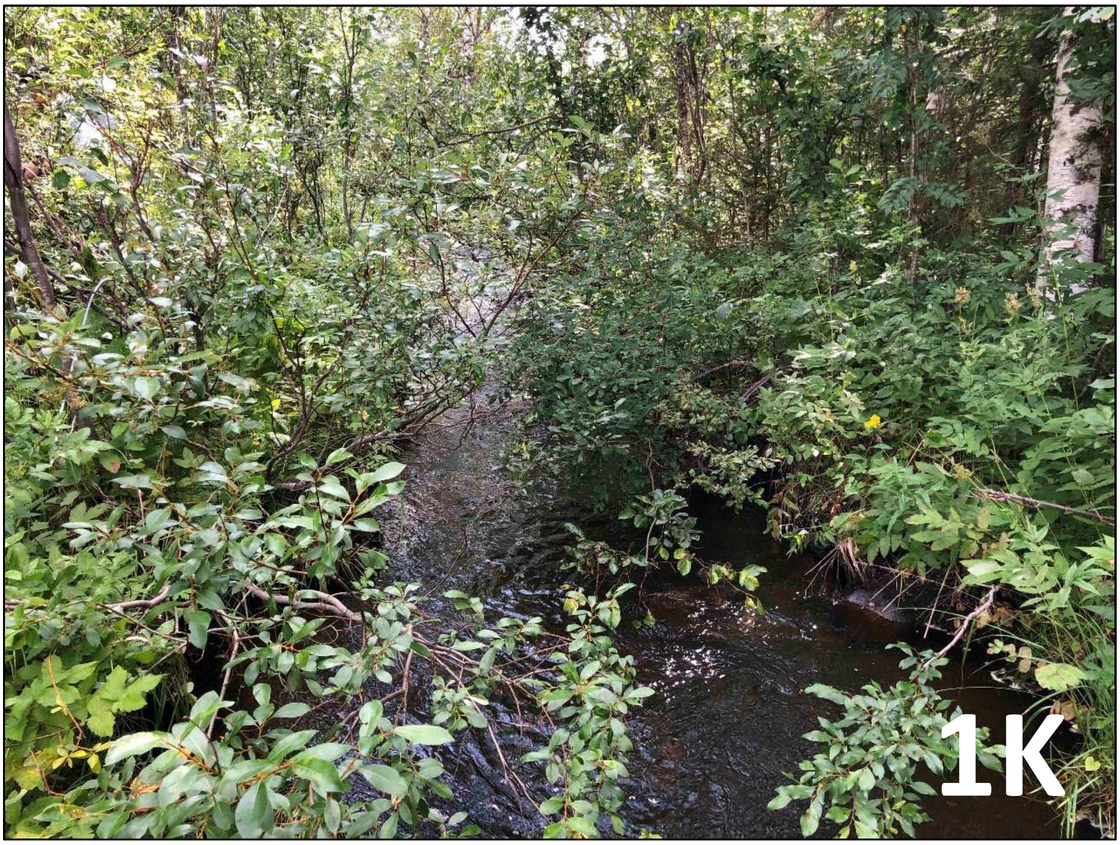

### Lule River: Andrensbäcken

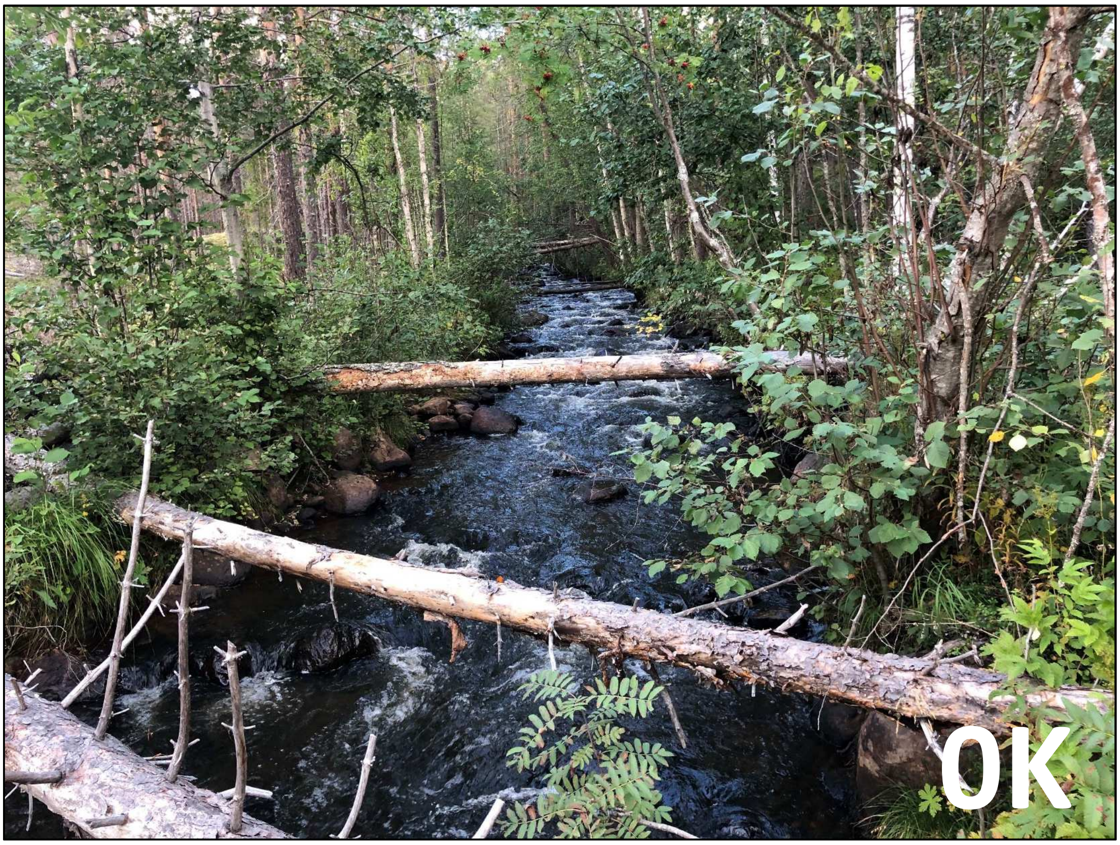


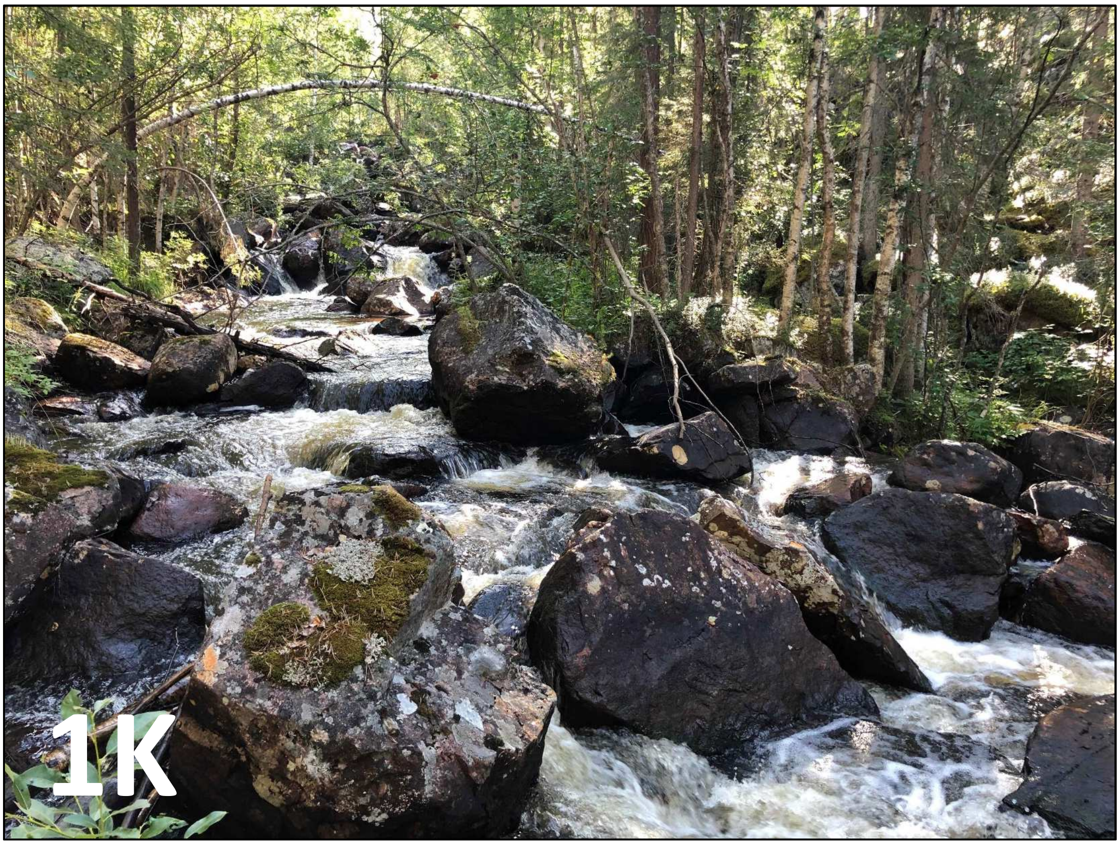

### Lule River: Kanibäcken

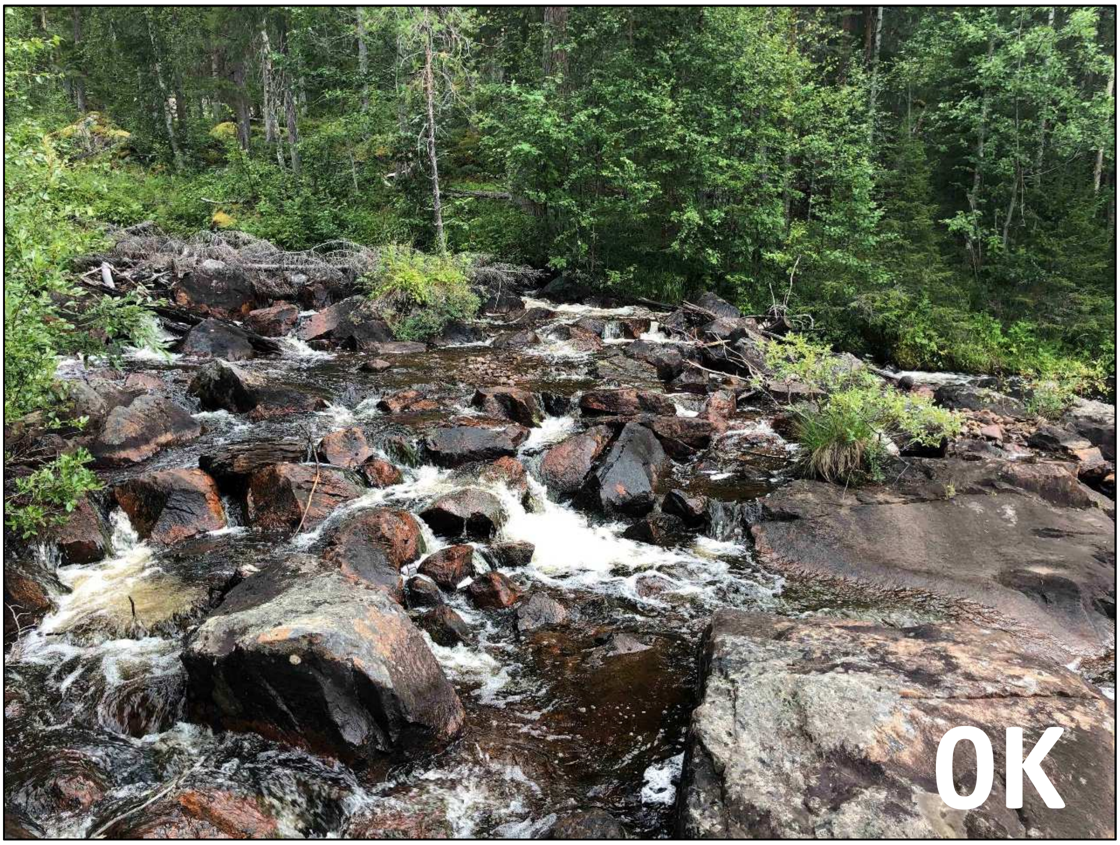


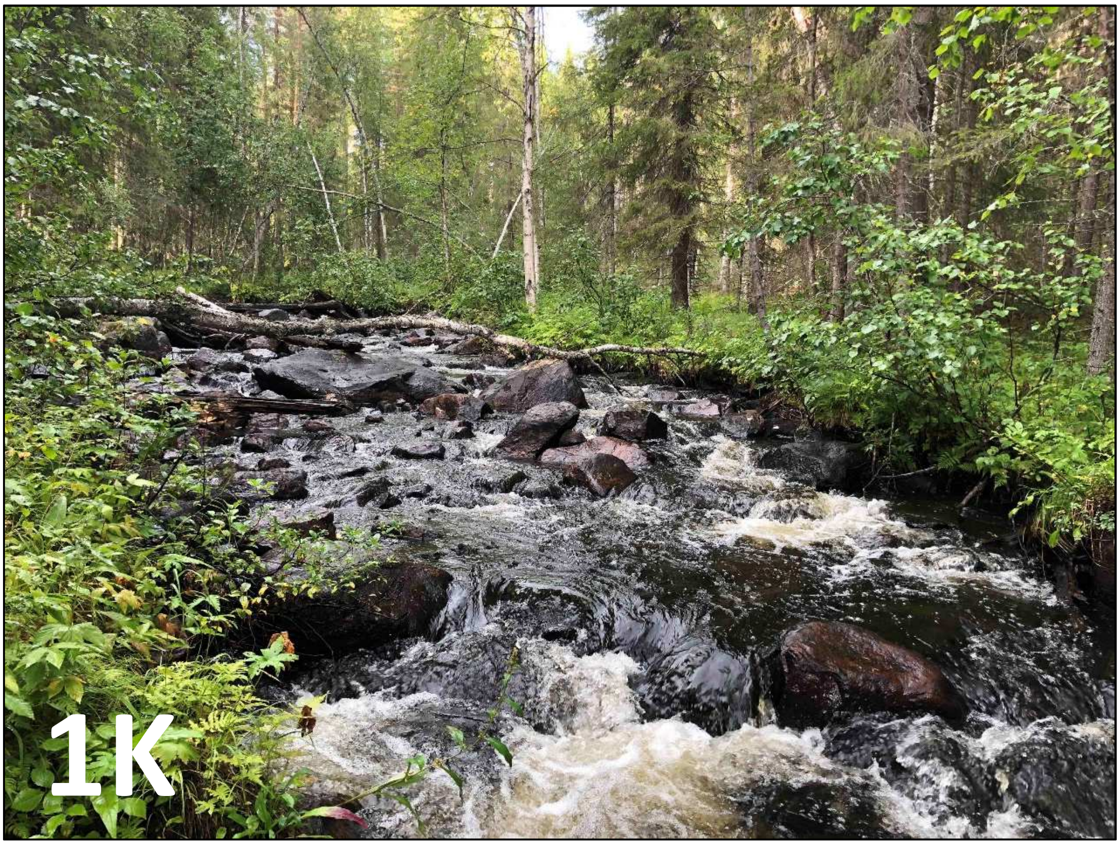
